# Breaking anterior-posterior symmetry in the moth fly *Clogmia albipunctata*

**DOI:** 10.1101/2025.01.13.632851

**Authors:** Ezra E. Amiri, Ayse Tenger-Trolander, Muzi Li, Alexander Thomas Julian, Koray Kasan, Sheri A. Sanders, Shelby Blythe, Urs Schmidt-Ott

## Abstract

Establishing the anterior-posterior (AP) body axis is a fundamental process during embryogenesis, and the fruit fly, *Drosophila melanogaster*, provides one of the best-known case studies. But for unknown reasons, different species of flies (Diptera) establish the AP axis through unrelated, structurally distinct anterior determinants (ADs). The AD of Drosophila, Bicoid (Bcd), initiates symmetry-breaking during nuclear cleavage cycles (NCs) when ubiquitous pioneer factors, such as Zelda (Zld), drive zygotic genome activation (ZGA) at the level chromatin accessibility by nucleosome depletion. While Bcd engages in a concentration-dependent competition with nucleosomes at the loci of a small set of transcription factor (TF) genes that are expressed in the anterior embryo, it remains unknown whether unrelated ADs of other fly species function in the same way and target homologous genes. We have examined the symmetry-breaking mechanism of a moth fly, *Clogmia albipunctata*, in which a maternally expressed transcript isoform of the pair-rule segmentation gene *odd-paired* serves as AD. We provide a *de novo* assembly and annotation of the Clogmia genome and describe how Clogmia’s orthologs of *zelda* (*Cal-zld*) and *odd-paired* (*Cal-opa*) affect chromatin accessibility and gene expression. Our results suggest direct roles of *Cal-zld* in opening and closing chromatin during nuclear cleavage cycles (NCs) and show that during the early phase of ZGA maternal *Cal-opa* activity promotes chromatin accessibility and anterior expression at Clogmia’s *homeobrain* and *sloppy-paired* loci. These genes are not known as key targets of Bcd but may serve a more widely conserved role in the initiation of anterior pattern formation given their early anterior expression and function in head development in insects. We conclude that the ADs of Drosophila and Clogmia differ in their target genes but share the mechanism of concentration-dependent nucleosome depletion.

## Introduction

The insect order of flies (Diptera) offers excellent opportunities for comparative study of developmental mechanisms, as many dipteran species can be reared cost-effectively and because the fruit fly, *Drosophila melanogaster*, provides a well-established model for functional comparisons at the molecular and cellular level. Dipteran embryos are morphologically similar across the clade (1, 2). However, the mechanisms of axis specification (3, 4), primordial germ cell specification (3), extraembryonic tissue specification (5, 6), morphogenesis (5, 7–10), and sex determination (11–14) differ between dipteran species. Explaining such unexpected plasticity in developmental gene networks requires functional comparisons, which can reveal underlying general principles and mechanisms of evolutionary change. Anterior-posterior (**AP**) axis specification, a long-standing model for pattern formation and gene regulation in Drosophila (15–23), is particularly suitable for comparative functional studies because the syncytial nature of early dipteran embryos facilitates visualization and perturbation of gene activity in non-traditional model organisms. Many dipterans establish the embryo’s head-to-tail polarity through transcription factors (**TF**s) that diffuse from maternally localized mRNA and form AP concentration gradients that regulate gene expression in a dose-dependent manner (24, 25). These anterior determinants (**AD**s) prevent the formation of bicaudal embryos that lack head and thorax (double abdomen). Irrespective of this fundamental role, unrelated ADs have been found in different fly species. While the AD of Drosophila and many other cyclorrhaphan fly species is encoded by a homeobox gene, *bicoid* (26–30), diverse zinc finger genes take on this role in lower dipterans, including *pangolin* (dipteran ortholog *Tcf* gene family) in anopheline mosquitoes and crane flies, *cucoid* in culicine mosquitoes, *panish* in certain harlequin flies (Chironomini), and *odd-paired* (dipteran ortholog of *Zic* gene family) in moth flies (Psychodidae) (3, 4). The phylogenetic occurrence of these ADs indicates that they have been frequently replaced during the dipteran radiation and that only a small fraction of dipteran ADs has been identified to date. Moreover, the AD proteins identified so far differ widely in their structure and DNA-binding domains. Bicoid (**Bcd**) binds DNA via its homeodomain, Pangolin (**Pan**) through HMG and C-clamp domains, Panish through a C-clamp domain, and Odd-paired (**Opa**) and Cucoid through distinct C2H2 zinc finger domains (31–36). This molecular diversity prompted us to ask whether the mechanisms of dipteran ADs and their target genes differ between species or whether they are conserved due to contextual or downstream network constraints.

We have addressed these questions by analyzing the mechanism of action of the AD of the moth fly *Clogmia albipunctata*, a C2H2 zinc finger protein encoded by the *zic* gene family member *odd-paired* (3) and comparing it to the AD of *Drosophila melanogaster*, a homeodomain protein encoded by *bicoid* (26), which is missing in Clogmia and other midges. The embryos of Drosophila and Clogmia undergo 13 nuclear division cycles (**NC**) before they cellularize and initiate gastrulation (37, 38). By the end of NC9 (Clogmia) or the beginning of NC10 (Drosophila), most nuclei of the embryo have arranged in a monolayer between the egg membrane and the yolk. This ‘syncytial blastoderm’ undergoes four additional nuclear cleavage cycles before it cellularizes during NC14, thereby ending the syncytial phase of embryogenesis. During this period, a segmental body plan has been established by a regulatory network of transcription factor encoding genes, known as gap and pair-rule genes (19, 21, 39, 40). In Drosophila, only a few hundred genes are expressed before the midblastula transition (NC14) (41–43), when mitotic cycles become desynchronized and the cell cycle lengthens partly due to the addition of gap phases, as the embryo transition from maternal to zygotic control of development (44). Genes expressed during NC8-NC13 include regulators of the cell cycle, sex determination, dosage compensation, and early pattern formation (45–47). Widespread zygotic genome activation (**ZGA**), however, is delayed until NC14 due to constraints on transcription imposed by rapid cell cycling and the requirement of global changes in chromatin accessibility (47–49). After each DNA replication cycle, the formation of nucleosomes reduces chromatin accessibility. This barrier is overcome by a special class of sequence-specific TFs capable of binding nucleosome occupied DNA and destabilizing nucleosomes (50, 51). The major driver of chromatin accessibility during Drosophila’s nuclear cleavage cycles is Zelda (**Zld**), a ubiquitously expressed zinc finger protein (46, 52–55). Zld’s activity is required throughout ZGA and enables the binding of other TFs including those involved in embryonic patterning (53, 56–60). Two additional pioneer TFs, GAGA Factor (GAF) and CLAMP, act independently and cooperatively with Zld to establish open chromatin and drive zygotic gene expression once major ZGA is underway (61–64).

Bcd’s role in AP patterning becomes indispensable after blastoderm formation at the beginning of NC10 (65). During early blastoderm stages, chromatin accessibility measured by ATAC-seq (assay for transposase-accessible chromatin with sequencing) is largely homogeneous across the embryo. Indeed, Zld establishes chromatin accessibility ubiquitoulsy in the embryo at many sites that are targeted by Bcd (57, 59, 66–72). However, a small cohort of early segmentation gene enhancers that drive Bcd-dependent anterior expression exhibits anteriorly restricted accessibility (33, 73–79). The underlying mechanism has been inferred by measuring accessibility and activity of the Bcd-dependent proximal P2 enhancer element of *hunchback*, an early target of Bcd, under experimentally modulated Bcd input and nucleosome stability at the enhancer (80). Bcd engages in a concentration-dependent competition with nucleosomes to ‘open’ otherwise inaccessible regulatory regions. Bcd activity thereby effectively breaking symmetry along the AP axis at the level of chromatin accessibility, well before the bulk of zygotic genome activation and pattern formation occurs (81).

In Clogmia, knockdown of the maternal *Cal-opa* transcript by isoform-specific RNA interference (RNAi) results in a symmetrical double abdomen phenotype, and posterior mis-expression of *Cal-opa* by mRNA injection prior to blastoderm formation causes a bicephalic (double head) phenotype. Early Cal-Opa activity is therefore both necessary and sufficient for establishing head-to-tail polarity of the Clogmia embryo. An alternative zygotic transcript isoform of Clogmia *odd-paired* appears during blastoderm cellularization at NC14 and exerts a conserved function in the late segmentation gene network but has no role in establishing head-to-tail polarity of the embryo (3, 82). No overlapping functions have been observed between the maternal and the zygotic *Cal-opa* isoforms, which encode nearly identical proteins, suggesting that Clogmia’s AD functions exclusively before NC14, in contrast to Bcd, which remains essential for AP patterning until the end of NC14 (65). However, in Drosophila, zygotic Opa pioneers chromatin accessibility in late NC14 embryos to advance the progression of segmentation and dorsoventral patterning (78, 83). We therefore asked whether the maternal *Cal-opa* gradient of Clogmia breaks axial symmetry by driving chromatin accessibility at gene loci that are transcribed in the anterior blastoderm before the beginning of NC14 and whether these genes are homologous to Bcd target genes.

To set the stage for addressing these questions we first assembled and annotated the Clogmia genome and identified homologs of Drosophila’s segmentation genes. We then characterized chromatin accessibility in single wild-type embryos at NC11, NC12, NC13, and both early and late NC14 to predict putative enhancer and promoter regions and to capture their stage-dependent accessibility. We then assessed how chromatin accessibility patterns and transcription are affected by *Cal-zld* activity. Finally, we examined how reduced maternal *Cal-opa* expression affects chromatin accessibility and TF gene expression during NC12 and NC13. Our results reveal groups of gene loci with similar dynamic changes in chromatin accessibility, point to direct roles of Cal-Zld in opening and closing chromatin, and provides evidence for a role of maternal Cal-Opa in opening chromatin during early blastoderm stages at a small set of loci with anterior expression, including homologs of *sloppy-paired* and *homeobrain.* We conclude that the conserved feature of anterior determination in Clogmia and Drosophila is symmetry breaking at the level of chromatin accessibility, even though specific target genes may differ between the two species.

## Results

### *De novo* assembly and annotation of the genome of *Clogmia albipunctata* reveals multiple lineage-specific duplications of early segmentation gene homologs

We generated a high quality *de novo* genome assembly of *Clogmia albipunctata* with 6 chromosome-size scaffolds (**Figure 1**, **Table 1, S1 Appendix**), consistent with the known chromosome number in this species (84, 85). The total length of the chromosome-size scaffolds is 304.3 Mb, close to genome size estimates based on flow cytometry data (316.6 Mb) (86). An additional 268 small scaffolds (< 1 MB) contained mostly repetitive sequences. We annotated this new reference genome using RNA-seq transcripts from embryonic, larval, pupal, and adult stages (**S1 Table**), protein sequences, and *ab initio* gene prediction software, and identified 15,047 protein coding genes. The annotated genome recovered 88%-94% of Complete Benchmarking Universal Single Copy Orthologs (BUSCO) (**Table 2**), in line with recently published dipteran genomes (87–89). While synteny between the chromosomes within the Cyclorrhapha clade can be highly conserved (time of divergence ∼145 million years ago) (88), synteny between the chromosomes of *D. melanogaster* and *C. albipunctata* (divergence time ∼250 million years ago) was essentially lost (**Figure 1**).

**Figure 1:**
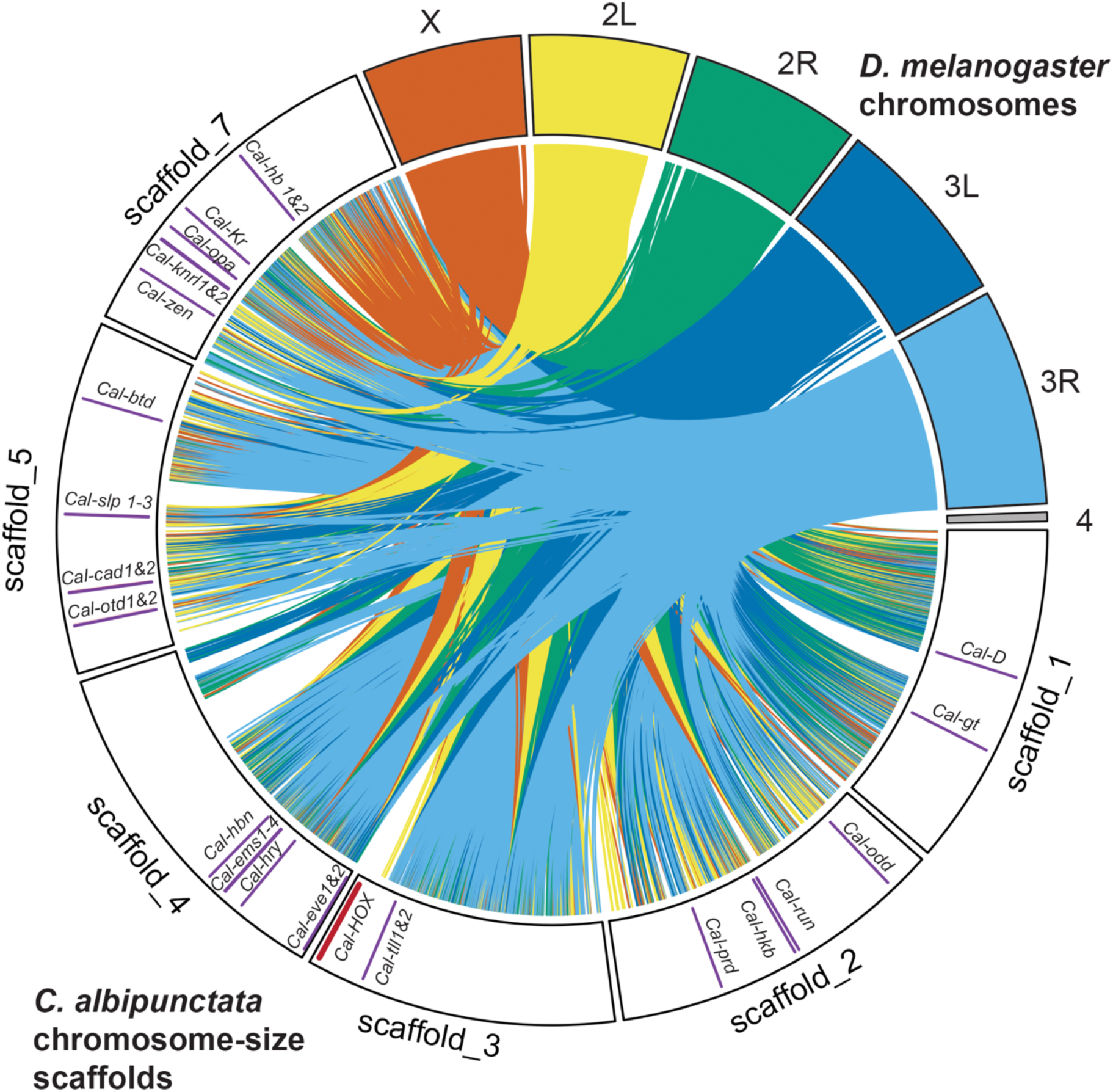
Synteny analysis of *Drosophila melanogaster* chromosomes and *Clogmia albipunctata* scaffolds. Synteny analysis between *Clogmia albipunctata* chromosomes-sized scaffolds *and Drosophila melanogaster* chromosomes. Groups of collinear genes are represented by contact lines connecting the positions shared between *D. melanogaster* chromosomes and *C. albipunctata* scaffolds. Colors correspond to *D. melanogaster* chromosomes. Positions of developmental genes of interest indicated by solid purple bars. Position of the HOX gene cluster indicated by a solid brown bar.

**Table 1:** *Clogmia albipunctata* reference assembly statistics and quality metrics. Completeness and continuity of the assembly are assessed by Benchmarking Universal Single-Copy Orthologs (BUSCO) and scaffold N50 and L50, respectively. Eukaryotic BUSCOs recovered: the percentage of complete universal orthologs from the eukaryotic BUSCO dataset identified in the assembly. Scaffold N50: The length (in bp) of the shortest scaffold such that all scaffolds of equal or greater length account for at least 50% of the total assembly length (174). L50: The minimum number of scaffolds whose collective length accounts for 50% of the total assembly length (175).

| Reference Assembly Statistics |  |
| --- | --- |
| Total length (bp) | 320,011,659 |
| Number of scaffolds | 274 |
| Scaffold N50 (bp) | 51,218,107 |
| Scaffold L50 | 4 |
| # of Ns per 100kb | 0.19 |
| GC (%) | 32.34% |
| Heterozygosity (%) | 0.018 - 0.038% |
| Masked (%) | 48.55% |
| Eukaryotic BUSCOs recovered (%) | 98.04% |

**Table 2:**
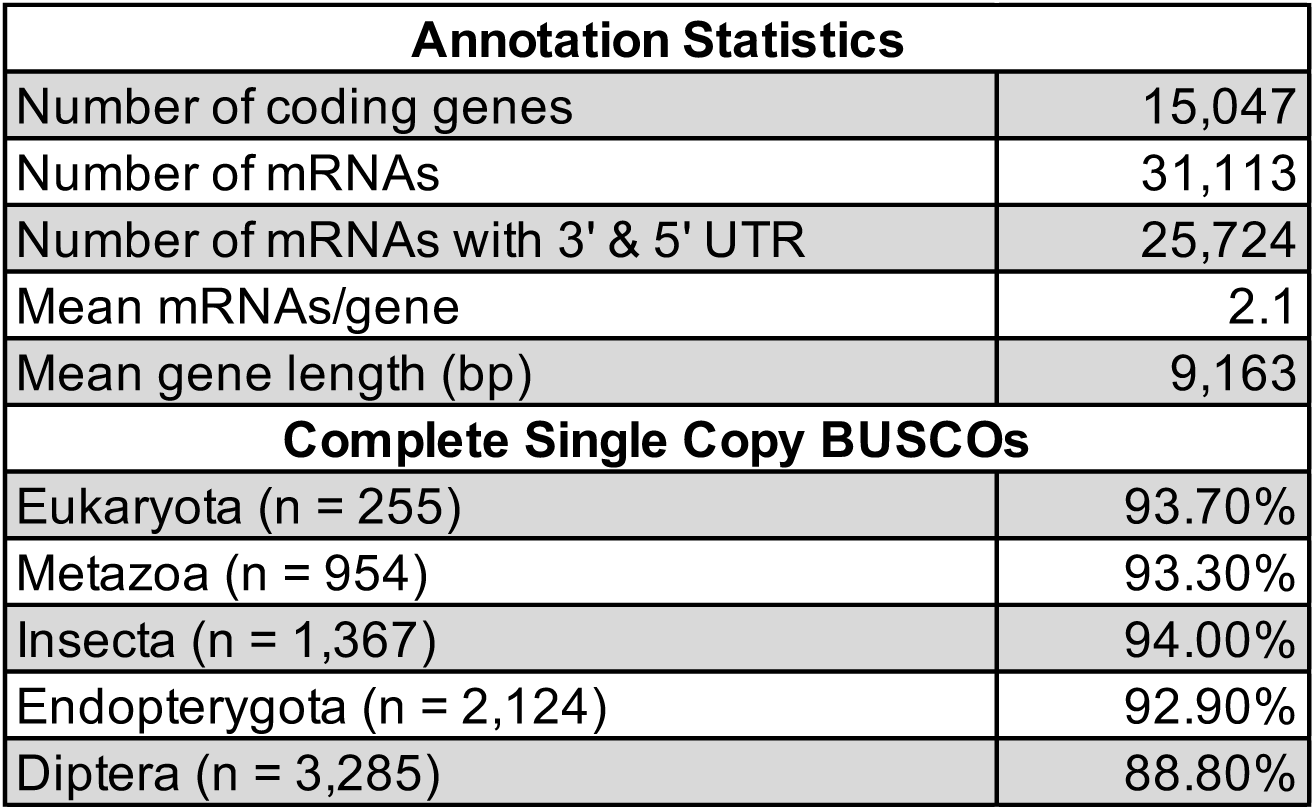
*Clogmia albipunctata* annotation statistics and BUSCO scores. The quality metrics include the percentage of complete universal orthologs identified in the annotation across five BUSCO datasets. The number of genes in each lineage-specific BUSCO database is shown in parentheses.

Next, we interrogated the genome for the presence of predicted early segmentation genes. As expected, no *bicoid* ortholog was found but we recovered two putative Hox class 3 genes to which *bicoid* belongs (90), including the previously reported homolog of *zerknüllt* (*Cal-zen*) (91) on scaffold 7 and a more diverged potential *Cal-zen* paralog in position 3 of the Hox gene complex on scaffold 3 (evm.TU.scaffold_3.2943.1.65be5b92). Our annotation also included homologs of *Drosophila melanogaster*’s early segmentation genes, including gap genes (*tailless*, *huckebein*, *orthodenticle/ocelliless*, *empty spiracles*, *buttonhead*, *hunchback*, *Krüppel*, *giant, knirps*), pair-rule genes (*even-skipped*, *fushi tarazu*, *odd-skipped*, *runt*, *sloppy-paired*, *paired, hairy, odd-paired*), and other important regulators of the segmentation network (*caudal*, *Dichaete*) (19, 21). Some of these genes have more than one copy in the Clogmia genome, including 2 copies of *hunchback* (*Cal-hb1, Cal-hb2*), *knirps/knirps-like* (*Cal-kni/knrl1, Cal-kni/knrl2*), *orthodenticle* (*Cal-otd1, Cal-otd2*), *even-skipped* (*Cal-eve1, Cal-eve2*), *tailless* (*Cal-tll1, Cal-tll2*) and *caudal* (*Cal-cad 1, Cal-cad2*), 3 copies of *sloppy-paired1/2/fd19B* (*Cal-slp1, Cal-slp2, Cal-slp3*), and 4 copies of *empty spiracles* (*Cal-ems1, Cal-ems2, Cal-ems3, Cal-ems4*) (**Figure 1, S2 Table**). Such gene duplications are a potential source of redundancies in Clogmia’s segmentation gene network (see below).

### Dynamic patterns of chromatin accessibility during consecutive blastoderm stages suggest coordinated regulation of associated genes

To identify accessible chromatin before and during the main wave of zygotic genome activation, we performed single embryo ATAC-seq at NC11, NC12, NC13, NC14 (before cellularization), and Late NC14 approximately 20 minutes before gastrulation. The embryos were staged *in vivo* using developmental time after egg activation at 25°C and morphological criteria as previously described (38) and validated under our laboratory conditions. At least three replicates were performed for NC11 though NC14, and two replicates were performed for Late NC14. We pooled the ATAC-seq data from all stages to create a master peak list of 32157 open chromatin regions (peaks), using MACS2 (92). We also computed the number of peaks for each developmental stage excluding stage-specific peaks that were not recovered as significant in the master peak list. In this analysis, we found that the overall number of peaks increases with each nuclear cycle, suggesting that the chromatin landscape is continuously changing (**Table 3**). Since promoter accessibility does not necessarily indicate gene expression (93), we divided the accessible chromatin regions into transcription start site (TSS) proximal regions (within 500 bp of TSS) and TSS distal regions (>500 bp) to facilitate the distinction of open promoters and other open cis-regulatory elements. ATAC-seq peaks that only mapped to intergenic or intronic sequence were treated as candidate enhancers or silencers (henceforth collectively referred to as enhancers). The relative proportion of candidate enhancers increased by about 10% from NC11 (46.8%) to NC14 (58%) and then decreased from NC14 to Late NC14 (52.7%) (**S1 Figure**). These observations suggest similar dynamic changes in chromatin accessibility during zygotic genome activation in Clogmia and Drosophila (94, 95).

**Table 3:**
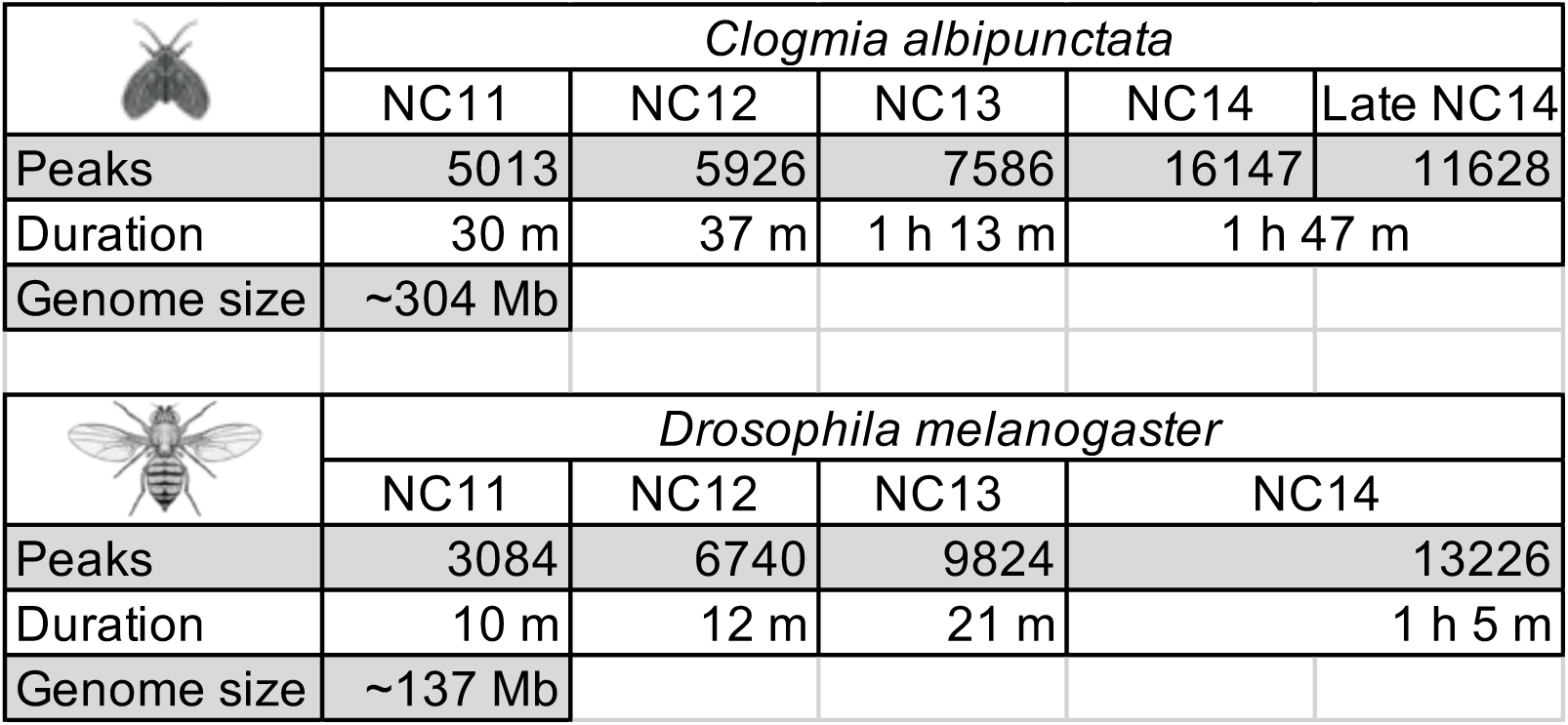
Comparison of chromatin accessibility peaks, nuclear cycle length in minutes, and haploid genome size of *Clogmia albipunctata* and *Drosophila melanogaster*. Peaks for *Drosophila melanogaster* are based on (73, 81) and nuclear cycle lengths are based on (38).

|  | *Clogmia albipunctata* |  |  |  |  |
| --- | --- | --- | --- | --- | --- |
|  | NC11 | NC12 | NC13 | NC14 | Late NC14 |
| Peaks | 5013 | 5926 | 7586 | 16147 | 11628 |
| Duration | 30 m | 37 m | 1 h 13 m | 1 h 47 m |  |
| Genome size | ~304 Mb |  |  |  |  |
|  | *Drosophila melanogaster* |  |  |  |  |
|  | NC11 | NC12 | NC13 | NC14 |  |
| Peaks | 3084 | 6740 | 9824 | 13226 |  |
| Duration | 10 m | 12 m | 21 m | 1 h 5 m |  |
| Genome size | ~137 Mb |  |  |  |  |

To identify peaks with dynamic patterns of chromatin accessibility, we compared the degree of accessibility of each peak between successive timepoints (**Figure 2A-D**) using DESeq2 (96). This analysis identified a total of 6930 dynamically regulated peaks (adjusted p-value ≤ 0.05 and |log_2_ Fold Change| ≥ 1; 21.6% of all 32157 peaks) and enabled us to quantitatively determine the behavior of each peak (i.e., maintaining, gaining or losing accessibility) between consecutive developmental stages. We then clustered dynamic peaks into groups based on shared changes in accessibility from stage to stage and delineated 21 dynamic groups (**S2 Figure**). Nearly 80% (5403 / 6930) of the dynamic peaks fell into one of 8 groups, which reveal the major accessibility trends we observe in the data (**Figure 2E)**. Many of the dynamic peaks closed (14.6%; Groups 5,13) or opened (∼20.4%; Groups 8,12) gradually over time. The remaining dynamic peak groups exhibited more complex dynamics between NC11 and Late NC14, commonly with an inflection at NC13 (e.g., Groups 3,1,15,14; see also **S2 Figure**). These observations suggest that remodeling of chromatin accessibility between NC11 and Late NC14 at individual peaks often follows patterns that are shared by many peaks and may reflect co-regulation of the associated genes.

**Figure 2:**
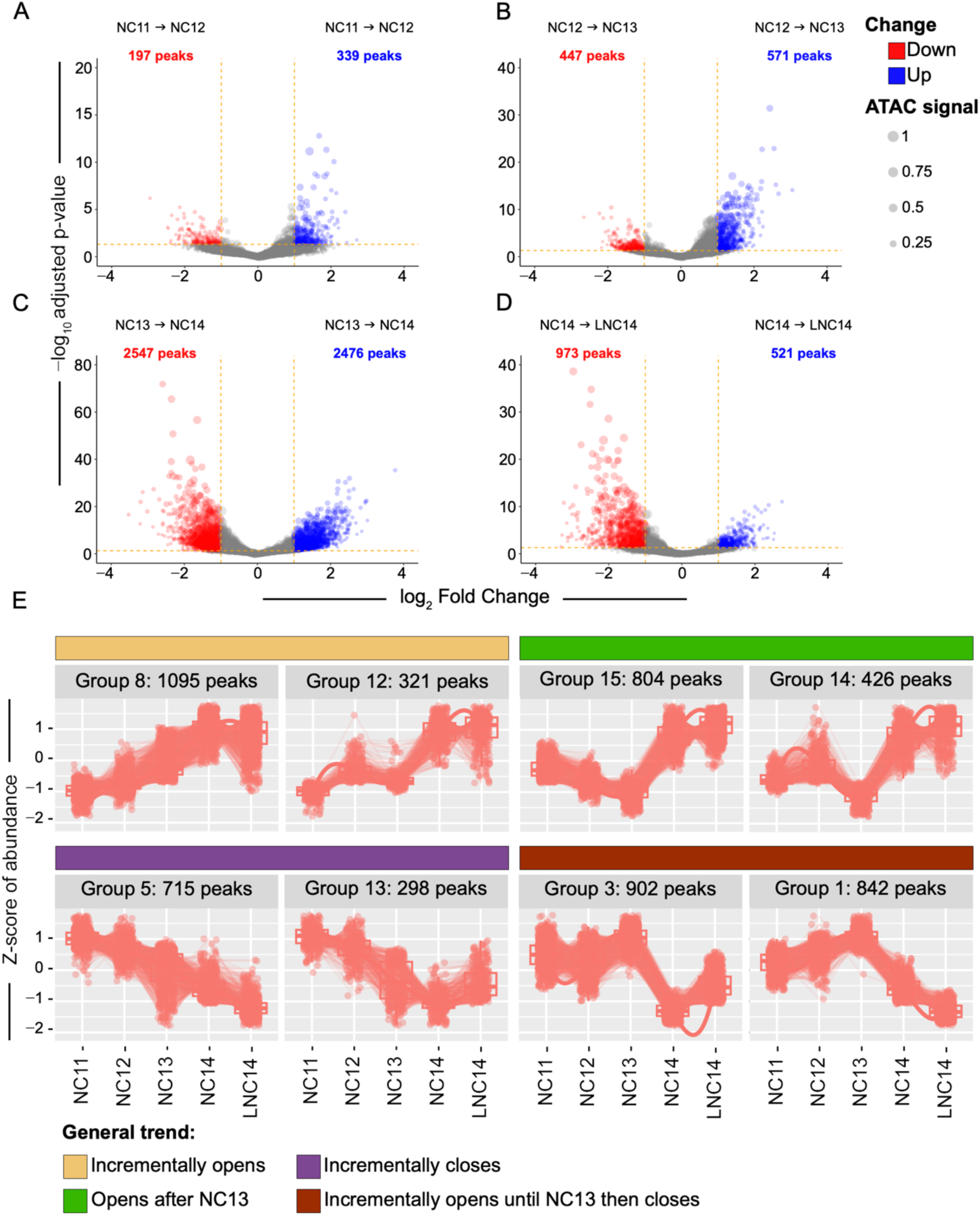
Changes in chromatin accessibility during NC11 to Late NC14. A-D) Volcano plots of differentially accessible ATAC-seq peaks between consecutive developmental stages. Significant change of adjusted p-value ≤ 0.05 and |log_2_ Fold Change| ≥ 1highlighted; red (log_2_ Fold Change ≤ –1) and blue (log_2_ Fold Change ≥ 1). E) Dynamic peak groups produced using DEGreport (Pantano, 2017) (R package version 1.38.5). Y-axis, Z-score of abundance. X-axis, stage.

### *Cal-zld* is required for both opening and closing chromatin at the blastoderm stage in Clogmia

Previous studies in Drosophila have shown that temporal changes in chromatin accessibility are regulated by the interplay of transcription factors that can interact with nucleosome-bound DNA (pioneer factors) and transcription factors that have limited or no ability to initiate chromatin accessibility (49, 56). The earliest changes in chromatin accessibility are primarily driven by the pioneer factor Zld (46, 54, 58, 94, 97). As zygotic genome activation accelerates, additional pioneer factors, such as GAF (63), CLAMP (61, 62), and Opa (78, 83) are required for maintaining, modulating, or gaining chromatin accessibility. In Clogmia, a motif enrichment analysis of peaks with the MEME software package (98, 99) revealed an enrichment of a Zelda-binding motif (5’-CAGGTA), GA-rich motifs recognized by GAF and CLAMP (63, 100–102), in addition to (CA)_n_ motifs recognized by Combgap (Cg) (103), and others (**S3 Figure**). Opa-binding motifs were not recovered in this analysis. These results are comparable to findings in Drosophila (78, 81, 83) and suggest that motif enrichment within chromatin accessible at ZGA is very similar between Drosophila and Clogmia.

Homologs of *zelda* have been identified in many insects and some crustaceans (104), and several transcriptomic studies suggest that Zld’s function as pioneer transcription factor during zygotic genome activation is conserved across insects (105–109). However, these comparative studies did not directly test for Zld-dependent effects on chromatin accessibility. We identified Clogmia *zld* (*Cal-zld*) (**S4 Figure**) and confirmed high levels of *Cal-zld* expression at preblastoderm and blastoderm stages (**S5 Figure**). Loss of Zld in *Drosophila* results in a failure to activate a large cohort of early zygotic genes, including those associated with the process of cellularization (46). To examine phenotypic effects of *Cal-zld* depletion, batches of ∼30 embryos were injected with *Cal-zld* dsRNA (double-stranded RNA) within the first hour of development (NC2/3). Injected embryos were monitored *in vivo* and allowed to develop until the end of NC14 (roughly 7-7.5 hours after injection). Across three independent replicates, a subset of injected embryos reached NC14 but failed to cellularize and developed abnormally (**S1 movie**). This phenotype is comparable to the phenotype of *zld* mutant Drosophila embryos (**S2 movie**) (58) and suggests that the role of Zld for the regulation of early zygotic genes is conserved in Clogmia. More than half of the injected embryos (33/59) did not reach the blastoderm stage while early mortality of an uninjected control group was ∼27% (6/22), but it is unclear whether this difference reflects a requirement for Zelda earlier in development or an artifact of the injection procedure.

To assess *Cal-zld*’s role in shaping chromatin accessibility and zygotic transcription, we adapted our ATAC-seq protocol to isolate RNA from single-embryo ATAC-seq library preparations of Late NC14 *Cal-zld* RNAi embryos (see *Materials and Methods*). To identify RNAi embryos with maximally reduced *Cal-zld* mRNA levels, we used the RNA-seq data to measure *Cal-zld* expression and selected embryos with knockdown efficiencies greater than 75% (n= 4/8, **S6 Figure** and see *Materials and Methods*). We then compared gene expression and ATAC-seq peaks of these *Cal-zld* RNAi embryos to those of uninjected control embryos using DESeq2. Loss of *Cal*-*zld* affected 2230 genes of which 865 (38.8%) were downregulated and 1365 (61.2%) were upregulated. We then compared genes differentially expressed in *Cal-zld* RNAi embryos to genes we found to be expressed maternally, zygotically, or both (**Figure 3A, S7A Figure** and *Materials and Methods*). 691 genes were classified as zygotically expressed of which 209 (30.2%) were downregulated in *Cal-zld* RNAi embryos and 63 (9.1%) were upregulated (corresponding to 24.2% of all 865 downregulated and 4.6% of all 1365 upregulated genes, respectively). 6658 genes were classified as expressed maternally and zygotically of which 593 (8.9%) were downregulated and 1232 (18.5%) were upregulated. Finally, of the 227 genes classified as maternally expressed, only 5 (2.2%) were found to be downregulated and 14 (6.2%) upregulated in *Cal-zld* RNAi embryos. Since the strictly zygotic genes were ∼3 times more often downregulated than upregulated (209 versus 63), our results suggest that *Cal-zld* plays an important role in activating zygotic genes (**Figure 3A),** consistent with the role of Zld during zygotic genome activation in Drosophila embryos (46, 54). Changes in the levels of maternal transcripts in response to *Cal-zld* RNAi could reflect indirect posttranscriptional roles of *Cal-zld* in their regulation.

**Figure 3:**
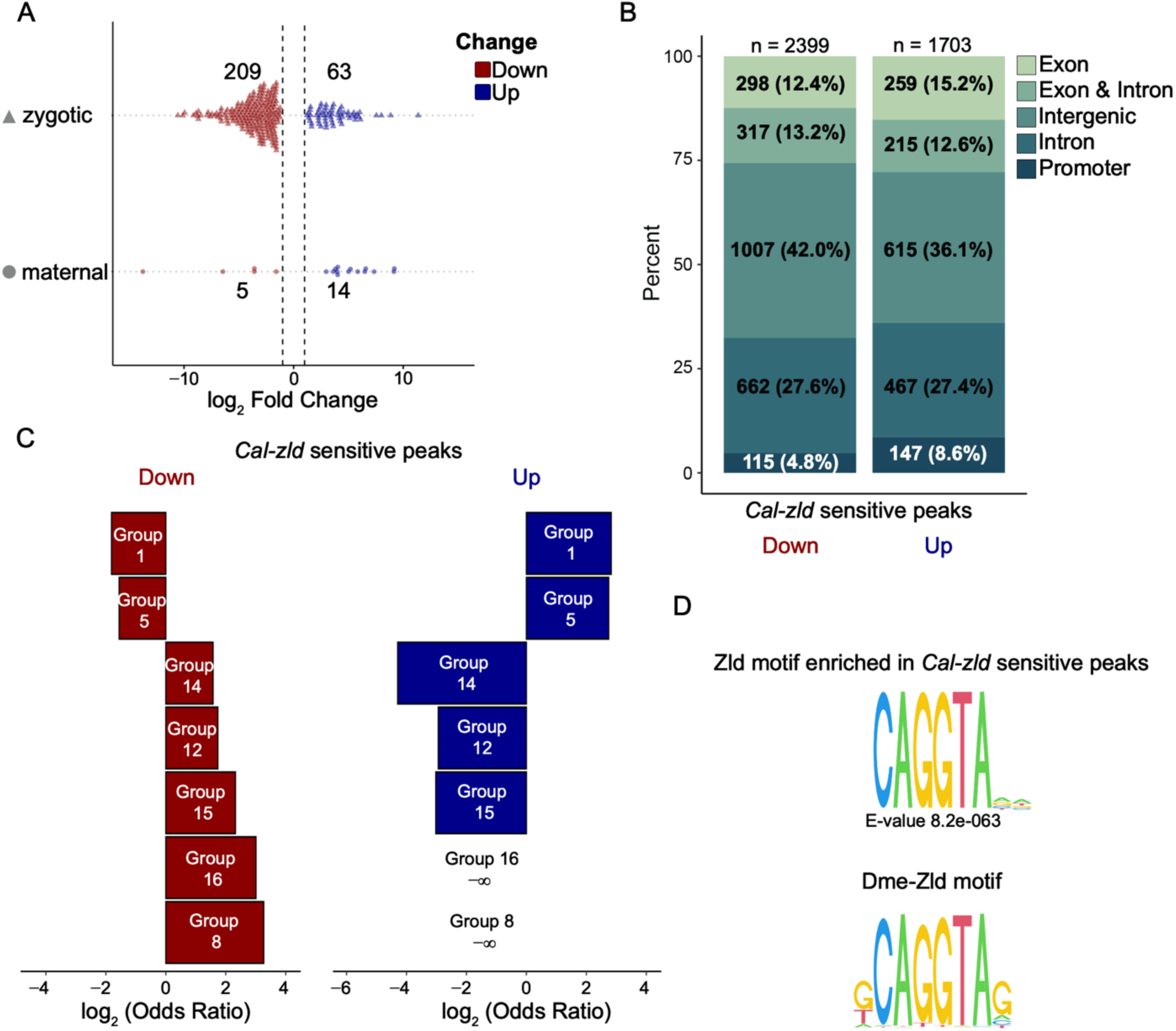
Cal-Zld regulates chromatin accessibility and zygotic transcription positively and negatively. A) Observed expression change of maternal or zygotic genes that are differentially expressed in *Cal-zld* RNAi embryos. On the y-axis genes are grouped by maternal or zygotic classification (see *Materials and Methods*). Log_2_ Fold Change on the x-axis. Only genes with an adjusted p-value ≤ 0.05 and |log_2_ Fold Change| ≥ 1 are shown. B) Genomic distribution of *Cal-zld* sensitive peaks that were up-regulated (blue) and down-regulated (red). C) Dynamic groups whose accessibility increases over time are more likely to be *Cal-zld* sensitive peaks that lose accessibility (Group 8 p-value, 3.4e-176; Group 16 p-value, 8.7e-05; Group 15 p-value, 1.3e-55; Group 12 p-value, 4.6e-12; Group 14 p-value, 7.2e-12) unlike dynamic groups whose accessibility decrease over time or after NC13 (Group 5 p-value, 3.5e-08; Group 1 p-value, 2.7e-12). Dynamic groups whose accessibility decreases over time are more likely to be *Cal-zld* sensitive peaks that gain accessibility (Group 5 p-value, 1.7e-70; Group 1 p-value, 1.4e-95), unlike dynamic groups whose accessibility increases over time (Group 8 p-value, 3.4e-25; Group 16 p-value, 6.3e-01; Group 15 p-value, 1.8e-11; Group 12 p-value, 9e-05; Group 14 p-value, 2.6e-07). Odds ratios and p-values were calculated by a two-sided Fisher’s exact test on contingency tables constructed by the presence/absence of dynamic groups in *Cal-zld* sensitive peaks up or down. Only groups with a log_2_(Odds Ratio) > 1.5 in at least one comparison are shown. D) Position weight matrix (PWM) logo of the Zld motif enriched in *Cal-zld* sensitive peaks as identified by MEME (top) and the canonical Zld motif in *Drosophila melanogaster*, shown for reference (bottom).

At the level of chromatin accessibility, *Cal-zld* depletion affected 4102 of all 32157 peaks (12.8%). 2399 of them lost accessibility (58.4%) and 1703 gained accessibility (41.5%). The majority (67%) of the peaks mapped to intergenic or intronic sequences (**Figure 3B)**. Depletion of *Cal-zld* mRNA affected both dynamic peaks and non-dynamic peaks (**S7B Figure**). Dynamic peaks were affected by *Cal-zld* knockdown in different ways (**Figure 3C**). Peaks in groups that progressively gained accessibility in wild-type embryos (Groups 8, 12, 15 and 14) were more likely to be *Cal-zld* dependent and to close in *Cal-zld* RNAi background (**Figure 3C, *left*** and **S8A Figure**). This observation is consistent with an important role of Cal-Zld in opening new chromatin regions during zygotic genome activation. Additionally, our data suggest that Cal-Zld also plays an important role in closing chromatin regions during zygotic genome activation (**S8B Figure**). For example, peaks in dynamic groups whose accessibility decreased in wild-type with each nuclear cycle or that closed after NC13 (Groups 5 and 1) were more likely to require Cal-Zld to close (**Figure 3C, *right*** and **S8B Figure**).

To assess if the changes in accessibility we observed were a direct consequence of *Cal-zld* knockdown, we performed two types of motif analyses. We first performed a motif enrichment analysis with MEME on *Cal-zld* sensitive peaks to identify motifs enriched centrally within 50 bp of the peak summit (**S9 Figure**). The Zld motif was enriched in peaks that lost accessibility following *Cal-zld* knockdown, consistent with a direct role of Cal-Zld in nucleosome depletion. A Zld motif was not enriched in peaks that gained accessibility following *Cal-zld* knockdown (**Figure 3D** and **S9 Figure)**. The enrichment of Zld motifs in only one class of *Cal*-*zld* sensitive peaks may reflect the prevalence of Cal-Zld binding in those regions. In support of this hypothesis, we note that both motif affinity and occupancy positively correlate with Zld’s pioneering activity in Drosophila (56, 110). Next, we extended the search for Cal-Zld motifs to the full width of each peak, using Zld motifs derived from our MEME analysis in Clogmia (5-CAGGTA; cf. **S3 Figure)** and experimental data in *Drosophila melanogaster* (Fly Factor Survey) (111). Overall, 2857 of the 4102 *Cal-zld* sensitive peaks contained at least one Zld motif within the full peak width. Consistent with the motif enrichment analysis, a Zld motif was prevalent in peaks whose accessibility was lost following *Cal-zld* depletion (81%, 1952/2399 peaks). However, we also observed Zld motifs in peaks whose accessibility increased following *Cal-zld* knockdown (53%, 905/1703 peaks). Therefore, these regions may close due to Cal-Zld activity directly. Taken together, our findings suggest that Cal-Zld plays a central role in both opening and closing chromatin accessibility in pre-gastrulation embryos.

### Chromatin accessibility and gene expression at NC12 and NC13 following knockdown of maternal *Cal*-*opa*

Having established that *Cal-zld* regulates chromatin accessibility in Clogmia embryos, we asked whether maternal *Cal-opa* affects chromatin accessibility during early stages of axial pattern formation. Since *Cal-opa* exerts its symmetry-breaking function in axis specification prior to the onset of zygotic *Cal-opa* expression during blastoderm cellularization at mid NC14 (3), we were able to examine the function of *Cal-opa* without confounding effects of zygotic *Cal-opa* activity. To assess the impact of maternal Cal-Opa activity on chromatin accessibility, we performed ATAC-seq and RNA-seq on single *Cal-opa* RNAi embryos at NC12 (6 replicates) and NC13 (8 replicates). In parallel, we processed stage-matched embryos that were prepared for injection under oil but not injected (alignment controls; 4 replicates for each developmental stage), and embryos that were injected with dsRNA of the extraneous gene *DsRed* (injection controls; 3 replicates for each stage). To identify RNAi embryos with maximally reduced *Cal-opa* mRNA levels, we used the RNA-seq data to measure *Cal-opa* expression and selected embryos with knockdown efficiencies greater than 75% (**S10 Figure** and see *Materials and Methods*). We identified 4 embryos at NC12 and 5 embryos at NC13 with strong *Cal-opa* knockdowns and assessed changes in chromatin accessibility in these embryos by performing DESeq2 analysis with the alignment control group as reference.

At NC12, very few changes in chromatin accessibility were observed in response to *Cal-opa* RNAi. Out of the 86 differentially accessible regions, 79 (91.9%) lost accessibility and 7 (8.1%) gained accessibility (**Figure 4A**). At NC13, depletion of *Cal-opa* had a more pronounced effect on chromatin accessibility, with 250/292 (85.6%) peaks losing accessibility and 42/292 (14.4%) gaining accessibility **(Figure 4B**). Most of the differentially accessible regions mapped to putative enhancer regions (**S11 Figure**). We also performed a motif enrichment analysis using the MEME suite on all peaks as well as the peaks that either lost or gained accessibility in *Cal-opa* RNAi embryos at either NC12 or NC13 (**Figure 4C**). In regions that lost accessibility in *Cal-opa* RNAi embryos, the most enriched motifs were GAF/CLAMP and Opa binding motifs. In addition, we recovered motifs that have not been shown to be bound by a TF in Drosophila. Regions that gained accessibility were enriched in DNA binding motifs of GAF/CLAMP, Abdominal-B (Abd-B), Twist (Twi), and two motifs with no associated transcription factor.

**Figure 4:**
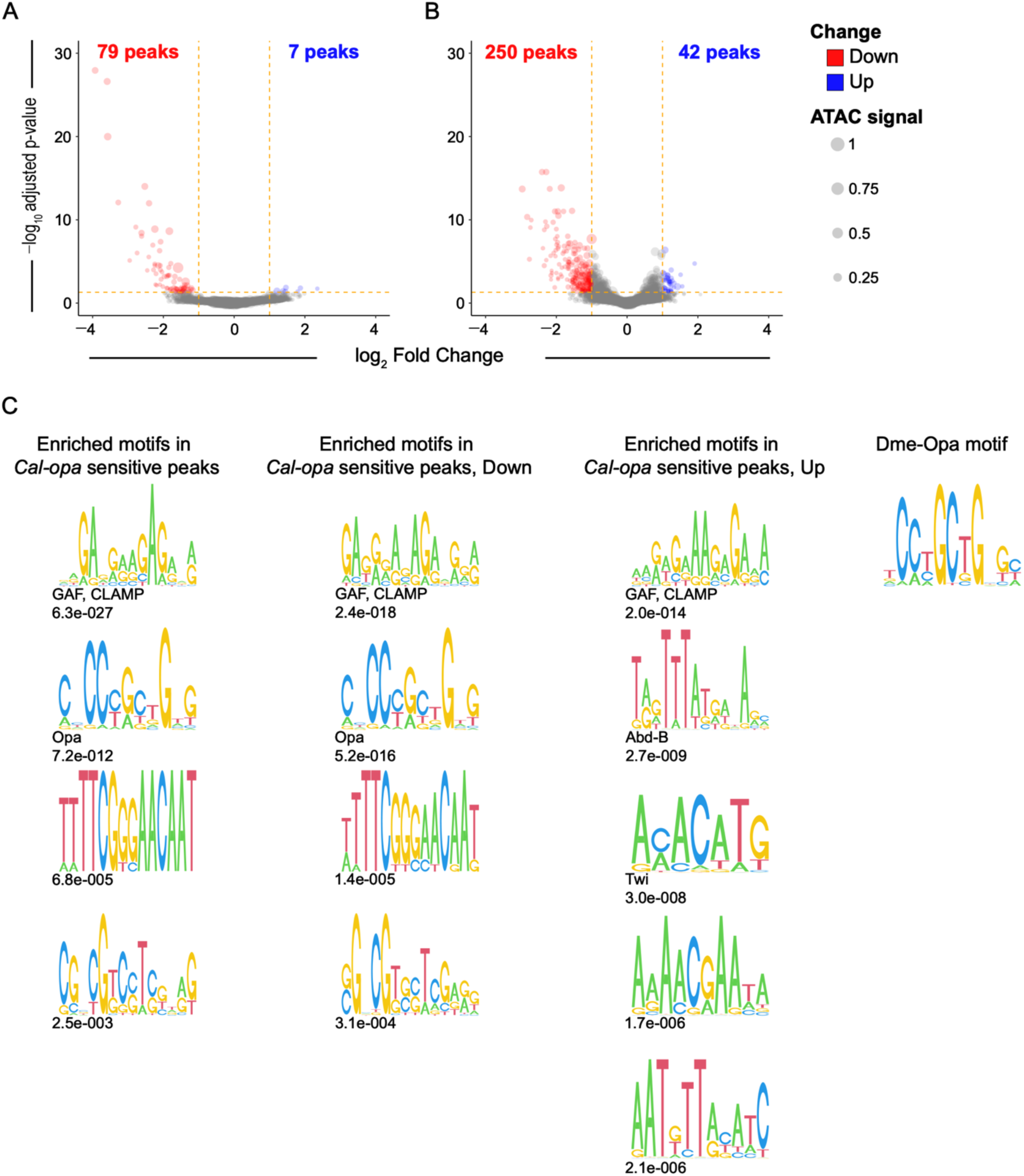
Cal-Opa is required for establishing a selective set of chromatin accessibility peaks. Volcano plots of differentially accessible peaks between the *Cal-opa* RNAi group and the alignment control group at A) NC12 and B) NC13. Significant change of adjusted p-value ≤ 0.05 and |log_2_ Fold Change| ≥ 1 highlighted; red (log_2_ Fold Change ≤ –1) and blue (log_2_ Fold Change ≥ 1). C) PWM logos of motifs enriched in all *Cal-opa* sensitive peaks (n = 326), in *Cal-opa* sensitive peaks that lose accessibility (n = 277), and in *Cal-opa* sensitive peaks that gain accessibility (n = 49), as identified by the MEME suite. Data combined for both NC12 and NC13 stages. DNA binding proteins of *Drosophila melanogaster* that bind to these or similar motifs as identified by MEME are indicated. We include CLAMP in this list as it has been shown to bind GA-rich motifs (see text). E-value denotes the significance of recovered motifs. The canonical Opa motif in *Drosophila melanogaster* is shown for reference.

Next, we determined the number of *Cal-opa* sensitive peaks that contained putative Cal-Opa binding sites, including Opa motifs derived from our MEME analysis and a MEME analysis of ATAC-seq data from Drosophila (78). At NC12, 63 of the 79 peaks losing accessibility and none of the 7 peaks gaining accessibility in response to reduced *Cal-opa* activity contained at least one Opa motif (**S2 Appendix**). At NC13, 112 of the 250 peaks losing accessibility and 6 of the 42 peaks gaining accessibility in response to reduced *Cal-opa* activity contained at least one Opa motif (**S3 Appendix**). These results support the hypothesis that Opa directly promotes chromatin accessibility during the early phase of zygotic genome activation.

### Maternal Cal-Opa drives chromatin accessibility and expression at *homeobrain* and *sloppy-paired* loci in complementary anterior domains

The early zygotic segmentation gene network of dipterans is dominated by TF-encoding genes, such as the gap genes and pair-rule genes (19, 21, 29, 39, 112–115). Since Bcd breaks axial symmetry by targeting these genes in a concentration-dependent manner, we focused our search of Cal-Opa targets on TF genes that are expressed during blastoderm formation. Using the RNA-seq data from the embryos that were used to identify regions of differential chromatin accessibility, we identified 84 differentially expressed genes at NC12, including 11 transcription factor genes, all of which were downregulated upon *Cal-opa* knockdown (**Figure 5A** and **S2 Appendix**). At NC13, we identified 1047 differentially expressed genes, including 116 putative transcription factor genes of which 67 were downregulated and 49 were upregulated in *Cal-opa* RNAi embryos (**Figure 5B** and **S3 Appendix**). We then asked which of the differentially expressed transcription factor genes are located near open chromatin regions (ATAC-seq peaks) with an Opa motif. This analysis was restricted to peaks within a 40 kb region centered on each gene body of interest and further narrowed the list of potential key target genes of Cal-Opa to 10 genes at NC12 and 72 genes at NC13 (**S2 Appendix and S3 Appendix)**. At NC12, 4 of the 10 candidate genes were associated with at least one Cal-Opa dependent ATAC-seq peak, including *Cal-opa,* and homologs of *homeobrain* (*Cal-hbn*) and *sloppy-paired* (*Cal-slp1*, *Cal-slp2*) (**Figure 6, S3 Table,** and **S12 Figure**). At NC13, we thereby identified *Cal-opa, Cal-slp1*, *Cal-slp2* and *Cal-hbn* in addition to Clogmia orthologs of *Dichaete* (*Cal-D*), *Krüppel* (*Cal-Kr*), and an unidentified C2H2 zinc finger gene among the 72 candidates (**Figure 7, S3 Table, S12 Figure and S13 Figure**).

**Figure 5:**
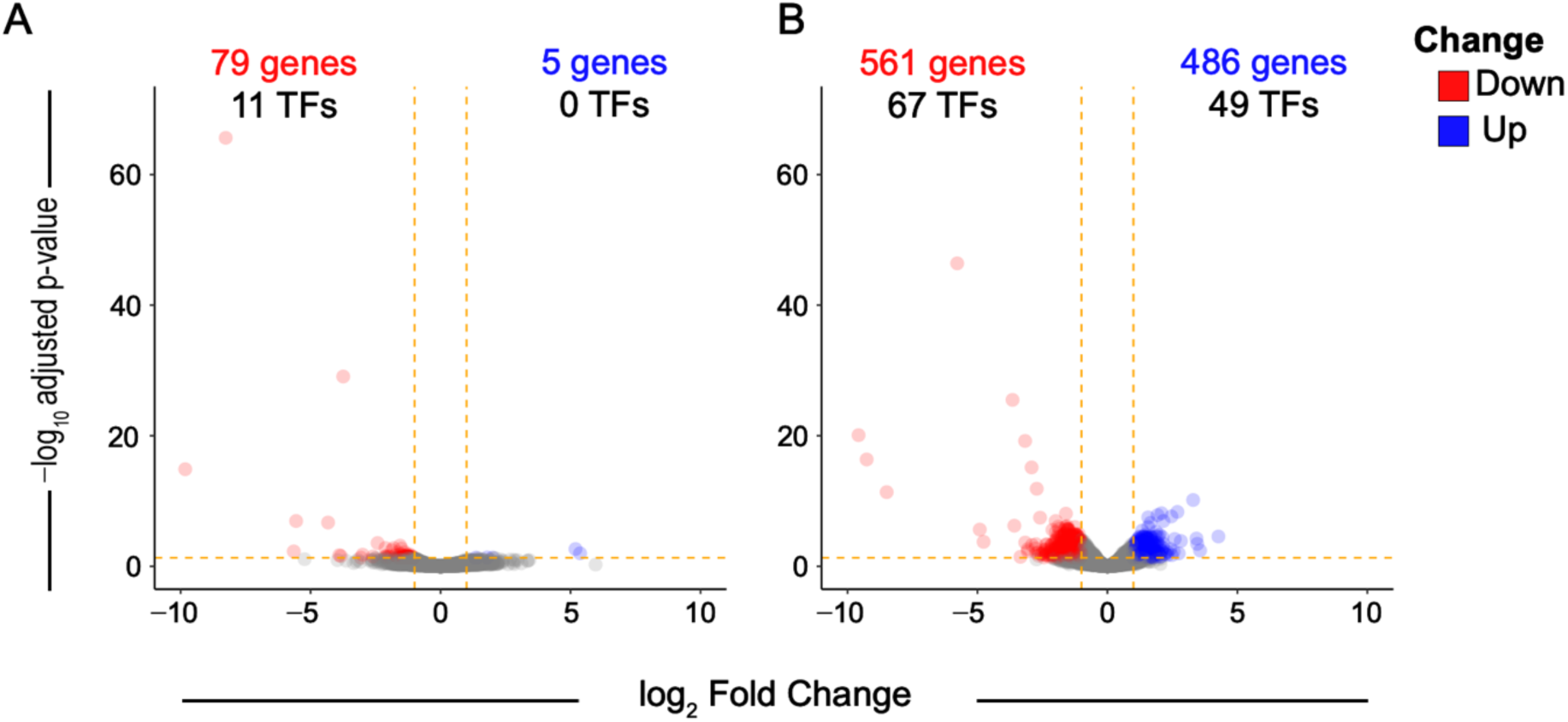
*Cal-opa* knockdown leads to the mis-regulation of a small set of transcription factor genes. Volcano plots of differentially expressed genes between the *Cal-opa* RNAi group and the alignment control group at A) NC12 and B) NC13. Significant change of adjusted p-value ≤ 0.05 and |log_2_ Fold Change| ≥ 1 highlighted; red (log_2_ Fold Change ≤ –1) and blue (log_2_ Fold Change ≥ 1). The number of transcription factors (TFs) is indicated in the text.

**Figure 6:**
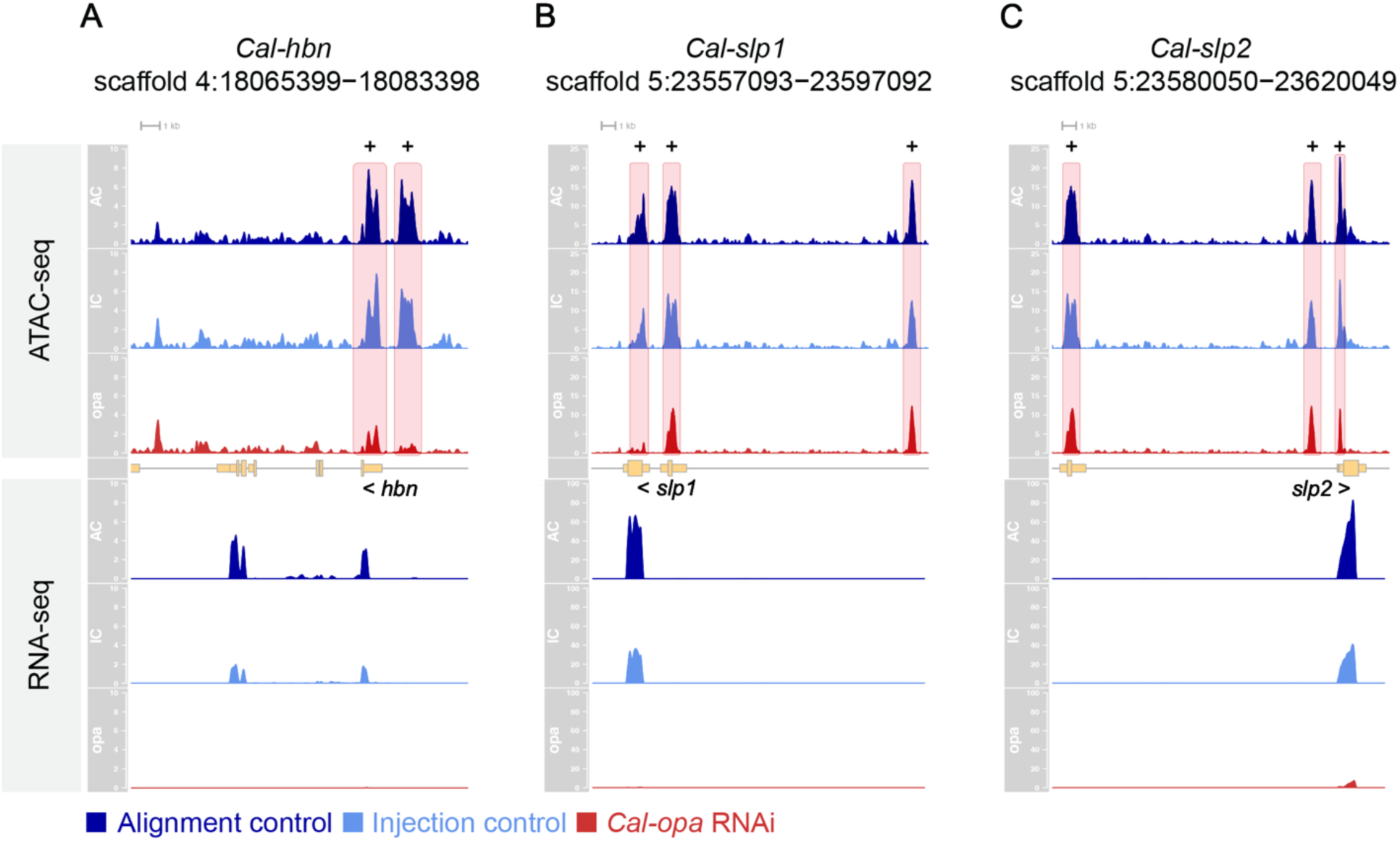
Key Cal-Opa targets at NC12. Gene loci of A) *Cal-hbn*, B) *Cal-slp1*, and C) *Cal-slp2* at NC12. Top tracks: CPM of ATAC-seq data (Alignment control in blue, Injection control in red, *Cal-opa* RNAi in yellow). Middle tracks show gene models in pale yellow. Bottom tracks: CPM of RNA-seq data. Peaks with significant changes in accessibility are highlighted in pink. Peaks containing Opa motifs are marked with a plus sign. The y-axis is the same for all tracks of the same data type at each locus. *Cal*-*hbn* ATAC-seq: 0-10 CPM. *Cal-hbn* RNA-seq: 0-10 CPM. *Cal-slp1* ATAC-seq: 0-25 CPM. *Cal-slp1* RNA-seq: 0-100 CPM. *Cal-slp2* ATAC-seq: 0-25 CPM. *Cal*-slp2 RNA-seq: 0-100 CPM. Blue: Alignment control. Light blue: Injection control. Red: *Cal-opa* RNAi.

**Figure 7:**
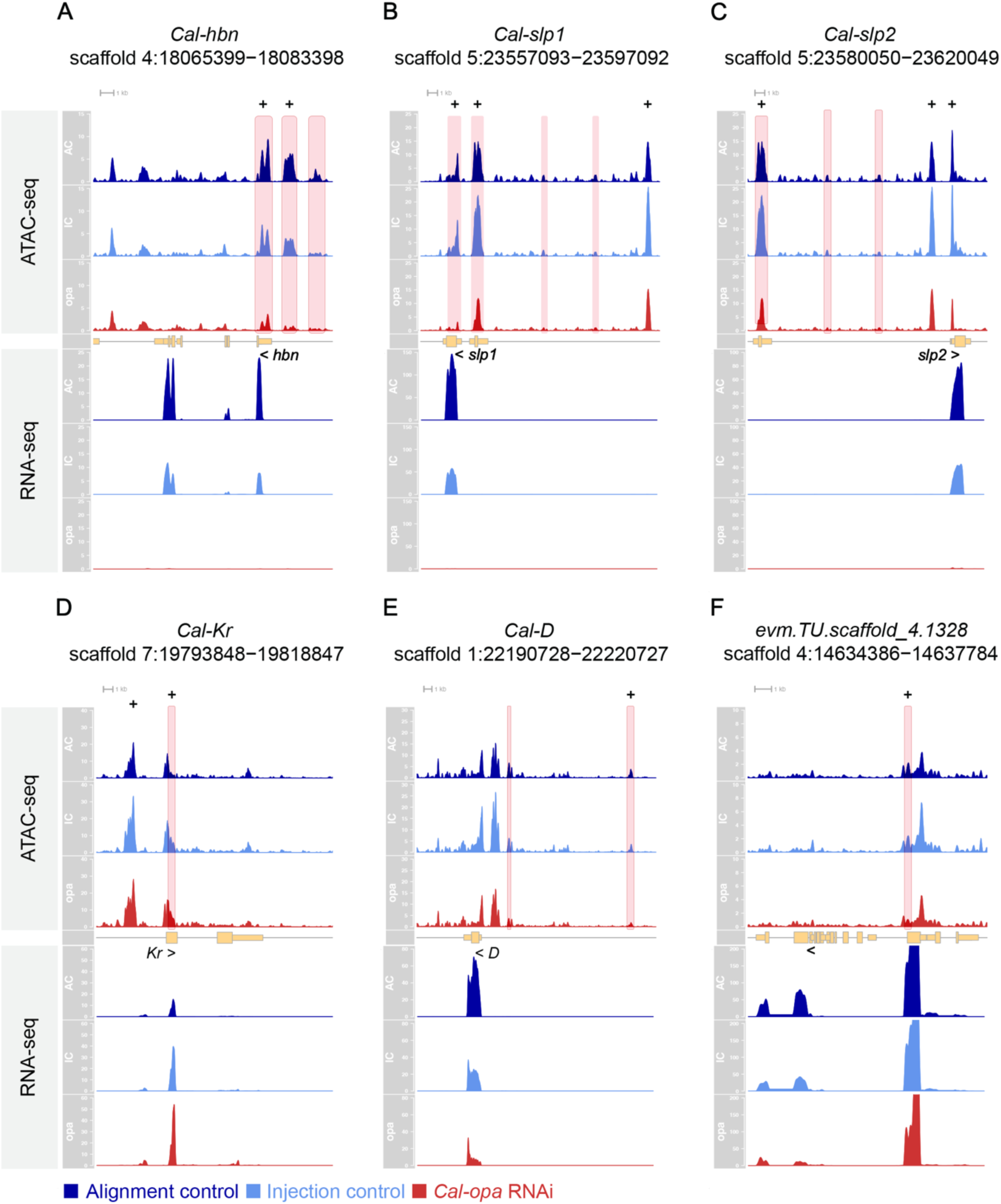
Key Cal-Opa targets at NC13. Gene loci of A) *Cal-hbn*, B) *Cal-slp1*, C) *Cal-slp2*, D) *Cal-Kr*, E) *Cal-D* and F) *evm.TU.scaffold_4.1328* at NC13. Top tracks: CPM of ATAC-seq data. Middle track gene model in pale yellow. Bottom tracks: CPM of RNA-seq data. Peaks with significant changes in accessibility highlighted in pink. Peaks containing Opa motifs are marked with a plus sign. The y-axis is the same for all tracks of the same data type at each locus. *Cal-hbn* ATAC-seq: 0-15 CPM. *Cal-hbn* RNA-seq: 0-25 CPM. *Cal*-*slp1* ATAC-seq: 0-25 CPM. *Cal-slp1* RNA-seq: 0-150 CPM. *Cal*-*slp2*: ATAC-seq: 0-25 CPM. *Cal*-*slp2* RNA-seq: 0-100 CPM. *Cal*-*Kr* ATAC-seq: 0-40 CPM. *Cal*-*Kr* RNA-seq: 0-60 CPM. *Cal*-*D* ATAC-seq: 0-30 CPM. *Cal*-*D* RNA-seq: 0-80 CPM. *evm.TU.scaffold_4.1328* ATAC-seq: 0-10 CPM. *evm.TU.scaffold_4.1328* RNA-seq: 0-200 CPM. Blue: Alignment control. Light blue: Injection control. Red: *Cal-opa* RNAi.

We tested the robustness of these results by repeating the analysis using the injection control group as reference. At NC12, we recovered *Cal-opa*, *Cal-hbn, Cal-slp1,* and *Cal-slp2* as candidate targets, but only three of them (*Cal-opa*, *Cal-hbn, Cal-slp1*) were associated with Cal-Opa dependent ATAC-seq peaks with Cal-Opa binding sites. In the case of *Cal-slp2*, the reduction of these peaks was insignificant (**S3 Table** and **S2 Appendix**). At NC13, we recovered *Cal-hbn, Cal-slp1 and Cal-slp2* (**S3 Table** and **S3 Appendix**). *Cal-opa*, *Cal-Kr*, *Cal-D* and the unidentified C2H2 zinc finger gene were not recovered in this analysis at NC13 because they did not meet the threshold for statistical significance (settings identical for NC12 and NC13). We therefore focused on Clogmia’s *homeobrain* and *sloppy-paired* homologs.

*Cal-hbn* and *Cal-slp1* expression was strictly zygotic and detected during and after NC11 when both genes were expressed in non-overlapping anterior domains (**Figure 8A, B**). Maternal *Cal-slp2* transcript was previously found to be distributed in a shallow anterior-to-posterior gradient and may contribute to the embryo’s head-to-tail polarity in subtle ways (3). Zygotic *Cal-slp2* expression during and after NC11 was detected in an anterior stripe (**Figure 8C**). A third *sloppy-paired* homolog that we identified in the Clogmia genome (evm.TU.scaffold_5.987; **S14 Figure and S15 Figure**) was expressed at low levels at NC12 and NC13 and robustly at NC14. The expression of this gene was reduced in *Cal-opa* RNAi embryos but associated ATAC-seq peaks did not contain Opa-binding motifs. Taken together, our findings suggest that *Cal-hbn*, and *Cal-slp1* and *Cal-slp2* are direct early targets of Clogmia’s AD and that Cal-Opa serves as their transcriptional activator.

**Figure 8:**
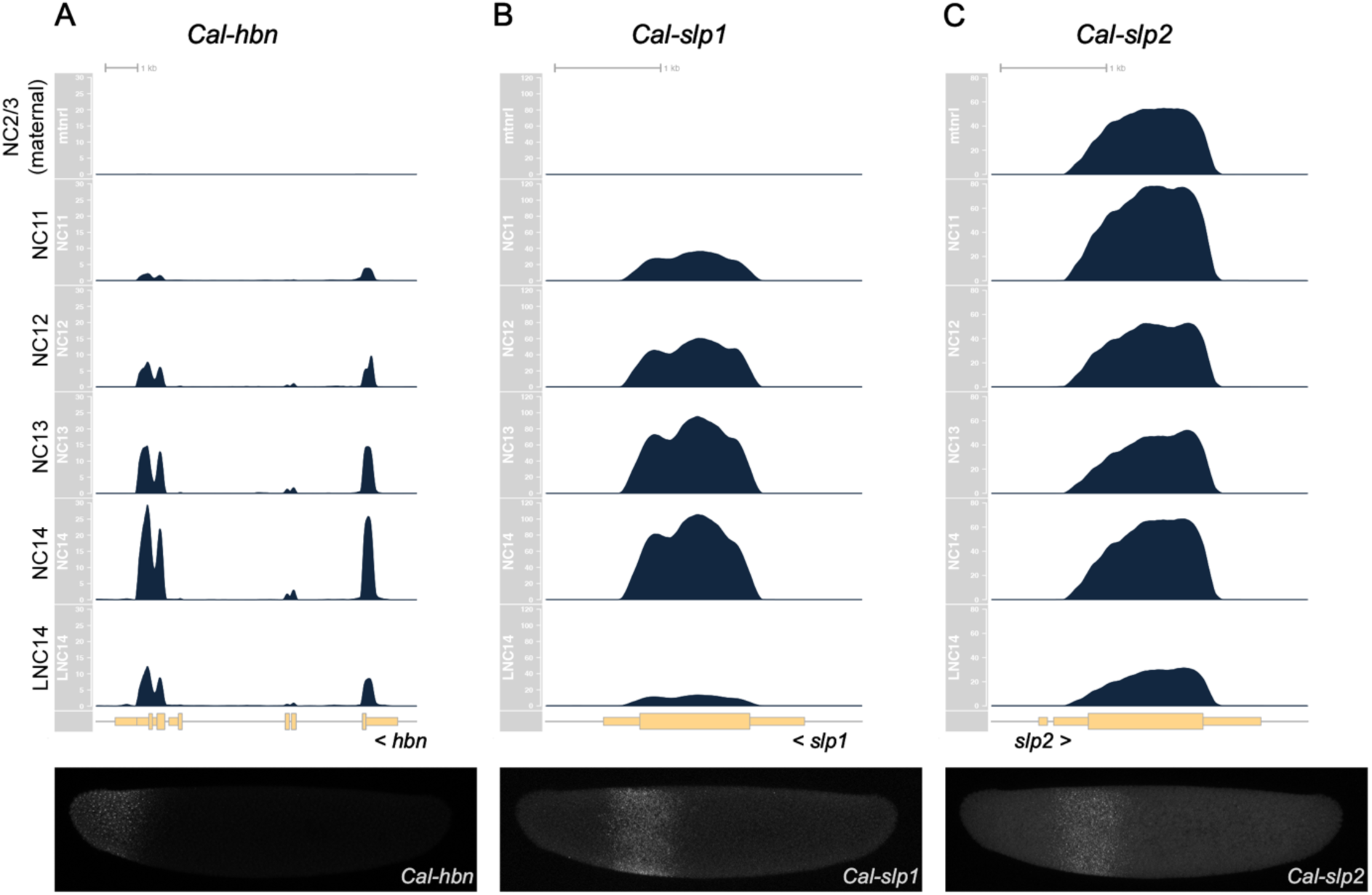
Time-course RNA-seq data for *Cal-hbn*, *Cal-slp1*, and *Cal-slp2*. Expression levels (CPM) of A) *Cal-hbn*, B) *Cal-slp1*, C) *Cal-slp2* at 6 consecutive stages, including NC2/3 (maternal, ∼45-minute old embryos), NC11, NC12, NC13, NC14 and Late NC14 are shown above fluorescent *in situ* hybridization chain reaction (HCR) staining of each gene at NC13. Anterior, left. Dorsal, up. Gene models are shown in pale yellow. The y-axis is the same for all tracks at each locus. *Cal*-*hbn* RNA-seq: 0-30 CPM. *Cal*-*slp1* RNA-seq: 0-120 CPM. *Cal*-*slp2* RNA-seq: 0-80 CPM.

## Discussion

### Dynamic chromatin states in the embryo of *Clogmia albipunctata*

In this study, we have established the resources for dissecting developmental gene networks of early Clogmia embryos by providing an annotated chromosome-level genome sequence for our inbred laboratory strain and by describing chromatin accessibility of precisely staged consecutive blastoderm stages. We have used these resources to examine how the actives of *zelda* and maternal *odd-paired* shape chromatin accessibility and gene expression in Clogmia. We found that, as in Drosophila, the period spanning zygotic genome activation is accompanied by large-scale changes in chromatin accessibility. However, there are notable differences between Clogmia and Drosophila in the early patterns of establishment and maintenance of accessible states before NC14. In Clogmia, the chromatin landscape is highly dynamic in that gains as well as losses occur during the nuclear cleavage cycles of the blastoderm stage. In Drosophila, only gains in accessibility are observed until widespread ZGA and lengthening of the cell cycle at NC14 (76, 78, 79, 81, 116). This difference could be a consequence of cell cycle timing. In Clogmia, NC11 through NC13 last ∼3x longer than in Drosophila (38), in which NC11 only lasts ∼10 minutes, NC12 ∼12 minutes, and NC13 ∼21 minutes (37). The much shorter cell cycle times of Drosophila present significant limitations to establish and maintain chromatin states necessary for transcriptional activity (81). Assuming that kinetic rates for nuclear import, DNA replication, and DNA binding of transcription factors is similar between Clogmia and Drosophila, the significantly longer syncytial interphase times in Clogmia could alleviate constraints on the initiation of zygotic transcription. However, the timing of AP pattering between the two species appears comparable (112, 117), suggesting that the timing of ZGA might also be conserved.

Alternatively, the longer syncytial interphase times in Clogmia could alleviate constraints on maintaining or silencing active chromatin states. In Drosophila, Polycomb group (PcG) proteins begin to establish heritable, transcriptionally silenced chromatin states during NC14 (118), but in Clogmia, the longer nuclear cleavage cycles may provide time for the maturation of repressive histone marks at accessible chromatin regions before NC14. This scenario would be consistent with the observed requirement of Cal-Zld for closing chromatin regions, as discussed in the next section.

### A unique role of Zld in regulating chromatin accessibility transitions in the Clogmia embryo

We demonstrate that reduced *Cal-zld* expression results in widespread losses of chromatin accessibility, consistent with a conserved role for Zld to pioneer chromatin accessibility within the context of zygotic genome activation. This study also describes a function of Cal-Zld in closing open chromatin, consistent with the unique chromatin dynamics of Clogmia. Specifically, many regions that lose accessibility during the syncytial cleavage divisions do so in a Zld-dependent manner. Based on the frequent occurrence of Zld motifs within this class of regions (53%), many of these effects are likely direct. We therefore propose that Cal-Zld functions not only to open and maintain chromatin accessibility but also to regulate the closure of many regions during nuclear cleavage cycles. This contrasts with Drosophila, in which Zld almost exclusively drives gains in chromatin accessibility during nuclear cleavage cycles (73, 78, 97, 119, 120). However, two recent studies found that during ZGA, Zld activity directs how repressive chromatin marks are deposited. Loss of *zld* results in context-specific losses and gains of H3K27me3 within specific PcG domains (118). Zld also enables histone acetylation at promoters and enhancers through its interaction with Chip, Drosophila’s homolog of LIM-domain-binding protein 1 (Ldb1), which in turn recruits the histone acetyltransferase CBP (121). The longer nuclear cleavage cycles in Clogmia might enable similar roles of Cal-Zld before NC14. In this case, Clogmia and Drosophila may differ in the timing of when specific histone marks are deposited and affected by Zld depletion. Differences in the activities of Zld and Cal-Zld could also reflect differences in their protein structures or isoforms. For example, Cal-Zld contains an additional zinc-finger that has been eroded in Drosophila (**S4 Figure**) (104). Additionally, stage-specific isoforms of Zld have been described in Drosophila, including a variant lacking the C-terminal three zinc-fingers (122–125). This isoform functions as a dominant negative form in cell culture although flies in which the expression of this isoform is suppressed are viable and fertile (124). Therefore, the molecular underpinnings of the distinct functional readouts of Zld and Cal-Zld could be complex and likely involve both direct and indirect mechanisms.

### Functional comparison of the ADs in Clogmia and Drosophila

Bcd contributes to the activation of many early segmentation genes; however, its role in the posterior embryo is much more subtle than in the anterior embryo, where all expression domains directly or indirectly depend on Bcd. Bcd target enhancers have been classified based on how their accessibility depends on Bcd and Zld (73). The accessibility of Bcd target enhancers that drive expression in the posterior of the embryo is established independent of Bcd (e.g., *Krüppel* CD1, *knirps* posterior, *giant* posterior). However, enhancers that open in a Bcd-dependent manner are associated with genes expressed in the anterior of the embryo (e.g., *orthodenticle*, *empty spiracles*, *buttonhead*, *giant*, *knirps*, *hunchback*, *sloppy-paired 1*, *fd19B*, *even-skipped*, *paired*, *cap’n’collar*, *huckebein, homeobrain*). Moreover, many correspond to enhancers that drive reporter gene expression in the anterior embryo including *giant* anterior (−6), *giant* anterior (−10), *knirps* anterior, *orthodenticle*, *buttonhead, even-skipped* stripe 1/5, *hunchback* P2, *hunchback* shadow enhancers, *eve-skipped* stripe 2, an early *paired* enhancer, *cap’n’collar*, and *huckebein* (reviewed in (73)). The majority of Bcd-dependent anterior expression domains first appear during NC13, with the noteworthy exception the P2 transcript of *hunchback*. The promoter of this transcript is in an open formation as early NC10 (81) and drives Bcd-dependent expression throughout the anterior half starting at NC11(47, 69, 126).

Clogmia’s anterior determinant is not required for *hunchback* expression before NC13 (**S16A Figure** and **S16B Figure**), and maternal *hunchback* activity does not contribute to embryo polarity given that knockdown of maternal *Cal-opa* transcripts in early embryos results in symmetrical double abdomens (3). However, Cal-Opa initiates symmetry-breaking at the level of chromatin accessibility and gene expression before NC12, notably at the loci of TF gene homologs of *sloppy-pai*red and *homeobrain*. These genes are expressed in complementary, evolutionarily conserved domains across the anterior half of the embryo. We note that these genes are among the Bcd targets discussed above; both loci contain Bcd-bound enhancers whose accessibility is sensitive to the concentration of Bcd. Mechanistically, Cal-Opa could prime accessibility at target loci through a canonical pioneering mechanism which requires binding of nucleosome occupied DNA and promoting nucleosome instability (49, 50, 58, 62, 127, 128). A vertebrate homolog of Opa, Zic3, was found to bind nucleosomal DNA *in vivo*, suggesting that Cal-Opa may pioneer chromatin directly (129). Alternatively, Cal-Opa binding may compete with nucleosomes to bind unoccupied DNA during each nuclear cleavage cycle (130), similar to Bcd (56, 73, 80).

Maternal Cal-Opa may target additional genes. Rigorously testing this possibility will require additional genetic tools or reagents to determine where maternal Cal-Opa binds *in vivo* and whether its target chromatin regions that become accessible independent of maternal Cal-Opa activity. We note that maternal Cal-Opa activity is expected to cease before the onset of zygotic *Cal-opa* expression during NC14. Bcd is still active at NC14, and because of this difference, the AD of Clogmia may target fewer genes than Bcd. We examined this possibility by individually analyzing our data on Clogmia orthologs of Bcd target genes other than *sloppy-paired* and *homeobrain*. The expression of Clogmia’s two *hunchback* orthologs (*Cal-hb1*, *Cal-hb2*) was reduced in *Cal-opa* RNAi embryos at NC13. We also observe a reduction in a few of the associated ATAC-seq peaks at these loci, but these regions did not include Opa motifs (**S16C Figure** and **S16D Figure**). Likewise, the expression of one of Clogmia’s two orthologs of *orthodenticle* (*Cal-otd1*) was reduced at NC13 in *Cal-opa* RNAi embryos, but this reduction was not associated with *Cal-opa* sensitive peaks containing Opa motifs (**S17A Figure**). The second ortholog (*Cal-otd2*) was not expressed at NC12 and NC13. In the case of Clogmia *buttonhead* (*Cal-btd*), we identified a *Cal-opa* sensitive peak with an Opa motif, but the observed reduction of *Cal-btd* expression was not significant (**S17B Figure**). *Cal-btd* expression could be under the control of AD-dependent and AD-independent enhancers, as previously reported for *orthodenticle* in Drosophila (131). Clogmia orthologs of *giant* (*Cal-gt*), *knirps* (*Cal-kni/knrl1, Cal-kni/knrl2*), and *empty spiracles* (*Cal-ems1-4*) were not expressed at NC12 or NC13, and none of these loci had significant changes in chromatin accessibility following knockdown of *Cal-opa*. The ortholog of *huckebein* (*Cal-hkb*) was not expressed at NC12 or NC13, either; however, its locus contained one Opa motif positive peak that lost accessibility following *Cal-opa* knockdown at NC13. The expression of Clogmia’s *cap’n’collar* (*Cal-cnc*) and *paired* (*Cal-prd*) was unaffected by *Cal-opa* RNAi, and the ATAC-seq peaks at their loci were largely *Cal-opa* insensitive. Only one small intronic peak within the Cal*-cnc* locus displayed reduced accessibility in *Cal-opa* RNAi embryos at NC13. The loci of *even-skipped* orthologs (*Cal-eve1, Cal-eve2*) contain ATAC-seq peaks with Opa motifs that were insensitive to *Cal-opa* RNAi, and the expression of *Cal-eve2* was elevated in *Cal-opa* RNAi embryos at NC13. Taken together, these observations point to significant differences in the target genes of Cal-Opa and Bcd, and suggest that *sloppy-paired* and *homeobrain* are key zygotic initiators of AP patterning – a role that is likely complemented by the distribution of maternal *Cal-slp2* transcript in an AP gradient (3).

### Role of *homeobrain* and *sloppy-paired* in breaking axial symmetry in insects

Neither *homeobrain* nor *sloppy-paired* has received much attention as target genes of Bcd. However, their identification in Clogmia as the only TF genes with strictly AD-dependent expression raises the question of whether they serve a more broadly conserved role in the initiation of anterior pattern formation. In Drosophila, *homeobrain* mutant embryos lack the labrum and anterior portions of the brain (132–135), indicating that *homeobrain* is essential for specifying the most anterior portion of the embryo. A more extreme *homeobrain* loss-of-function phenotype has been observed in the beetle *Tribolium castaneum*, where this gene plays an important zygotic role in establishing the embryo’s anterior-posterior polarity (136). In Tribolium, knockdown of *homeobrain* results in loss of the entire head and – in extreme cases – duplications of the tail end. The duplicated abdomen was defective, and thoracic structures persisted in these embryos. However, complete double abdomens could be obtained by simultaneously suppressing the expression *Tc-zen1*, thereby preventing the anterior repression of *caudal*, an important gene for posterior patterning (137, 138)

The *sloppy-paired* gene, a member of the *forkhead* domain gene family (139), is best known for its role as a pair-rule segmentation gene (21, 40), but it is initially expressed in a head domain of the syncytial Drosophila embryo, like its lineage-specific paralogs *sloppy-paired 2* and *fd19B* (138, 140–142). The partially redundant functions of *sloppy-paired* paralogs, which are common across insects (**Figure S2.14** and **Figure S2.15**), may have obscured a fundamental role of *sloppy-paired* activity in initiating anterior pattern formation in Drosophila. This also applies to the two *sloppy-paired* homologs of Tribolium, in which only the gnathocephalon (mandibular, maxillary, and labial segment) and even-numbered trunk segments are lost when *Tca-slp1* is knocked down (143, 144). We therefore propose that along with *homeobrain*, *sloppy-paired* may play a more widely conserved role in establishing the AP axis in insect embryos.

## Materials and Methods

### Clogmia culture and embryo collection

The Clogmia culture was maintained in 100mm x15mm Petri Dishes (Fisher Scientific FB0875712) on moist, compressed cotton (Jorvet SKU: J0197) sprinkled with parsley leaf powder (Starwest Botanicals SKU: 209895-5). For embryo collections, ∼2-day old females were transferred to round 250 mL glass bottles with a ∼1 inch bottom layer of moistened and compressed cotton and mated at room temperature for 2-3 days. Female ovaries were dissected with a pair of fine forceps (Dumont 5) and eggs released into water for egg activation and embryogenesis. For staged egg collections, embryos were released into flat bottom glass 100mm x15mm Petri Dishes (Corning CM: 101425255) containing 15ml of water. Care was taken to disperse the eggs on the bottom of the dish as crowding can delay development. Petri Dishes were covered with a lid and moved to an incubator set at 25° C until the desired stage. Egg harvesting post activation was on average 47 minutes for NC2/3, 261 minutes for NC11, 292 minutes for NC12, 332 minutes for NC13, 401 minutes for NC14, and 465 minutes for Late NC14.

### Embryo fixation

*Clogmia* Embryos were dechorionated using a 10% dilution of commercial bleach (8.25% sodium hypochloride) for 90 seconds. Dechorionated embryos were fixed in a 50 mL falcon tube, using 20 mL of boiling salt/detergent-solution (100 μL 10% triton-X; 500 μL 28% NaCl; up to 20 mL of water). The mixture was gently swirled for 10 seconds. To stop the heat fixation, 30 mL of deionized water was added. Embryos were transferred to 1.5 mL reaction vials and devitellinized in a 1:1 mixture of n-heptane and methanol by vigorous shaking. This step was followed by 3 washes with cold methanol (stored at –80C). Embryos were inspected under a stereomicroscope and manual devitellinizations were performed as needed using sharp tungsten needles in a 3% agar plate covered with methanol. Devitellinized embryos were stored in 100% methanol at −20°C.

### Microinjections of embryos and RNAi efficiency

*Clogmia albipunctata* embryo injection was done as previously described (3). We note here that RNAi efficiency in Clogmia embryos varies (3). This variability could be due to technical variables (e.g., injection needle, amount injected) or biological variables (e.g., differences in the RNAi response between individual embryos, or the general health or genetic background of the embryos or their mothers) or a combination of such effects. Therefore, we took care to control the age of the females and selected all experimental embryos from females with healthy looking ovaries and eggs.

To examine phenotypic effects of *Cal-zld* depletion, batches of ∼30 embryos were injected with *Cal-zld* dsRNA within the first hour of development (NC2/3). A subset of embryos from each batch (3 replicates) were monitored *in vivo* and allowed to develop until the end of NC14 (roughly 7-7.5 hours after injection). We also quantified knockdown efficiencies of the *Cal-zld* transcript in individual embryos by RT-qPCR and performed FC calculations of *Cal-zld* mRNA levels with the average level in uninjected control embryos as reference (**S4 Table**). In one quarter of the *Cal-zld* RNAi embryos (5/20), *Cal-zld* mRNA levels were reduced by ∼30-90%.

To assess maternal *Cal-opa* RNAi efficiency, we injected embryos in the first hour of development (NC2/3) with dsRNA of the isoform-specific first exon of the maternal *Cal-opa* isoform. In three independently conducted experiments we reproduced the double abdomen phenotype, confirming that our knockdown reagent was satisfactory. We also quantified knockdown efficiencies of the *Cal-opa* transcript in individual embryos by RT-qPCR and performed FC calculations of *Cal-opa* mRNA levels with the average level in uninjected control embryos as reference (**S5 Table**). This analysis suggests that *Cal-opa* RNAi can nearly abolish the *Cal-opa* transcript.

### Experimental setup of ATAC/RNA-seq experiments in *Cal-zld* and *Cal-opa* RNAi embryos

To assess *Cal-zld*’s role in shaping chromatin accessibility and zygotic transcription, we adapted our ATAC-seq protocol to isolate RNA from single-embryo ATAC-seq library preparations of Late NC14 *Cal-zld* RNAi embryos (8 replicates). In parallel, we processed stage-matched untreated embryos that were allowed to develop under water in the dish used for egg activation (wild-type controls; 5 replicates), and embryos that were prepared for injection under oil but not injected (alignment controls; 5 replicates). The isolated RNA was used to generate RNA-seq libraries to measure the efficiency of the *Cal-zld* knockdown as well as transcriptional effects of the knockdown in individual embryos. To identify individual injected embryos with strong *Cal-zld* knockdowns, we quantified *Cal-zld* expression levels using the RNA-seq data and chose embryos with knockdown efficiencies greater than 75% (n= 4, **S6 Figure** and see *Fold change calculations*).

To assess the impact of maternal Cal-Opa activity on chromatin accessibility and zygotic transcription, we performed ATAC-seq and RNA-seq on single *Cal-opa* RNAi embryos at NC12 (6 replicates) and NC13 (8 replicates). In parallel, we processed stage-matched untreated controls that were allowed to develop under water in the dish used for egg activation (wild-type controls; 2 replicates for NC12 and 3 replicates for NC13), as well as controls that were prepared for injection under oil but not injected (alignment controls; 4 replicates for each developmental stage), and embryos that were injected with dsRNA of the extraneous gene *DsRed* (injection controls; 3 replicates for each stage). To identify RNAi embryos with maximally reduced *Cal-opa* mRNA levels, we used the RNA-seq data to measure *Cal-opa* expression and selected embryos with knockdown efficiencies greater than 75% (**S10 Figure** and see *Fold change calculations*). We identified 4 embryos at NC12 and 5 embryos at NC13 with strong *Cal-opa* knockdowns and assessed changes in chromatin accessibility in these embryos by performing DESeq2 analysis with the alignment control group as reference.

### dsRNA synthesis

A PCR template was made using primers specific for the target gene with T7 overhangs and either cDNA, gDNA or a plasmid depending on the target sequence. The PCR product was run on a gel and purified using the Zymo Gel DNA Recovery Kit (D4007/D4008) following the manufactures instructions. The PCR template was used as the input for the transcription reaction producing dsRNA. dsRNA synthesis and purification were done using the Invitrogen MEGAscript RNAi Kit (AM1626) following the manufacturer’s instructions. Purified dsRNA was eluted in injection buffer (0.1 mM NaH_2_PO_4_ Ph 6.8; 5 mM KCl).

Forward and reverse primer sequences for dsRNA (lengths of dsRNAs in brackets; gene specific sequence of primers underlined):

*Cal-opa,* maternal transcript (201 bp):

5’-CAGAGATGCATAATACGACTCACTATAGGGAGAAAACAATTGTGAAGTGCGACA

5’-CAGAGATGCATAATACGACTCACTATAGGGAGA<u>CAAATTTCCAAACGATGACAGA</u>

*Cal-zld* (a combination of two dsRNAs were used 579 bp and 668 bp):

5’-TAATACGACTCACTATAGGGAGAAGTCCCGCAATTGATACAGC

5’-TAATACGACTCACTATAGGGAGAAGGATGTTGGTGGACACTCC

5’-TAATACGACTCACTATAGGGAGACGGAATGGTGTGTGAAACAG

5’-TAATACGACTCACTATAGGGAGATTCGAGGGGTTATTGTCCTG

dsRed (273 bp):

5’-CAGAGATGCATAATACGACTCACTATAGATGCAGAAGAAGACTATGG

5’-CAGAGATGCATAATACGACTCACTATAG<u>CTACAGGAACAGGTGGTG</u>

### RT-qPCR

RT-qPCR reactions were performed using the Luna® Universal One-Step RT-qPCR Kit (New England Biolabs Ipswich, MA).

Forward and reverse primer sequences for targets (lengths of amplicon in brackets):

*Cal-opa*, maternal transcript (112 bp):

5’-TTGCGCTTCATTCTGTCATCG

5’-CACTGAGGGCTGATTCTCGG

*Cal-zld* (138 bp):

5’-AAAGATGAACCACCGTGCGA

5’-CATTGATGGCAGTGGACCCT

*Cal-rpl35* (119 bp):

5’-CTGTCCAAAATCCGAGTCGT

5’-AGGTCGAGGGGCTTGTATTT

### Fluorescent HCR *in situ* hybridizations

Gene specific probes and amplifier-fluorophore sets were designed and synthesized by Molecular Instruments Inc (Los Angeles, CA). The amplifier-fluorophore set used was B1-647 for *Cal-slp1*, B2-546 for *Cal-hbn*, and B3-488 for *Cal-slp2*. Buffers (probe hybridization, probe wash and amplification buffers) throughout the protocol were purchased from Molecular Instruments Inc. The HCR *in situ* hybridizations were performed as described (145, 146). Briefly, fixed embryos were rehydrated and washed 3 times in PBS-Tween20 (1x PBS, 0.1% Tween-20). Embryos were permeabilized for 30 minutes at room temperature (100 ml Permeabilization buffer: 100 µL 100% Triton X-100, 50 µL Igepal CA-630, 50 mg Sodium Deoxycholate (Sigma D6750-25G), 50 mg Saponin (Sigma 47036-50G-F), 200 mg BSA Fraction V (Sigma A5611-5G). Following embryos were pre-hybridized in probe hybridization buffer for 30 minutes in a 37° C water bath. Once pre-hybridization was complete, fresh hybridization solution containing gene specific probes were added and the embryos were left to incubate overnight in a 37° C water bath. On the second day the hybridization solution was removed, and the embryos were washed 4 times with pre-warmed probe wash buffer followed by two washes in 5x SSCT (5x SSC, 0.1% Tween-20). An amplifier hairpin solution was prepared following the manufacturer’s instructions. Embryos were incubated with pre-warmed amplification buffer for 30 minutes at room temperature before adding amplifier hairpin solution. Embryos were left to incubate overnight at room temperature in the dark. The following day the embryos were washed 5 times in 5x SSCT at room temperature. The embryos were then counter stained with DAPI and mounted in Aqua-Polymount for imaging. Confocal imaging was performed on the Zeiss LSM 900.

### Image Acquisition and Processing

Confocal imaging was performed on the Zeiss LSM 900 using a Plan-Apochromat 20x/0.8 M27 objective. Pixel-density of 1024 x 1024. Excitation/Emission wavelengths (in nanometers) were: 493/517 for Alexa Flour 488, 653/668 for Alexa Flour 647, 401/422 for DAPI, and 548/561 for Alexa Flour 546. For each image a Z-stack of images was captured from the embryo’s surface to approximately midway through the embryo. Then a max projection was created from this Z-stack using FIJI/Image J. Brightness was increased in Photoshop. No further adjustments to the images were made.

### ATAC-seq library preparation

Single embryo ATAC-seq library preparations were done essential as described previously in *Drosophila melanogaster* (78, 81). Embryos developed in an incubator at 25°C until the desired nuclear cycle. Nuclear cycle was confirmed by time after egg activation and the morphology was checked under water using a transmission light microscope (Zeiss Stemi 2000-C). Once at the desired nuclear cycle individual embryos were placed in the lid of a 1.5 ml low-retention Eppendorf tube and homogenized in 10 μl of cold lysis buffer (10 mM Tris-HCl pH 7.4; 10 mM NaCl; 3 mM MgCl2; 1% Igepal CA-630) (147), using a fire-sealed microcapillary tube (Drummond Microcap 25). Once homogenized, 40 µl of lysis buffer was added to the lid, and the body of the tube was carefully closed over the lid. Samples were place on ice until all embryos were processed. The nuclei were pelleted by centrifugation at 800 RCF at 4°C for 10 minutes.

Supernatant removal was observed under a dissection scope to ensure that the nuclei pellet was not dislodged. The nuclear pellet was resuspended in 7.5 µl of Tagmentation buffer (Illumina Tagment DNA TDE1 Enzyme and Buffer Kit) and placed on ice until all samples were resuspended. Then 2.5 µl of Tn5 Transposase was added and mixed by pipetting up and down. Tagmentation and amplification of ATAC-seq libraries were performed as described previously (Buenrostro et al., 2015). The tagmentation reaction was performed at 37°C and 800 RPM for 30 minutes using an Eppendorf Thermomixer. Immediately after this step, the reactions were purified using the Qiagen Minelute kit following the manufacturer’s instructions. Purified tagmented DNA was eluted in 10 µl of buffer EB. Library amplification was performed as described previously (Buenrostro et al., 2015). Typically, 12-14 total PCR were performed.

Amplified libraries were purified using 1.8x Ampure SpRI beads following the manufacturer’s instructions. Purified ATAC-seq libraries were eluted in 15 ul. The ATAC-seq library profile was evaluated using Agilent high sensitivity bio-analyzer. Concentrations were estimated using a Qubit fluorometer. Libraries from the same experiment were pooled and sequenced together.

Sequencing was done at the University of Chicago Genomics Core (Chicago, IL, USA) using the Illumina NextSeq (PE 75 bp) and at Admera Health (South Plainfield, NJ, USA) using the Illumina HiSeq (PE 150 bp) and the Illumina NovaSeq X Plus (PE 150 bp).

ATAC-seq libraries were amplified using a set of modified Buenrostro primers that introduced Unique Dual Indexes (UDI) (78). The generalized UDI ATAC primer sequences are:

ATAC UDI Index Read 2 (i5):

5’-AATGATACGGCGACCACCGAGATCTACACnnnnnnnnTCGTCGGCAGCGTCAGATG T*G –3’

ATAC UDI Index Read 1 (i7):

5’-CAAGCAGAAGACGGCATACGAGATnnnnnnnnGTCTCGTGGGCTCGGAGATG*T –3’

Primers were synthesized with a terminal phosphorothioate bond (*) by IDT (Integrated DNA Technologies, Coralville, Iowa).

### RNA isolation for RT-qPCR and RNA-seq experiments

Embryos developed in an incubator at 25°C until the desired nuclear cycle. Nuclear cycle was confirmed by time after egg activation and the morphology was checked under water using a transmission light microscope (Zeiss Stemi 2000-C). RNA isolation was performed using the Zymo Quick-RNA Tissue/Insect Microprep Kit following the manufacturer’s instructions except for the tissue homogenization step. Instead, homogenization was done as follows. Once at the desired nuclear cycle individual embryos were placed in the lid of a 1.5 ml low-retention Eppendorf tube and homogenized in 10 μl of cold lysis buffer (10 mM Tris-HCl pH 7.4; 10 mM NaCl; 3 mM MgCl2; 1% Igepal CA-630) (147), using a fire-sealed microcapillary tube (Drummond Microcap 25). Once homogenized the lysate was added to 450 µl of Zymo RNA Lysis Buffer and thoroughly mixed. RNA isolation resumed following the manufacturer’s instructions including DNAse treatment. Purified RNA was eluted in DNase/RNase-Free Water.

### RNA-seq library preparation

RNA quality and quantity was assessed using the Agilent bio-analyzer. Non-Strand-specific RNA-seq libraries were prepared using the Smarter v4 Ultra-Low Input protocol from Takara for cDNA synthesis and Nextera XT from Illumina for Library prep (protocols provided by Takara and Illumina). Library quality and quantity was assessed using the Agilent bio-analyzer. Libraries from the same experiment were pooled and sequenced together. Sequencing was done at the University of Chicago Genomics Core (Chicago, IL, USA) using the Illumina NovaSeq X (PE 150 bp) and at Admera Health (South Plainfield, NJ, USA) using the Illumina NovaSeq X Plus (PE 150bp).

### ATAC-seq library preparation and RNA isolation

The first steps are the same as the standard ATAC-seq protocol above. Individual embryos are homogenized in cold lysis buffer, and a centrifugation step is performed to pellet the nuclei. Following the supernatant (∼47 ul) is removed and mixed thoroughly with 450 µl Zymo RNA lysis buffer in a fresh low-bind tube. The nuclear pellet is resuspended in 7.5 µl Tag mentation buffer and the ATAC-seq protocol continues as above with no deviations. RNA isolation proceeded immediately, or the RNA lysate was stored at temperately at –20°C. RNA isolation was performed using the Zymo Quick-RNA Tissue/Insect Microprep Kit following the manufacturer’s instructions. Isolated RNA was for RT-qPCR and RNA-seq experiments.

### Preparation of injected embryos for ATAC-seq /ATAC-seq and RNA isolation

Following the injection embryos were left to develop on the injection slide at 25°C. Once the embryos reached the desired nuclear stage any excess injection oil was removed using a pipette. The embryos were then washed 6 times using DI water. Embryos were removed from the injection slide using a blunted tungsten needle and transferred to the lid of a low bind tube. 10 μl of cold lysis buffer was added and the embryos were homogenized using a fire-sealed microcapillary tube. The ATAC-seq protocol proceeds as above.

### ATAC-seq data processing

Demultiplexed reads were trimmed of adapters using TrimGalore! (148) (version 0.6.10, cutadapt version 4.4) and mapped to the *Clogmia albipunctata* assembly (calbi-uni4000) using Bowtie2 (149) (version 2.4.1) with option –X 2000. Suspected optical and PCR duplicates were marked by Picard MarkDuplicates (version 2.21.4) using default parameters (https://broadinstitute.github.io/picard/). Mapped, trimmed, duplicate marked reads were imported into R using the GenomicAlignments (150) (version 1.38.2) and Rsamtools (151) (version 2.18.0). Reads that mapped, were properly paired, non-secondary and had a map quality scores ≥ 10 were imported. Only reads mapping to scaffolds 1-5 and scaffold 7 were used for downstream analysis. The start and end coordinates of the reads were adjusted to account for the Tn5 interaction site. Watson strand start sites had four base pairs subtracted, and Crick-strand start sites had five base pairs subtracted (147). Reads with mapped length ≤ 120 bp were considered to have originated from ‘open’ chromatin.

#### MACS2

Accessible chromatin peaks from NC11 through NC14 were determined using MACS2 (92) (version 2.2.7.1) with options –f BEDPE –q 1e-25 on a merged dataset comprising all ATAC reads corresponding to open chromatin (length ≤ 120 bp). Peaks were assigned to a genomic feature on the basis of overlapping with one or more features of a gene (e.g. exon, intron, promoter). Peaks that overlapped with no gene feature were classified as intergenic.

#### DESeq2

A count matrix for differential enrichment analyses using DESeq2 (96) (version 1.42.1) was generated by counting the number of open ATAC reads (length ≤ 120 bp) that overlapped with a MAC2 identified peak. Before running DESeq2, peaks with low count values were removed. DESeq2 design parameters were specific to the experimental data set. For the WT stage-wise comparisons (e.g. NC12 vs NC11, NC13 vs NC12, NC14 vs NC13 and LNC14 vs NC14) the design parameters passed to DESeq2 were stage and sequencing platform (∼instrument.atac.data + stage). For the comparison between RNAi embryos and alignment control embryos (e.g., *Cal-zld* RNAi vs alignment control and *Cal-opa* RNAi vs alignment control) the design components were embryo batch and genotype (∼Experiment.Date + genotype). All control groups were given their own genotype to account for differences in embryo manipulation, and the genotype for the RNAi embryos was determined by the strength of knockdown (see *Fold change calculations*). The criteria for a statistical significance change were a log_2_ Fold Change ≥ 1 and an adjusted p-value ≤ 0.05 or a log_2_ Fold Change ≤ –1 and an adjusted p-value ≤ 0.05.

Dynamic peaks were defined as peaks that underwent a statistically significant change in accessibility in at least one of the WT stage wise comparisons. Different dynamic peak classes were found using the R package DEGreport (152) (R package version 1.38.5) with the parameter of the minimum group size set to 10.

#### MEME

MEME (98) (version 5.4.1) analysis was performed using the following options –order 1 –meme-mod zoops –maxw 14 –meme-nmotifs 15 –meme-searchsize 100000 –centrimo-score 5.0 – centrimo-ethresh 10.0. Motifs were reported from all enrichment or discovery programs within the MEME suite (i.e., MEME, STREME (99), CentriMo), unless specified otherwise. MEME was given a set of sequences containing the 100 bp sequence centered on the summit of the peak. A set of background sequences were also given to MEME. To determine motifs enriched in dynamic peaks a set of non-dynamic peak sequences were provided as background sequences. To determine motifs enriched in *Cal*-*zld* sensitive peaks a set of sequences from peaks unaffected by *Cal*-*zld* were provided as background sequences. To determine motifs enriched in *Cal*-*opa* sensitive peaks a set of sequences from peaks unaffected by *Cal*-*opa* were provided as background sequences. Identified motifs were compared to known *Drosophila melanogaster* motifs from the following databases: OnTheFly_2014, Fly Factor Survey, FLYREG, iDMMPMM, and DMMPMM databases (111, 153–156).

### RNA-seq data processing

Demultiplexed reads were trimmed of adapters using TrimGalore! (148) (version 0.6.10, cutadapt version 4.4; https://www.bioinformatics.babraham.ac.uk/projects/trim_galore/) and mapped to the *Clogmia albipunctata* assembly (calbi-uni4000) using Bowtie2 (149) (version 2.4.1) with default parameters. Suspected optical and PCR duplicates were marked by Picard MarkDuplicates (version 2.21.4) using default parameters. Mapped, trimmed, duplicate marked reads were imported into R using the GenomicAlignments (150) (version 1.38.2) and Rsamtools (151) (version 2.18.0). Reads that mapped, were properly paired, non-secondary and had a map quality scores ≥ 10 were imported.

#### DESeq2

A gene count matrix for differential enrichment analyses was generated by counting the number of RNA-seq reads that overlapped with a transcript of a gene. Before running DESeq2, genes with low count values were removed. DESeq2 design parameters were specific to the experimental data set. For the comparison between RNAi embryos and alignment control embryos (e.g., *Cal-zld* RNAi vs alignment control and *Cal-opa* RNAi vs alignment control) the design components were embryo batch and genotype (∼Experiment.Date + genotype). All control groups were given their own genotype to account for differences in embryo manipulation, and the genotype for the RNAi embryos was determined by the strength of knockdown (see *Fold change calculations*). The criteria for a statistically significant change were a log_2_ Fold Change ≥ 1 and an adjusted p-value ≤ 0.05 or a log_2_ Fold Change ≤ –1 and an adjusted p-value ≤ 0.05.

#### Maternal, zygotic, maternal and zygotic classification

Gene expression was compared in NC2/3 (maternal, ∼45-minute old) and syncytial stage embryos (NC11-Late NC14). We assume here that NC2/3 embryos have little to no active transcription and instead are enriched in maternally deposited mRNAs. The distribution of gene expression was plotted for both stages, and a gene was determined to be expressed if its expression level was above 2^5 counts. Then genes were classified based on their expression profile in NC2/3 and syncytial stage embryos. We found 227 maternal genes, 691 zygotic genes, 6658 maternal and zygotic genes, and 7473 genes not expressed during either stage.

#### Fold change calculations

Fold change was calculated as the sample’s counts per million divided by the control’s counts per million for the gene of interest. First a counts per million (CPM) matrix was made by counting the number of RNA-seq reads that overlapped with a gene’s transcript and then normalizing to counts per million. Fold change calculations were done with embryos grouped by experimental date (i.e. embryos from the same egg packet). For each embryo the CPM was divided by the average CPM of wild-type and alignment control embryos. If a date did not have any wild-type or alignment control embryos, fold change was calculated using the average CPM of all wild-type and alignment control embryos in the data set. For *Cal-zld* RNAi experiments a fold change (FC) calculation of *Cal-zld* mRNA levels was conducted with the average of control embryos (untreated wild-type and alignment controls) from the same experimental date as reference. *Cal*-*zld* RNAi embryos with a FC < 0.25 (knockdown efficiency greater than 75%) were used for further analysis. Similarly, for *Cal-opa* RNAi experiments FC calculations were done separately for each stage. The fold change (FC) calculation of *Cal-opa* mRNA levels was conducted with the average of control embryos (untreated wild-type and alignment controls) from the same experimental date as reference. *Cal*-*opa* RNAi embryos with a FC < 0.25 (knockdown efficiency greater than 75%) were used for further analysis.

## Genome Assembly

We used Cantata Bio’s genome assembly services to generate a *de novo* reference genome for *Clogmia albipunctata* from ∼70 freshly eclosed males of a 12-generation inbred line of an established laboratory culture (157). High-fidelity (HiFi) PacBio reads were used to generate an initial draft assembly subsequently refined with Omni-C reads (**S1 Appendix**). Details on sequencing and software used to assemble the genome have been described elsewhere (88). We estimated genome heterozygosity and sequencing error rates using jellyfish to count 21-mers and GenomeScope2.0 to analyze the k-mer frequencies.

## Genome Annotation

We used *RepeatModeler* (158) (version v2.0.4) to identify repeat elements in the *Clogmia albipunctata* genome and *RepeatMasker*’s (159) (version v4.1.5) *fambd.py* script to extract *Arthropoda* records from the *Dfam* database. These sequences were combined to generate a custom repeat library, which was then used to soft-mask the genome. To predict gene structures, we used RNA-seq and protein evidence. mRNA data from 26 individual embryos of various stages, pooled samples of first and fourth-instar larvae, one female and one male pupa generated in this study were mapped to the genome (**S1 Table**). Additionally, 11 RNA samples (9 embryonic and two adult) from other studies were downloaded from NCBI (**S1 Table**). For protein evidence, we downloaded all available Dipteran sequences from NCBI’s RefSeq (January 2024) and UniProt’s complete protein sequences (Release 2023_04). A detailed description of the annotation pipeline is available elsewhere (88). Briefly, we assembled and mapped the RNA transcripts and protein sequences to the reference and used these data as evidence of gene structures and to predict gene models. *Evidence Modeler* (160) (EVM, version 2.1.0) was used to create a consensus gene set, which was further refined with Program to Assemble Spliced Alignments’ (PASA version 2.5.3) to add UTRs and identify alternative transcripts. Gene structures overlapping with repeats and transposable elements (TEs) were filtered out. Functional annotation was performed with *EggNOG-mapper* (161, 162) (version 2.1.12), assigning gene names based on orthology. We assessed the final annotation quality using BUSCO (Benchmarking Universal Single-Copy Orthologs) (163, 164) on the predicted protein sequences. Finally, we analyzed synteny between the *C. albipunctata* and *D. melanogaster* genomes using *MCScanX* (165) (primary release) with parameters: match_score = 50 (default), match size = 20 (minimum genes per syntenic block), gap_penalty = –1 (default), and max_gaps = 100. Results were visualized using the *circlize* R package (166) (version 4.1.2). Annotations of *Cal-eve1, Cal-eve2, Cal-tll1,* and *Cal-tll2* were manually corrected in the final output file (.gff3) based on RNA-seq data and transcript data available on NCBI.

## Transcription factor identification

Any gene in our annotation of the Clogmia genome that contained the Cluster-of-Orthologous-Gene (COG) symbol “K” for transcription or the Gene Ontology (GO) term “0003700” for DNA-binding transcription factor activity was treated as a putative transcription factor gene.

## Acknowledgments

We thank Lily Shiue and Qianyu Jin at Cantata Bio for assembling the genome, Melissa Harrison for the Cry2-zld mutant Drosophila strain, Natasha Megherea for managing the *Clogmia albipunctata* culture, and Suryadi Mcqueen for the illustrations of *Clogmia albipunctata* and *Drosophila melanogaster*. Edwin “Chip” Ferguson critically reviewed versions of this manuscript and provided valuable feedback. This work used Jetstream2 at Indiana University through allocation BIO220075 from the Advanced Cyberinfrastructure Coordination Ecosystem: Services & Support (ACCESS) program, which is supported by National Science Foundation grants #2138259, #2138286, #2138307, #2137603, and #2138296. Bioinformatic work was supported in part by the Notre Dame University Genomics and Bioinformatics Core Facility.

## Data and Resource Availability

<u>Genome</u>: The annotated *Clogmia albipunctata* genome can be found in NCBI’s Genome database under BioProject Accession PRJNA1165226.

<u>RNA-seq and ATAC-seq data:</u> All fastq files containing the raw sequencing reads for each sample have been uploaded to NCBI’s Sequence Read Archive (SRA) and can be found under the BioProject Accession PRJNA1198932. Individual BioSample and SRA accession numbers for the annotation of the genome are listed in S1 Table, including those samples that were not generated in this study but used as evidence for the annotation of the genome.

<u>Genome Browser:</u> The genomic data produced in this study can be further interrogated using a genome browser ecosystem centered on JBrowse2 (167) and delivered as a cloud image designed for individual use, currently available on NSF’s Jetstream2 cloud platform, with a portable Docker image with documentation and BSGenome package created for R available at Clogmia – github: https://github.com/kallistaconsulting/genomic_resources_clogmia. Jetstream2’s user-friendly interface (image: *Clogmia albipunctata* Genome Resources) (168, 169) integrates tools for genomic analysis, including BLAST search (SequenceServer2.0) (170), CRISPR guide RNA design (modified crisprDesigner) (171), R shiny-driven differential gene expression (DGE) analysis (freecount) (172), and synteny mapping for comparative genomics (ShinySyn) (173). Each tool is connected back to the browser, leveraging JBrowse2’s ability to dynamically visualize and contextualize the genomic data. BLAST results link directly to genomic regions, enabling users to explore expression tracks, variants, and synteny relationships. CRISPR tools provide sgRNA design with metrics for selection and visualization of target regions alongside expression data, off-target effects, and variants. DGE analysis visualizations are linked back to the genome browser for further exploration. All gene model annotations in the browser connect to common external databases, such as NCBI, Uniprot, FlyBase, and Google Scholar, for further exploring and interpreting the data. Future enhancements will include expanded sgRNA profiling, support for Docker to enable deployment on commercial cloud platforms, and further optimization of workflows, ensuring this ecosystem remains a powerful and accessible solution for the use of our genomic data presented here and elsewhere (88) for research and education. To access the *Clogmia albipunctata* genome browser and related tools, create an ACCESS ID and use this ID to create an ACCESS account to log in to Jetstream2 to apply for an ‘allocation’ of credits that can be used on Jetstream2. Detailed instructions on the use of Jetstream2 can be found at https://jetstream-cloud.org/get-started/index.html. This on-demand model reduces resource costs and security concerns compared to hosting a centralized site. Administrative tools, (e.g. automated reboot scripts, system service management, and load balancing for multi-user groups or workshops) enhance usability while minimizing operational overhead.

## Author Contributions

EA: Conceptualization, Methodology, Data curation, Investigation, Formal analysis, Visualization, Writing – original draft, and Writing – review and editing. ATT: Methodology, Formal analysis, Visualization, Writing – review and editing. ML: Data curation and Investigation. AJ: Formal analysis and Methodology. KK: Formal analysis and Visualization. SS: Project administration, Resources, Writing – review and editing. SB: Conceptualization, Resources, Funding acquisition, Project administration, Supervision, Writing – original draft, Writing – review and editing. USO: Conceptualization, Resources, Funding acquisition, Project administration, Supervision, Writing – original draft, Writing – review and editing.

