## Supplemental Data for "Breaking anterior-posterior symmetry in the moth fly *Clogmia albipunctata*"

**S1 Table: RNA-seq samples used for annotation.**

| SRA file ID | Experiment Title | Instrument | Submitter | Study Title | Sample Accessio | Stage | Sex |
|---|---|---|---|---|---|---|---|
| SRR7132659 | RNA-Seq of Clogmia albipunctata: cellular blastoderm stage | Illumina HiSeq 1000 | National Institute of Allergy and Infectious Diseases | Evolution of an Embryonic Axis Determinant via Alternative Transcription | SRS3270620 | Embryo, cellular blastoderm | Unknown |
| SRR7132660 | RNA-Seq of Clogmia albipunctata: early gastrula stage | Illumina HiSeq 1000 | National Institute of Allergy and Infectious Diseases | Evolution of an Embryonic Axis Determinant via Alternative Transcription | SRS3270619 | Embryo, early gastrulation | Unknown |
| SRR7132661 | RNA-Seq of Clogmia albipunctata: maternal stage | Illumina HiSeq 1000 | National Institute of Allergy and Infectious Diseases | Evolution of an Embryonic Axis Determinant via Alternative Transcription | SRS3270618 | Embryo, maternal | Unknown |
| SRR7132662 | RNA-Seq of Clogmia albipunctata: syncytial stage | Illumina HiSeq 1000 | National Institute of Allergy and Infectious Diseases | Evolution of an Embryonic Axis Determinant via Alternative Transcription | SRS3270616 | Embryo, syncytial blastoderm | Unknown |
| SRR7132663 | RNA-Seq of Clogmia albipunctata: posterior embryo maternal stage replicate 2 | Illumina HiSeq 1000 | National Institute of Allergy and Infectious Diseases | Evolution of an Embryonic Axis Determinant via Alternative Transcription | SRS3270615 | Embryo, maternal | Unknown |
| SRR7132664 | RNA-Seq of Clogmia albipunctata: anterior embryo maternal stage replicate 2 | Illumina HiSeq 1000 | National Institute of Allergy and Infectious Diseases | Evolution of an Embryonic Axis Determinant via Alternative Transcription | SRS3270617 | Embryo, maternal | Unknown |
| SRR7132665 | RNA-Seq of Clogmia albipunctata: posterior embryo maternal stage replicate 1 | Illumina HiSeq 1000 | National Institute of Allergy and Infectious Diseases | Evolution of an Embryonic Axis Determinant via Alternative Transcription | SRS3270614 | Embryo, maternal | Unknown |
| SRR7132666 | RNA-Seq of Clogmia albipunctata: anterior embryo maternal stage replicate 1 | Illumina HiSeq 1000 | National Institute of Allergy and Infectious Diseases | Evolution of an Embryonic Axis Determinant via Alternative Transcription | SRS3270613 | Embryo, maternal | Unknown |
| SRR5559337 | C_albipunctata_M_RNAseq | Illumina Genome Analyzer | Bachtrog Lab, UC Berkeley | Illumina sequencing of male and female transcriptomes of B.oleae, C.riparius, C.albipunctata, C.trivittatus, T.oleracea, C.fuscipes, C.patibulatus, M.abdita and S.bullata | SRS2199522 | Adult | Male |
| SRR5559338 | C_albipunctata_F_RNAseq | Illumina Genome Analyzer | Bachtrog Lab, UC Berkeley | Illumina sequencing of male and female transcriptomes of B.oleae, C.riparius, C.albipunctata, C.trivittatus, T.oleracea, C.fuscipes, C.patibulatus, M.abdita and S.bullata | SRS2199521 | Adult | Female |
| ERR160071 | RNA-Seq of Clogmia albipunctata embryos (8hrs, 10hrs, 12hrs at 25C) | Illumina HiSeq 2000 | CRG | Comparative transcriptomics of early dipteran development | ERS158633 | Unknown | Unknown |
| 23003FL-09-01-01 | Initiation of anterior pattern formation in moth fly embryos by Opa/Zic | Illumina NovaSeq X Plus | Schmidt-Ott Lab, University of Chicago | This study | SAMN45858185 | Embryo, maternal | Unknown |
| 23003FL-09-01-02 | Initiation of anterior pattern formation in moth fly embryos by Opa/Zic | Illumina NovaSeq X Plus | Schmidt-Ott Lab, University of Chicago | This study | SAMN45858186 | Embryo, maternal | Unknown |
| 23003FL-09-01-03 | Initiation of anterior pattern formation in moth fly embryos by Opa/Zic | Illumina NovaSeq X Plus | Schmidt-Ott Lab, University of Chicago | This study | SAMN45858187 | Embryo, maternal | Unknown |
| 23003FL-09-01-04 | Initiation of anterior pattern formation in moth fly embryos by Opa/Zic | Illumina NovaSeq X Plus | Schmidt-Ott Lab, University of Chicago | This study | SAMN45858188 | Embryo, maternal | Unknown |
| 23003FL-09-01-05 | Initiation of anterior pattern formation in moth fly embryos by Opa/Zic | Illumina NovaSeq X Plus | Schmidt-Ott Lab, University of Chicago | This study | SAMN45858189 | Embryo, NC11 | Unknown |
| 23003FL-09-01-06 | Initiation of anterior pattern formation in moth fly embryos by Opa/Zic | Illumina NovaSeq X Plus | Schmidt-Ott Lab, University of Chicago | This study | SAMN45858190 | Embryo, NC11 | Unknown |
| 23003FL-09-01-07 | Initiation of anterior pattern formation in moth fly embryos by Opa/Zic | Illumina NovaSeq X Plus | Schmidt-Ott Lab, University of Chicago | This study | SAMN45858191 | Embryo, NC11 | Unknown |
| 23003FL-09-01-08 | Initiation of anterior pattern formation in moth fly embryos by Opa/Zic | Illumina NovaSeq X Plus | Schmidt-Ott Lab, University of Chicago | This study | SAMN45858192 | Embryo, NC11 | Unknown |
| 23003FL-09-01-09 | Initiation of anterior pattern formation in moth fly embryos by Opa/Zic | Illumina NovaSeq X Plus | Schmidt-Ott Lab, University of Chicago | This study | SAMN45858196 | Embryo, NC12 | Unknown |
| 23003FL-09-01-10 | Initiation of anterior pattern formation in moth fly embryos by Opa/Zic | Illumina NovaSeq X Plus | Schmidt-Ott Lab, University of Chicago | This study | SAMN45858193 | Embryo, NC12 | Unknown |
| 23003FL-09-01-11 | Initiation of anterior pattern formation in moth fly embryos by Opa/Zic | Illumina NovaSeq X Plus | Schmidt-Ott Lab, University of Chicago | This study | SAMN45858194 | Embryo, NC12 | Unknown |
| 23003FL-09-01-12 | Initiation of anterior pattern formation in moth fly embryos by Opa/Zic | Illumina NovaSeq X Plus | Schmidt-Ott Lab, University of Chicago | This study | SAMN45858195 | Embryo, NC12 | Unknown |
| 23003FL-09-01-13 | Initiation of anterior pattern formation in moth fly embryos by Opa/Zic | Illumina NovaSeq X Plus | Schmidt-Ott Lab, University of Chicago | This study | SAMN45858197 | Embryo, NC13 | Unknown |
| 23003FL-09-01-14 | Initiation of anterior pattern formation in moth fly embryos by Opa/Zic | Illumina NovaSeq X Plus | Schmidt-Ott Lab, University of Chicago | This study | SAMN45858198 | Embryo, NC13 | Unknown |
| 23003FL-09-01-15 | Initiation of anterior pattern formation in moth fly embryos by Opa/Zic | Illumina NovaSeq X Plus | Schmidt-Ott Lab, University of Chicago | This study | SAMN45858199 | Embryo, NC13 | Unknown |
| 23003FL-09-01-16 | Initiation of anterior pattern formation in moth fly embryos by Opa/Zic | Illumina NovaSeq X Plus | Schmidt-Ott Lab, University of Chicago | This study | SAMN45858200 | Embryo, NC13 | Unknown |
| 23003FL-09-01-17 | Initiation of anterior pattern formation in moth fly embryos by Opa/Zic | Illumina NovaSeq X Plus | Schmidt-Ott Lab, University of Chicago | This study | SAMN45858201 | Embryo, NC14 | Unknown |
| 23003FL-09-01-18 | Initiation of anterior pattern formation in moth fly embryos by Opa/Zic | Illumina NovaSeq X Plus | Schmidt-Ott Lab, University of Chicago | This study | SAMN45858202 | Embryo, NC14 | Unknown |
| 23003FL-09-01-19 | Initiation of anterior pattern formation in moth fly embryos by Opa/Zic | Illumina NovaSeq X Plus | Schmidt-Ott Lab, University of Chicago | This study | SAMN45858203 | Embryo, NC14 | Unknown |
| 23003FL-09-01-20 | Initiation of anterior pattern formation in moth fly embryos by Opa/Zic | Illumina NovaSeq X Plus | Schmidt-Ott Lab, University of Chicago | This study | SAMN45858204 | Embryo, NC14 | Unknown |
| 23003FL-09-01-21 | Initiation of anterior pattern formation in moth fly embryos by Opa/Zic | Illumina NovaSeq X Plus | Schmidt-Ott Lab, University of Chicago | This study | SAMN45858205 | Embryo, Late NC14 | Unknown |
| 23003FL-09-01-22 | Initiation of anterior pattern formation in moth fly embryos by Opa/Zic | Illumina NovaSeq X Plus | Schmidt-Ott Lab, University of Chicago | This study | SAMN45858206 | Embryo, Late NC14 | Unknown |
| 23003FL-09-01-23 | Initiation of anterior pattern formation in moth fly embryos by Opa/Zic | Illumina NovaSeq X Plus | Schmidt-Ott Lab, University of Chicago | This study | SAMN45858207 | Embryo, Late NC14 | Unknown |
| 23003FL-09-01-24 | Initiation of anterior pattern formation in moth fly embryos by Opa/Zic | Illumina NovaSeq X Plus | Schmidt-Ott Lab, University of Chicago | This study | SAMN45858208 | Embryo, Late NC14 | Unknown |
| URS-EA-01 | Initiation of anterior pattern formation in moth fly embryos by Opa/Zic | Illumina NovaSeq 6000 | Schmidt-Ott Lab, University of Chicago | This study | SAMN45858209 | Embryos, Stage 13 | Female & Male |
| URS-EA-02 | Initiation of anterior pattern formation in moth fly embryos by Opa/Zic | Illumina NovaSeq 6000 | Schmidt-Ott Lab, University of Chicago | This study | SAMN45858210 | Embryos, Stage 15 | Female & Male |
| URS-EA-03 | Initiation of anterior pattern formation in moth fly embryos by Opa/Zic | Illumina NovaSeq 6000 | Schmidt-Ott Lab, University of Chicago | This study | SAMN45858211 | Larvae, L1 | Female & Male |
| URS-EA-04 | Initiation of anterior pattern formation in moth fly embryos by Opa/Zic | Illumina NovaSeq 6000 | Schmidt-Ott Lab, University of Chicago | This study | SAMN45858212 | Larvae, L4 | Female & Male |
| URS-EA-05 | Initiation of anterior pattern formation in moth fly embryos by Opa/Zic | Illumina NovaSeq 6000 | Schmidt-Ott Lab, University of Chicago | This study | SAMN45858213 | Pupa | Female |
| URS-EA-06 | Initiation of anterior pattern formation in moth fly embryos by Opa/Zic | Illumina NovaSeq 6000 | Schmidt-Ott Lab, University of Chicago | This study | SAMN45858214 | Pupa | Male |

**S2 Table: Homologs of early segmentation genes in *Clogmia albipunctata*.** Homologs were identified by reciprocal BLASTp (1) searches between *Drosophila melanogaster* and *Clogmia albipunctata*. Note that *knirps* and *knirps-like* as well as *sloppy-paired 1*, *sloppy-paired 2*, and *forkhead domain 19B* resulted from gene duplications specific to the Drosophila lineage, while *Cal-knrl1* and *Cal-knrl2* as well as *Cal-slp1*, *Cal-slp2* and *Cal-slp3* resulted from gene duplication specific to the Clogmia lineage.

| Gene name | Gene symbol | FlybaseID | Clogmia homolog | Clogmia gene symbol | Reference |
|---|---|---|---|---|---|
| *hunchback* | *hb* | FBgn0001180 | evm.TU.scaffold_7.1783 | *Cal-hb1* | AJ131041 (Rohr et al. 1999) |
| | | | evm.TU.scaffold_7.1785 | *Cal-hb2* | This study |
| *giant* | *gt* | FBgn0001150 | evm.TU.scaffold_1.1986 | *Cal-gt* | GU137318 (Garcia Solache et al. 2010) |
| *knirps* | *kni* | FBgn0001320 | None | N. A. | NA |
| *knirps-like* | *knrl* | FBgn0001323 | evm.TU.scaffold_7.729 | *Cal-kni/knrl1* | GU137321 (Garcia Solache et al. 2010) |
| | | | evm.TU.scaffold_7.724 | *Cal-kni/knrl2* | This study |
| *Krüppel* | *Kr* | FBgn0001325 | evm.TU.scaffold_7.1074 | *Cal-Kr* | GU137323 (Garcia Solache et al. 2010) and AJ131042 (Rohr et al. 1999) |
| *empty spiracles* | *ems* | FBgn0000576 | evm.TU.scaffold_4.1375 | *Cal-ems1* | This study |
| | | | evm.TU.scaffold_4.1371 | *Cal-ems2* | This study |
| | | | evm.TU.scaffold_4.1372 | *Cal-ems3* | This study |
| | | | evm.TU.scaffold_4.1374 | *Cal-ems4* | This study |
| *buttonhead* | *btd* | FBgn0000233 | evm.TU.scaffold_5.1944 | *Cal-btd* | This study |
| *orthodenticle/ocelliless* | *otd/oc* | FBgn0004102 | evm.TU.scaffold_5.235 | *Cal-otd1* | MN122108 (Yoon et al. 2019) |
| | | | evm.TU.scaffold_5.236 | *Cal-otd2* | This study |
| *huckebein* | *hkb* | FBgn0261434 | evm.TU.scaffold_2.1090 | *Cal-hkb* | GU137311 (Garcia Solache et al. 2010) |
| *tailless* | *tll* | FBgn0003720 | evm.TU.scaffold_3.2548a | *Cal-tll1* | GU137320 (Garcia Solache et al. 2010) |
| | | | evm.TU.scaffold_3.2548b | *Cal-tll2* | This study |
| *hairy* | *hry* | FBgn0001168 | evm.TU.scaffold_4.1097 | *Cal-hry* | AY645026 (Bullock et al. 2004) |
| *even-skipped* | *eve* | FBgn0000606 | evm.TU.scaffold_4.40a | *Cal-eve1* | AY645024 (Bullock et al. 2004) and GU137324 (Garcia Solache et al. 2010) |
| | | | evm.TU.scaffold_4.40b | *Cal-eve2* | AY645025 (Bullock et al. 2004) |
| *runt* | *run* | FBgn0003300 | evm.TU.scaffold_2.1058 | *Cal-run* | This study |
| *fushi tarazu* | *ftz* | FBgn0001077 | evm.TU.scaffold_3.2936 | *Cal-ftz* | This study |
| *odd-skipped* | *odd* | FBgn0002985 | evm.TU.scaffold_2.281 | *Cal-odd* | This study |
| *paired* | *prd* | FBgn0003145 | evm.TU.scaffold_2.1785 | *Cal-prd* | This study |
| *sloppy-paired 1* | *slp1* | FBgn0003430 | None | N. A. | N.A. |
| *sloppy-paired 2* | *slp2* | FBgn0004567 | evm.TU.scaffold_5.983 | *Cal-slp1* | This study |
| | | | evm.TU.scaffold_5.985 | *Cal-slp2* | MN122106 (Yoon et al. 2019) |
| | | | evm.TU.scaffold_5.987 | *Cal-slp3* | This study |
| *forkhead domain 19B* | *fd19B* | FBgn0031086 | None | N. A. | N. A. |
| *caudal* | *cad* | FBgn0000251 | evm.TU.scaffold_5.401 | *Cal-cad1* | Rivera-Pomar et al.1996 |
| | | | evm.TU.scaffold_5.403 | *Cal-cad2* | This study |
| *odd-paired* | *opa* | FBgn0003002 | evm.TU.scaffold_7.878 | *Cal-opa* | MN122104 (Yoon et al. 2019) |
| *Dichaete* | *D* | FBgn0000411 | evm.TU.scaffold_1.1497 | *Cal-D* | This study |
| *zerknüllt* | *zen* | FBgn0004053 | evm.TU.scaffold_7.614 | *Cal-zen* | AJ419659 (Stauber et al. 2002) |
| *homeobrain* | *hbn* | FBgn0008636 | evm.TU.scaffold_4.1647 | *Cal-hbn* | This study |

**S3 Table: Lists of transcription factor genes at NC12 and NC13 that are sensitive to *Cal-opa* knockdown.** 1) Differentially expressed transcription factor genes (DE TFs) containing Opa motif positive ATAC-seq peaks (potential direct Cal-Opa targets). 2) Differentially expressed transcription factor genes with *Cal-opa* sensitive ATAC-seq peaks. 3) Differentially expressed transcription factor genes with *Cal-opa* sensitive ATAC-seq peaks containing an Opa motif. Gene model name is listed in addition to a flybaseID if a Drosophila homolog was identified from a BLASTp search. Genes with an asterisk (*) are also identified in the *Cal-opa* RNAi versus injection control DESeq2 analysis.

| | | | | |
|---|---|---|---|---|
| **NC12** | | | | |
| DE TFs with Opa+ ATAC-seq peaks | evm.TU.scaffold_2.1802 | evm.TU.scaffold_3.1882(FBgn0039270) | evm.TU.scaffold_4.1647(Cal-hbn) * | evm.TU.scaffold_5.1243(FBgn0031573) |
| | evm.TU.scaffold_5.150(FBgn0035160) | evm.TU.scaffold_5.184(FBgn0013753) | evm.TU.scaffold_5.983(Cal-slp1) * | evm.TU.scaffold_5.985(Cal-slp2) * |
| | evm.TU.scaffold_7.2949(FBgn0037491) | evm.TU.scaffold_7.878(Cal-opa) * | | |
| DE TFs with *Cal-opa* sensitive ATAC-seq peaks | evm.TU.scaffold_4.1647(Cal-hbn) * | evm.TU.scaffold_5.983(Cal-slp1) * | evm.TU.scaffold_5.985(Cal-slp2) | evm.TU.scaffold_7.878(Cal-opa) * |
| DE TFs with Opa+ *Cal-opa* sensitive ATAC-seq peaks | evm.TU.scaffold_4.1647(Cal-hbn) * | evm.TU.scaffold_5.983(Cal-slp1) * | evm.TU.scaffold_5.985(Cal-slp2) | evm.TU.scaffold_7.878(Cal-opa) * |
| **NC13** | | | | |
| DE TFs with Opa+ ATAC-seq peaks | evm.TU.scaffold_1.1194(FBgn0002735) | evm.TU.scaffold_1.1262 | evm.TU.scaffold_1.134 | evm.TU.scaffold_1.1422 |
| | evm.TU.scaffold_1.1497(Cal-D) | evm.TU.scaffold_1.1676(FBgn0010417) | evm.TU.scaffold_1.182(FBgn0043002) | evm.TU.scaffold_1.2409(FBgn0011715) |
| | evm.TU.scaffold_1.2511(FBgn0005427) | evm.TU.scaffold_1.2651 | evm.TU.scaffold_1.2674(FBgn0263757) | evm.TU.scaffold_1.2960(FBgn0086655) |
| | evm.TU.scaffold_1.348 | evm.TU.scaffold_1.824(FBgn0002573) | evm.TU.scaffold_2.1031(FBgn0010422) | evm.TU.scaffold_2.1246(FBgn0003687) |
| | evm.TU.scaffold_2.1413(FBgn0031776) | evm.TU.scaffold_2.1535(FBgn0037363) | evm.TU.scaffold_2.1830 | evm.TU.scaffold_2.2083 |
| | evm.TU.scaffold_2.250(FBgn0266557) | evm.TU.scaffold_2.275(FBgn0028398) | evm.TU.scaffold_2.751(FBgn0004606) | evm.TU.scaffold_2.825 |
| | evm.TU.scaffold_2.935(FBgn0032016) | evm.TU.scaffold_3.1271(FBgn0038499) | evm.TU.scaffold_3.1502(FBgn0004897) | evm.TU.scaffold_3.1619(FBgn0003612) |
| | evm.TU.scaffold_3.1882(FBgn0039270) | evm.TU.scaffold_3.2548b(Cal-tll2) | evm.TU.scaffold_3.2623(FBgn0001235) | evm.TU.scaffold_3.2809(FBgn0262127) |
| | evm.TU.scaffold_3.514(FBgn0038903) | evm.TU.scaffold_3.913 | evm.TU.scaffold_4.1248 | evm.TU.scaffold_4.1328 |
| | evm.TU.scaffold_4.1461(FBgn0033540) | evm.TU.scaffold_4.1647(Cal-hbn) * | evm.TU.scaffold_4.1673(FBgn0042085) | evm.TU.scaffold_4.1700(FBgn0032329) |
| | evm.TU.scaffold_4.1897 | evm.TU.scaffold_4.2102(FBgn0031822) | evm.TU.scaffold_4.2428 | evm.TU.scaffold_4.40b(Cal-eve2) |
| | evm.TU.scaffold_4.667(FBgn0261934) | evm.TU.scaffold_4.874(FBgn0027549) | evm.TU.scaffold_4.881(FBgn0014143) | evm.TU.scaffold_4.918(FBgn0040273) |
| | evm.TU.scaffold_4.958(FBgn0003415) | evm.TU.scaffold_5.1243(FBgn0031573) | evm.TU.scaffold_5.1294(FBgn0261617) | evm.TU.scaffold_5.1512(FBgn0035145) |
| | evm.TU.scaffold_5.175(FBgn0004859) | evm.TU.scaffold_5.1857(FBgn0036581) | evm.TU.scaffold_5.2222(FBgn0002931) | evm.TU.scaffold_5.235(Cal-otd1) |
| | evm.TU.scaffold_5.842(FBgn0086447) | evm.TU.scaffold_5.983(Cal-slp1) * | evm.TU.scaffold_5.985(Cal-slp2) * | evm.TU.scaffold_7.1036(FBgn0024841) |
| | evm.TU.scaffold_7.1074(Cal-Kr) | evm.TU.scaffold_7.1127(FBgn0014037) | evm.TU.scaffold_7.1185(FBgn0037085) | evm.TU.scaffold_7.1252(FBgn0023518) |
| | evm.TU.scaffold_7.1663(FBgn0038306) | evm.TU.scaffold_7.1925(FBgn0027529) | evm.TU.scaffold_7.228 | evm.TU.scaffold_7.2949(FBgn0037491) |
| | evm.TU.scaffold_7.53(FBgn0004396) * | evm.TU.scaffold_7.614(Cal-zen) | evm.TU.scaffold_7.878(Cal-opa) | evm.TU.scaffold_7.993(FBgn0051950) |
| DE TFs with *Cal-opa* sensitive ATAC-seq peaks | evm.TU.scaffold_1.1497(Cal-D) | evm.TU.scaffold_1.2511(FBgn0005427) | evm.TU.scaffold_3.2623(FBgn0001235) | evm.TU.scaffold_4.1328 |
| | evm.TU.scaffold_4.1449(FBgn0037834) | evm.TU.scaffold_4.1647(Cal-hbn) * | evm.TU.scaffold_4.2497 | evm.TU.scaffold_5.235(Cal-otd1) |
| | evm.TU.scaffold_5.983(Cal-slp1) * | evm.TU.scaffold_5.985(Cal-slp2) * | evm.TU.scaffold_5.987(Cal-slp3) | evm.TU.scaffold_7.1074(Cal-Kr) |
| | evm.TU.scaffold_7.1783(Cal-hb1) | evm.TU.scaffold_7.1785(Cal-hb2) * | evm.TU.scaffold_7.878(Cal-opa) | |
| DE TFs with Opa+ *Cal-opa* sensitive ATAC-seq peaks | evm.TU.scaffold_1.1497(Cal-D) | evm.TU.scaffold_4.1328 | evm.TU.scaffold_4.1647(Cal-hbn) * | evm.TU.scaffold_5.983(Cal-slp1) * |
| | evm.TU.scaffold_5.985(Cal-slp2) * | evm.TU.scaffold_7.1074(Cal-Kr) | evm.TU.scaffold_7.878(Cal-opa) | |

**S4 Table: *Cal-zld* RT-qPCR data.** Fold change calculated using the ΔΔCt method. Reference gene *Cal-rpl35*. Target gene *Cal-zld*. Percent knockdown was calculated as 100-(FC*100).

| Treatment | Embryo No | Fold change | Percent knockdown |
|---|---|---|---|
| Alignment control | 1 | 1.04427378 | -4.4273782 |
| Alignment control | 2 | 0.95760328 | 4.23967193 |
| *Cal-zld* RNAi | 3 | 1.92919637 | -92.919637 |
| *Cal-zld* RNAi | 4 | 1.1011416 | -10.11416 |
| *Cal-zld* RNAi | 5 | 1.61440215 | -61.440215 |
| *Cal-zld* RNAi | 6 | 0.88300897 | 11.6991029 |
| *Cal-zld* RNAi | 7 | 1.30450276 | -30.450276 |
| *Cal-zld* RNAi | 8 | 0.82188019 | 17.8119813 |
| Alignment control | 9 | 0.8630404 | 13.69596 |
| Alignment control | 10 | 1.15869431 | -15.869431 |
| *Cal-zld* RNAi | 11 | 2.17723933 | -117.72393 |
| *Cal-zld* RNAi | 12 | 0.4735209 | 52.6479103 |
| *Cal-zld* RNAi | 13 | 0.06091748 | 93.9082523 |
| *Cal-zld* RNAi | 14 | 1.63580412 | -63.580412 |
| *Cal-zld* RNAi | 15 | 2.08132174 | -108.13217 |
| *Cal-zld* RNAi | 16 | 0.68349373 | 31.6506274 |
| Alignment control | 17 | 0.98043915 | 1.95608492 |
| Alignment control | 18 | 1.01995111 | -1.995111 |
| *Cal-zld* RNAi | 19 | 1.11457987 | -11.457987 |
| *Cal-zld* RNAi | 20 | 0.51245598 | 48.7544016 |
| *Cal-zld* RNAi | 21 | 1.1045816 | -10.45816 |
| *Cal-zld* RNAi | 22 | 1.27059126 | -27.059126 |
| *Cal-zld* RNAi | 23 | 5.13014641 | -413.01464 |
| *Cal-zld* RNAi | 24 | 0.5913155 | 40.86845 |
| *Cal-zld* RNAi | 25 | 1.80562713 | -80.562713 |
| *Cal-zld* RNAi | 26 | 0.96392981 | 3.60701923 |

**S5 Table: *Cal-opa* RT-qPCR data.** Fold change calculated using the $\Delta\Delta$Ct method. Reference gene *Cal-rpl35*. Target gene *Cal-opa*. Percent knockdown was calculated as 100-(FC*100). Indicated is whether the RNA was isolated immediately or in parallel with an ATAC-seq library preparation. RNA was either isolated using the Zymo Quick-RNA Tissue/Insect Microprep Kit following the manufacturer's instructions kit or following an ATAC-seq library preparation using the protocol by Li et al. (2). Briefly following the tagmentation reaction, mRNA was isolated from the supernatant using Dynabeads® Oligo (dT)25 beads. Following washes were performed and mRNA was eluted directly from the beads.

| Treatment | Embryo No | Fold change | Percent knockdown | RNA isolation Method |
|---|---|---|---|---|
| Alignment control | 1 | 0.90093817 | 9.906183288 | Standard RNA isolation |
| *Cal-opa* RNAi | 2 | 0.73739012 | 26.26098757 | Standard RNA isolation |
| Injection control | 3 | 0.90469287 | 9.530713411 | RNA isolated from ATAC-seq library prep |
| Alignment control | 4 | 1.10995409 | -10.99540862 | Standard RNA isolation |
| *Cal-opa* RNAi | 5 | 0.54961756 | 45.03824413 | Standard RNA isolation |
| *Cal-opa* RNAi | 6 | 0.97857206 | 2.142793791 | Standard RNA isolation |
| *Cal-opa* RNAi | 7 | 1.08729996 | -8.72999598 | Standard RNA isolation |
| Injection control | 8 | 0.704416 | 29.55838 | RNA isolated from ATAC-seq library prep |
| Injection control | 9 | 0.661356 | 33.86435 | RNA isolated from ATAC-seq library prep |
| Alignment control | 10 | 1.0085272 | -0.85272042 | Standard RNA isolation |
| Alignment control | 11 | 0.99154489 | 0.845510579 | Standard RNA isolation |
| *Cal-opa* RNAi | 12 | 5.13815353 | -413.8153533 | Standard RNA isolation |
| *Cal-opa* RNAi | 13 | 0.57345413 | 42.65458685 | Standard RNA isolation |
| *Cal-opa* RNAi | 14 | 0.18884179 | 81.11582059 | Standard RNA isolation |
| *Cal-opa* RNAi | 15 | 2.14094836 | -114.0948363 | Standard RNA isolation |
| Injection control | 16 | 1.09619161 | -9.619161168 | RNA isolated from ATAC-seq library prep |
| Injection control | 17 | 0.56487361 | 43.51263927 | RNA isolated from ATAC-seq library prep |
| Alignment control | 18 | 0.96845036 | 3.154964497 | Standard RNA isolation |
| *Cal-opa* RNAi | 19 | 2.07663836 | -107.6638364 | Standard RNA isolation |
| *Cal-opa* RNAi | 20 | 0.63650776 | 36.34922435 | Standard RNA isolation |
| *Cal-opa* RNAi | 21 | 1.95782254 | -95.78225362 | Standard RNA isolation |
| *Cal-opa* RNAi | 22 | 1.23306598 | -23.3065981 | Standard RNA isolation |
| Injection control | 23 | 1.30359886 | -30.3598864 | RNA isolated from ATAC-seq library prep |
| Injection control | 24 | 0.8411879 | 15.88121017 | RNA isolated from ATAC-seq library prep |
| Alignment control | 25 | 1.03257745 | -3.257745202 | Standard RNA isolation |
| Alignment control | 26 | 0.73866903 | 26.13309682 | Standard RNA isolation |
| *Cal-opa* RNAi | 27 | 0.07282108 | 92.71789174 | Standard RNA isolation |
| *Cal-opa* RNAi | 28 | 0.1320357 | 86.79642968 | Standard RNA isolation |
| *Cal-opa* RNAi | 29 | 0.25578437 | 74.42156326 | Standard RNA isolation |
| *Cal-opa* RNAi | 30 | 0.03349292 | 96.65070793 | Standard RNA isolation |
| *Cal-opa* RNAi | 31 | 0.19895306 | 80.10469391 | Standard RNA isolation |
| *Cal-opa* RNAi | 32 | 0.63683874 | 36.31612623 | Standard RNA isolation |
| Injection control | 33 | 2.2137618 | -121.3761805 | Standard RNA isolation |
| Injection control | 34 | 1.93790789 | -93.79078898 | Standard RNA isolation |
| Injection control | 35 | 2.34323232 | -134.3232324 | Standard RNA isolation |
| Alignment control | 36 | 1.35378628 | -35.37862791 | Standard RNA isolation |
| Alignment control | 37 | 0.12904982 | 87.09501831 | Standard RNA isolation |
| Alignment control | 38 | 0.84177117 | 15.82288326 | Standard RNA isolation |
| *Cal-opa* RNAi | 39 | 0.23882073 | 76.11792659 | Standard RNA isolation |
| *Cal-opa* RNAi | 40 | 0.26370558 | 73.62944207 | Standard RNA isolation |
| *Cal-opa* RNAi | 41 | 0.18313761 | 81.68623907 | Standard RNA isolation |
| *Cal-opa* RNAi | 42 | 0.79802177 | 20.19782348 | Standard RNA isolation |
| *Cal-opa* RNAi | 43 | 0.30418059 | 69.58194103 | Standard RNA isolation |
| Injection control | 44 | 1.77727468 | -77.72746772 | Standard RNA isolation |
| Injection control | 45 | 1.90087896 | -90.08789554 | Standard RNA isolation |
| Injection control | 46 | 1.22235782 | -22.23578204 | Standard RNA isolation |
| *Cal-opa* RNAi | 47 | 0.99343673 | 0.656326528 | Standard RNA isolation |
| *Cal-opa* RNAi | 48 | 0.32522273 | 67.47772693 | Standard RNA isolation |
| Alignment control | 49 | 0.99584975 | 0.415024691 | Standard RNA isolation |
| Alignment control | 50 | 1.19292224 | -19.2922236 | Standard RNA isolation |
| *Cal-opa* RNAi | 51 | 0.118517 | 88.14829 | Standard RNA isolation |
| *Cal-opa* RNAi | 52 | 0.88403 | 11.59703 | Standard RNA isolation |
| Alignment control | 53 | 0.952418 | 4.758208 | Standard RNA isolation |
| Alignment control | 54 | 0.989428 | 1.057198 | Standard RNA isolation |
| *Cal-opa* RNAi | 55 | 0.007323 | 99.26769 | Standard RNA isolation |
| *Cal-opa* RNAi | 56 | 0.028206 | 97.17936 | Standard RNA isolation |
| *Cal-opa* RNAi | 57 | 5.637934 | -463.793 | Standard RNA isolation |
| *Cal-opa* RNAi | 58 | 0.20537512 | 79.46248786 | Standard RNA isolation |
| *Cal-opa* RNAi | 59 | 0.13395621 | 86.60437933 | Standard RNA isolation |
| *Cal-opa* RNAi | 60 | 0.52443408 | 47.55659247 | Standard RNA isolation |
| *Cal-opa* RNAi | 61 | 0.43719031 | 56.28096891 | Standard RNA isolation |
| *Cal-opa* RNAi | 62 | 0.6726059 | 32.73941027 | Standard RNA isolation |
| Alignment control | 63 | 1.061178 | -6.1178 | Standard RNA isolation |
| Alignment control | 64 | 0.81356715 | 18.64328497 | Standard RNA isolation |
| *Cal-opa* RNAi | 65 | 0.03331926 | 96.66807439 | RNA isolated from ATAC-seq library prep |
| *Cal-opa* RNAi | 66 | 0.001725 | 99.82754 | RNA isolated from ATAC-seq library prep |
| *Cal-opa* RNAi | 67 | 0.002495 | 99.75054 | RNA isolated from ATAC-seq library prep |
| *Cal-opa* RNAi | 68 | 0.758384 | 24.16162 | RNA isolated from ATAC-seq library prep |
| *Cal-opa* RNAi | 69 | 0.0045529 | 99.54471047 | RNA isolated from ATAC-seq library prep |
| *Cal-opa* RNAi | 70 | 0.83393104 | 16.60689562 | RNA isolated from ATAC-seq library prep |
| *Cal-opa* RNAi | 71 | 0.85916075 | 14.08392451 | RNA isolated from ATAC-seq library prep |
| *Cal-opa* RNAi | 72 | 0.00488306 | 99.51169441 | RNA isolated from ATAC-seq library prep |
| *Cal-opa* RNAi | 73 | 1.63240609 | -63.24060923 | RNA isolated from ATAC-seq library prep |
| Alignment control | 74 | 0.90114635 | 9.885364791 | Standard RNA isolation |
| Alignment control | 75 | 1.40915795 | -40.91579534 | Standard RNA isolation |
| Alignment control | 76 | 1 | 0 | RNA isolated from ATAC-seq library prep |
| Alignment control | 77 | 1 | 0 | RNA isolated from ATAC-seq library prep |
| Alignment control | 78 | 0.9862327 | 1.376729551 | RNA isolated from ATAC-seq library prep |
| Alignment control | 79 | 1.01395948 | -1.395947979 | RNA isolated from ATAC-seq library prep |
| *Cal-opa* RNAi | 80 | 0.78024548 | 21.97545198 | RNA isolated from ATAC-seq library prep |
| *Cal-opa* RNAi | 81 | 0.00900848 | 99.09915164 | RNA isolated from ATAC-seq library prep |
| *Cal-opa* RNAi | 82 | 0.00221036 | 99.77896355 | RNA isolated from ATAC-seq library prep |
| *Cal-opa* RNAi | 83 | 11.6924066 | -1069.240661 | Standard RNA isolation |
| Injection control | 84 | 0.882703 | 11.72970037 | Standard RNA isolation |
| Alignment control | 85 | 1 | 0 | RNA isolated from ATAC-seq library prep |
| Alignment control | 86 | 0.67618973 | 32.38102746 | Standard RNA isolation |
| Alignment control | 87 | 1.47887488 | -47.88748814 | Standard RNA isolation |
| *Cal-opa* RNAi | 88 | 1.40785622 | -40.78562166 | Standard RNA isolation |
| *Cal-opa* RNAi | 89 | 0.3023937 | 69.76062975 | Standard RNA isolation |
| *Cal-opa* RNAi | 90 | 0.70686176 | 29.31382409 | Standard RNA isolation |
| *Cal-opa* RNAi | 91 | 0.25836812 | 74.1631875 | Standard RNA isolation |
| *Cal-opa* RNAi | 92 | 0.64104609 | 35.89539068 | Standard RNA isolation |
| Injection control | 93 | 1.490883 | -49.0883 | Standard RNA isolation |
| Injection control | 94 | 1.85339 | -85.339 | Standard RNA isolation |
| Injection control | 95 | 2.033784 | -103.378 | Standard RNA isolation |
| Alignment control | 96 | 0.86314011 | 13.68598922 | Standard RNA isolation |
| Alignment control | 97 | 1.58190906 | -58.19090615 | Standard RNA isolation |
| Alignment control | 98 | 0.7323812 | 26.76187982 | Standard RNA isolation |
| *Cal-opa* RNAi | 99 | 1.43561278 | -43.56127753 | Standard RNA isolation |
| *Cal-opa* RNAi | 100 | 0.24496825 | 75.50317524 | Standard RNA isolation |
| *Cal-opa* RNAi | 101 | 1.04367076 | -4.367076041 | Standard RNA isolation |
| *Cal-opa* RNAi | 102 | 1.46984769 | -46.98476902 | Standard RNA isolation |
| *Cal-opa* RNAi | 103 | 1.28847689 | -28.84768947 | Standard RNA isolation |
| Injection control | 104 | 1.5613926 | -56.13925956 | Standard RNA isolation |
| Injection control | 105 | 1.639588 | -63.95879968 | Standard RNA isolation |
| Injection control | 106 | 2.34458622 | -134.4586219 | Standard RNA isolation |
| Injection control | 107 | 0.99711605 | 0.288394667 | Standard RNA isolation |

### Supplemental Figures and Figure Legends

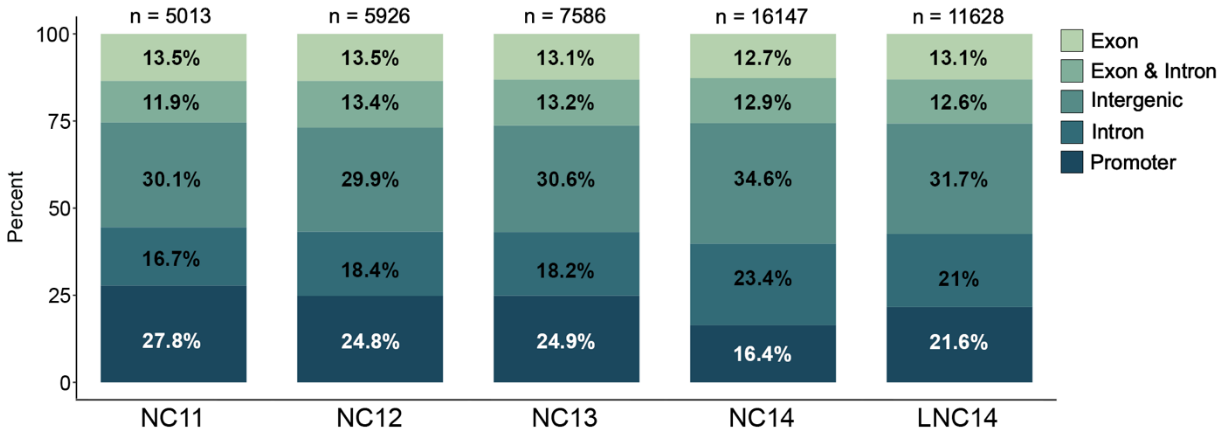


**S1 Figure, relating to Table 3: Genomic distribution of ATAC-seq peaks at nuclear cycles NC11, NC12, NC13, NC14 and Late NC14.**

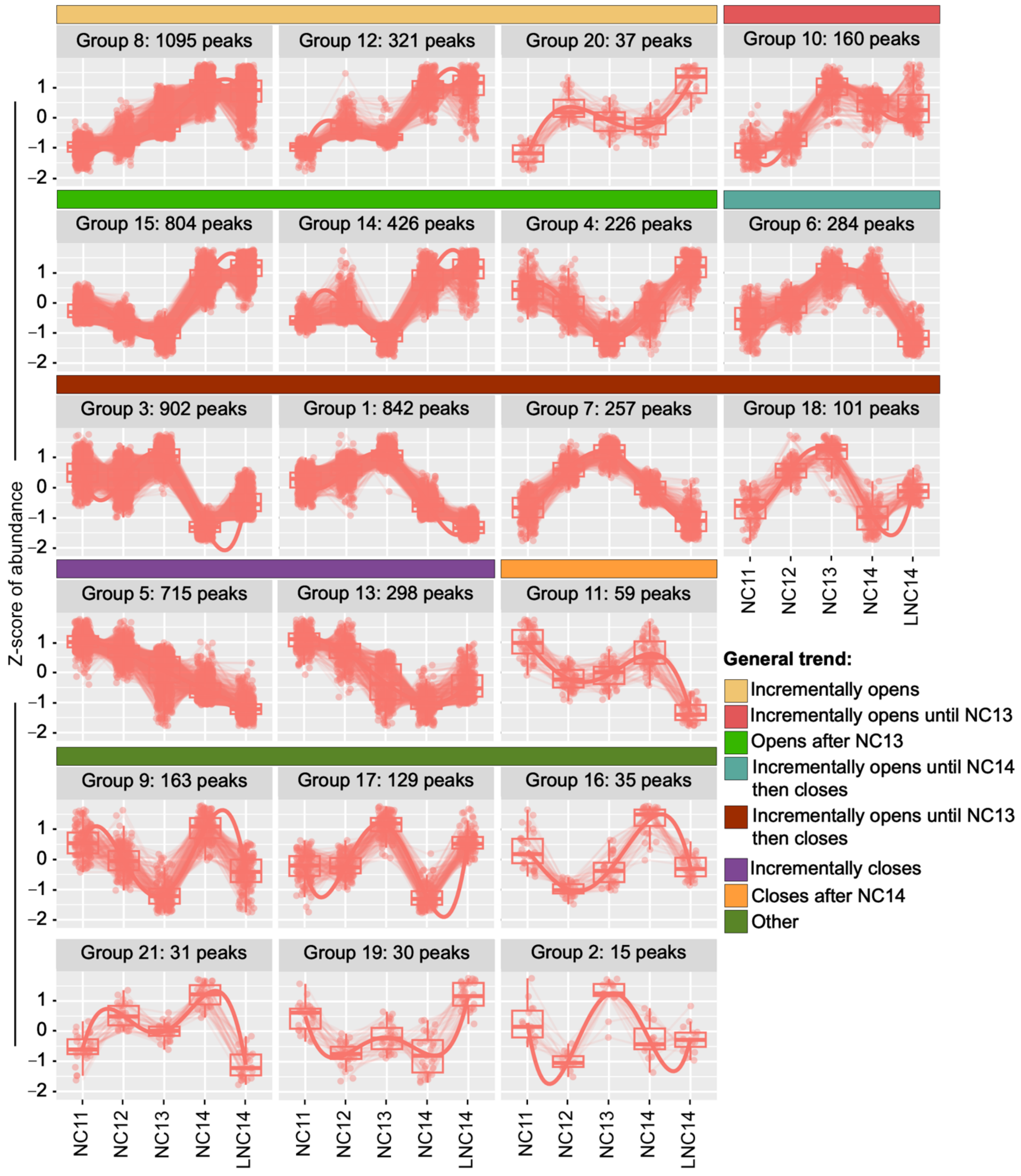


**S2 Figure, relating to Figure 2: Dynamic ATAC-seq peak groups.** Dynamic peak groups were generated using DEG reports (3) (R package version 1.38.5). Y-axis, Z-score of abundance. X-axis, developmental stage. Colored bars mark manually established super-groups with similar features described as follows: Incrementally opens (yellow bar; Groups 8, 12, 20); Incrementally opens until NC13 (red bar; Group 10); Opens after NC13 (green; Groups 15, 14, 4); Incrementally opens until NC14, then closes (blue; Group 6); Incrementally opens until NC13, then closes (red-brown; Groups 3, 1, 7, 18); Incrementally closes (violet; Groups 5, 13); Closes after NC14 (orange; Group 11); Other (olive; Groups 9, 17, 16, 21, 19, 15).

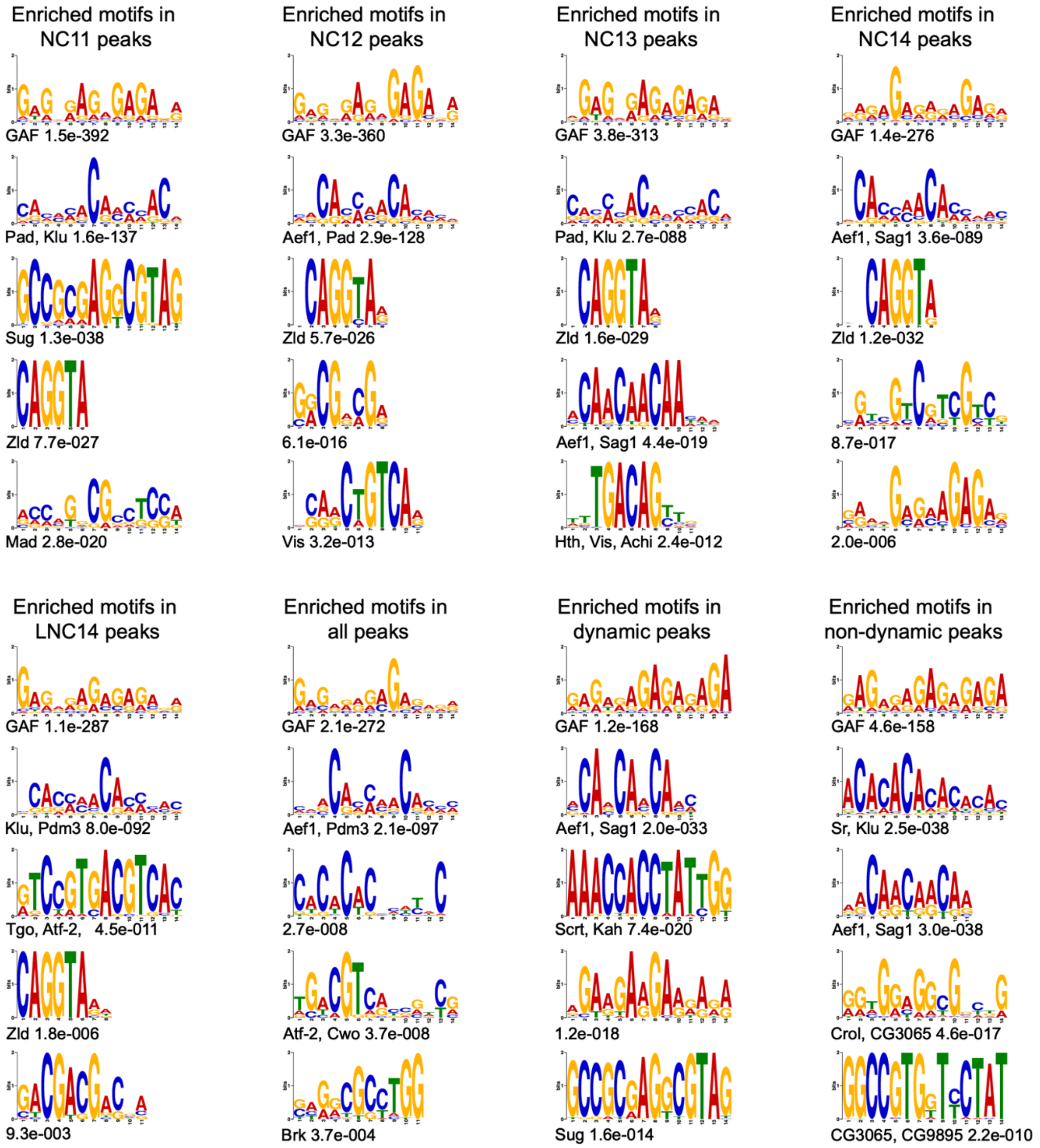


**S3 Figure, relating to Figure 2: Enriched motifs in ATAC-seq peaks.** Enriched motifs identified by MEME for 8 sets of peaks: NC11 peaks (n = 5013), NC12 peaks (n = 5926), NC13 peaks (n = 7586), NC14 peaks (n = 16147), Late NC14 peaks (n = 11628), all peaks (n = 32157), dynamic peaks (n = 6930), and non-dynamic peaks (right column; n = 25227). The top five motifs discovered by MEME are reported for each group. Motifs are shown as PWM logos with significance (E-value). DNA binding proteins of *Drosophila melanogaster* that bind to these or similar motifs as identified by MEME are indicated. We note that although not identified by MEME, CLAMP has been shown to bind GA-rich motifs and Cg has been shown to bind (CA)n like the motifs above (4-7).

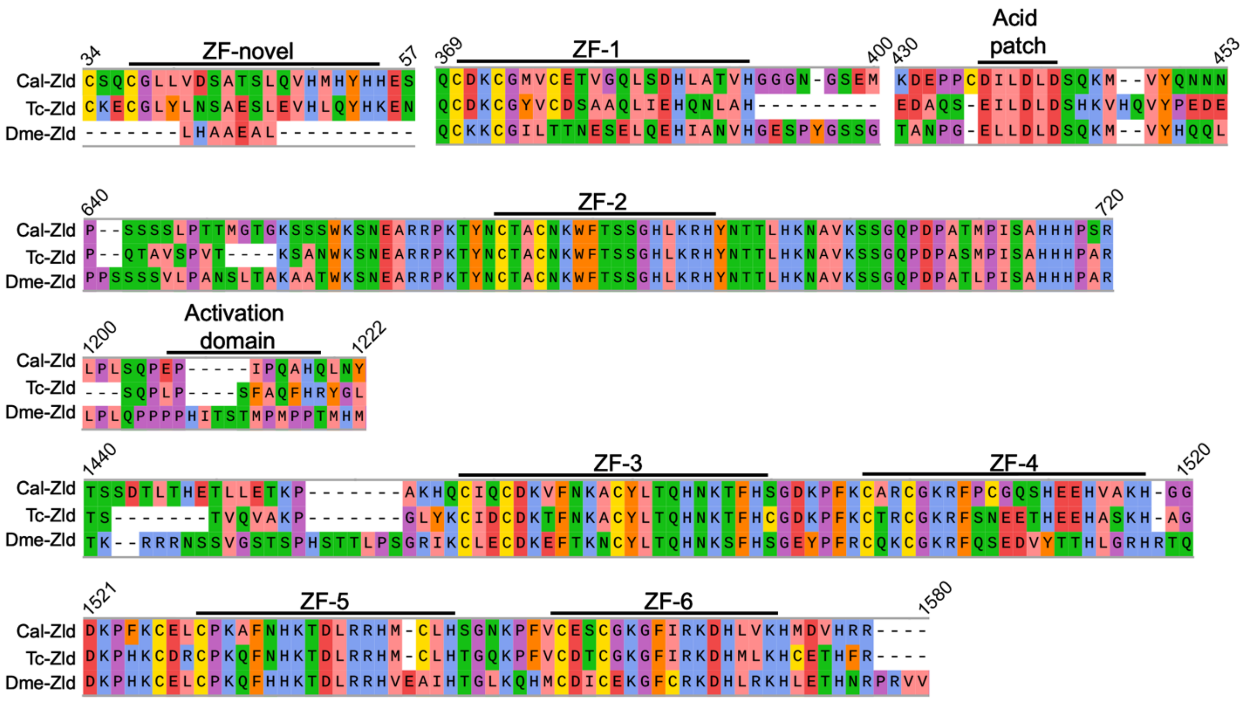


**S4 Figure: Protein alignment of conserved Zelda domains.** Cal-Zld sequence is shown above Zld sequences of *Tribolium castaneum* (Tc-Zld, XP_001812268.1) and *Drosophila melanogaster* (Dme-Zld, NP_608356.1). Alignment was performed using MAFFT (8) (v7) with the default settings. Amino acid color indicates physio-chemical property (based on Zappo Color Scheme). Important protein domains are indicated (9).

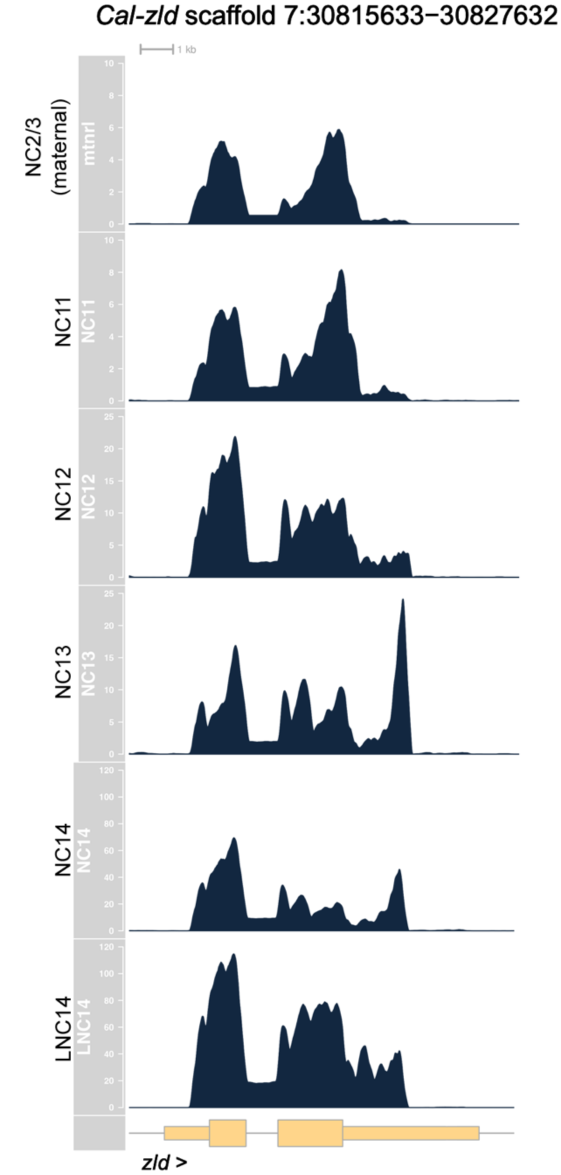


**S5 Figure: Time-course RNA-seq data of *Cal-zld* at 6 developmental stages.** Gene model in pale yellow. RNA-seq data CPM. Y-axis for NC2/3 (maternal, ~45-minute old embryos) and NC11: 0-10 CPM. Y-axis for NC12 and NC13: 0-25 CPM. Y-axis for NC14 and Late NC14 (LCN14): 0-120 CPM.

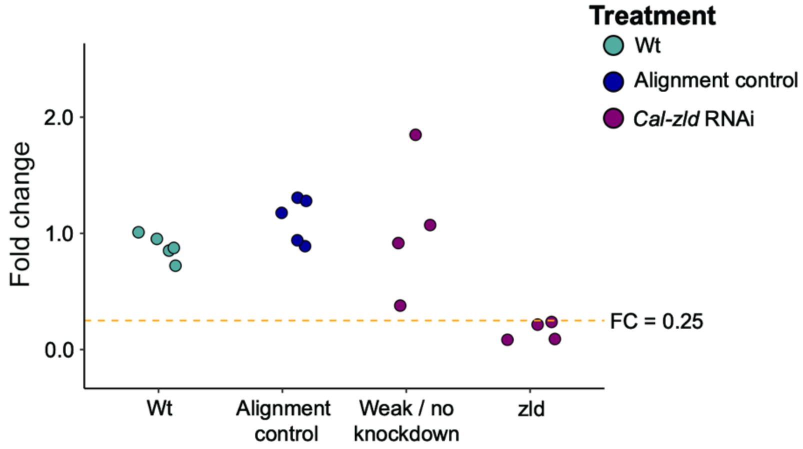


**S6 Figure**: **Identification of *Cal-zld* RNAi embryos with strong knockdowns.** Plot of *Cal-zld* fold change for individual embryos. Each embryo is represented by a circle. Embryos are grouped along the x-axis as indicated, spacing within a group was done arbitrarily for visualization. Treatment group of each embryo is indicated by color. Fold change (FC) for each embryo was calculated by dividing the embryo's CPM of *Cal-zld* by the average of wild-type and alignment control *Cal-zld* CPMs. Fold change was calculated using embryos from a single female (same activation batch). If controls from the same batch were lost, fold change was calculated using the average *Cal-zld* CPM of all wild-type and alignment control embryos in the data set. *Cal- zld* RNAi embryos with FC > 0.25 (weak / no knockdown of *Cal-zld*) were excluded from further analysis.

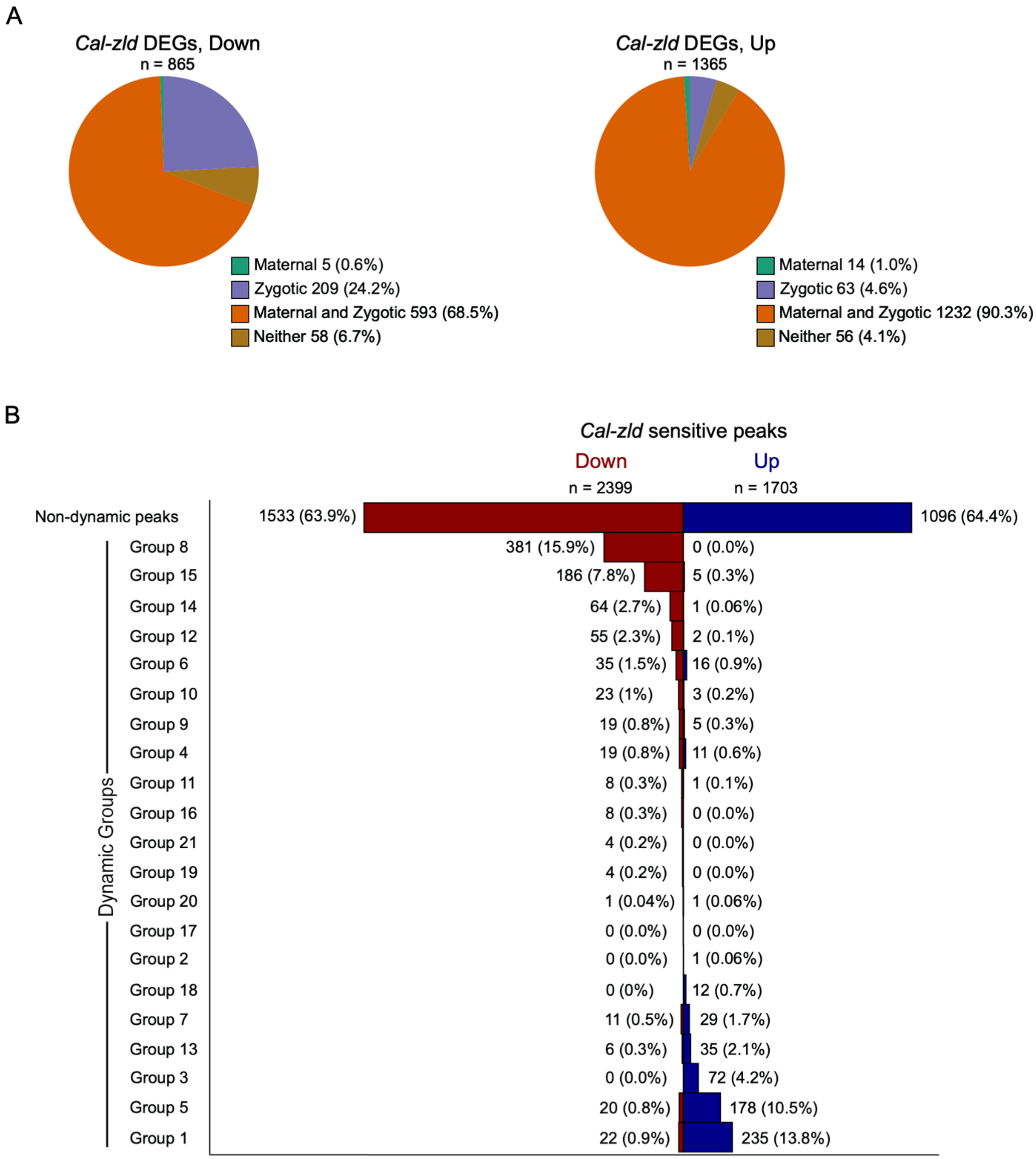


**S7 Figure, relating to Figure 3: Classifications of differentially expressed genes in *Cal-zld* RNAi embryos and *Cal-zld* sensitive ATAC-seq peaks.** A) Percentage of down-regulated (left) and up-regulated (right) genes in *Cal-zld* RNAi embryos classified as maternal, zygotic, maternal and zygotic or with little to no expression (neither) in wild type. Classifications were determined by expression analysis of genes at NC2/3 (~45-minute old embryos) or at syncytial blastoderm stages (NC11 through Late NC14, see *Materials and Methods*). B) *Cal-zld* sensitive ATAC-seq peaks in dynamic and non-dynamic groups. Peaks in different groups are ordered by *Cal-zld* sensitivity. Peaks that gained accessibility are shown in blue and peaks that lost accessibility are shown in red. Numbers of peaks and percentage relative to all down- or upregulated peaks are indicated.

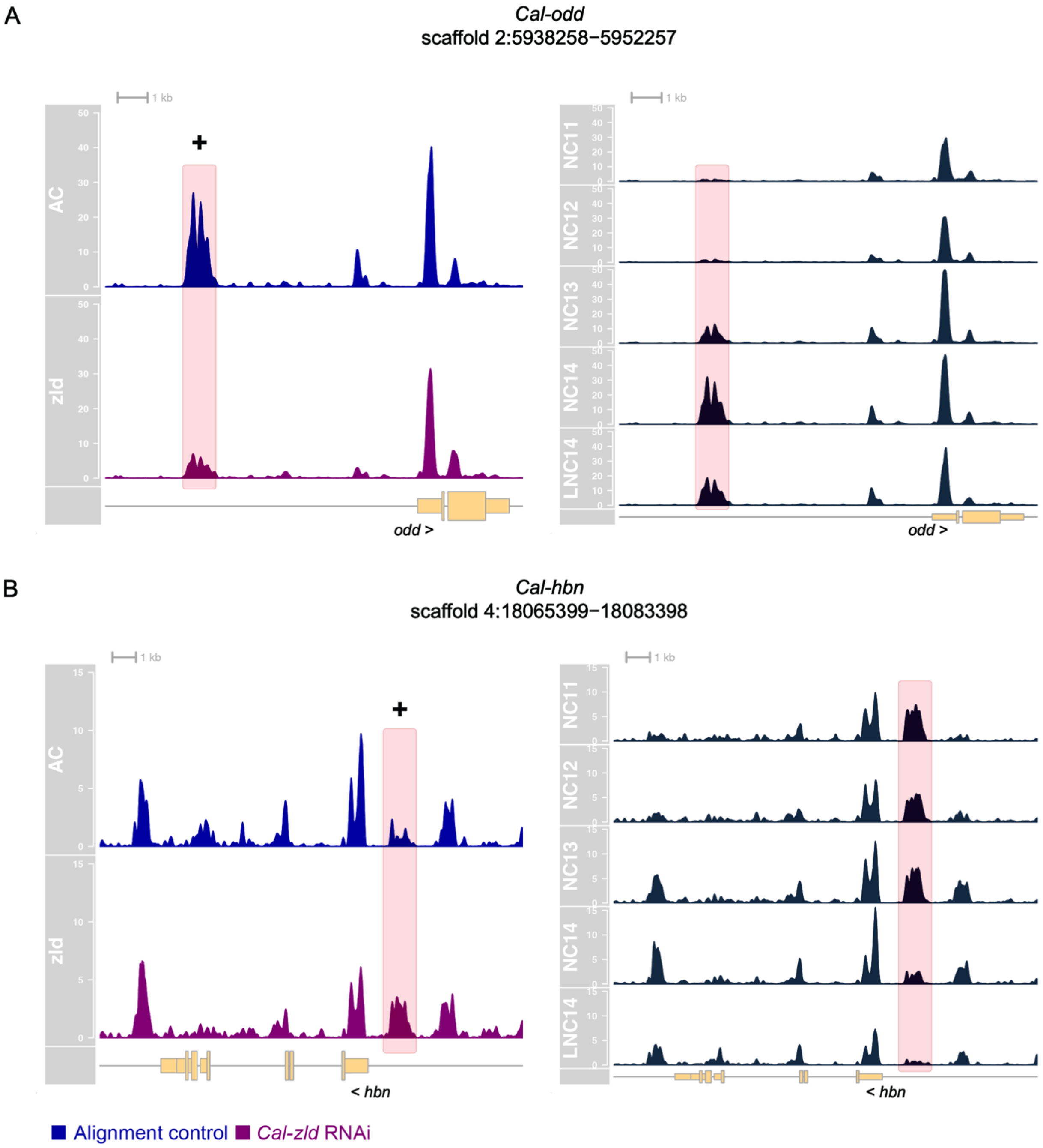


**S8 Figure, relating to Figure 3: Examples from the loci of two developmental genes that display *Cal-zld* sensitive peaks.** A) The locus of *Cal-odd* contains one chromatin accessibility peak whose accessibility is dependent on *Cal-zld* to open and contains a Zld motif (highlighted). On the right is the same view of the locus in wild-type embryos at NC11 through Late NC14. The peak is in dynamic group 8 and progressively gains accessibility as nuclear cycles progress. ATAC-seq tracks: 0-50 CPM. B) The locus of *Cal-hbn* contains one chromatin accessibility peak whose accessibility is dependent on *Cal-zld* to close and contains a Zld motif (highlighted). On the right is the same view of the locus in WT embryos at NC11 through Late NC14. The peak is in dynamic group 7 and progressively closes as nuclear cycles progress. ATAC-seq tracks: 0-15 CPM.

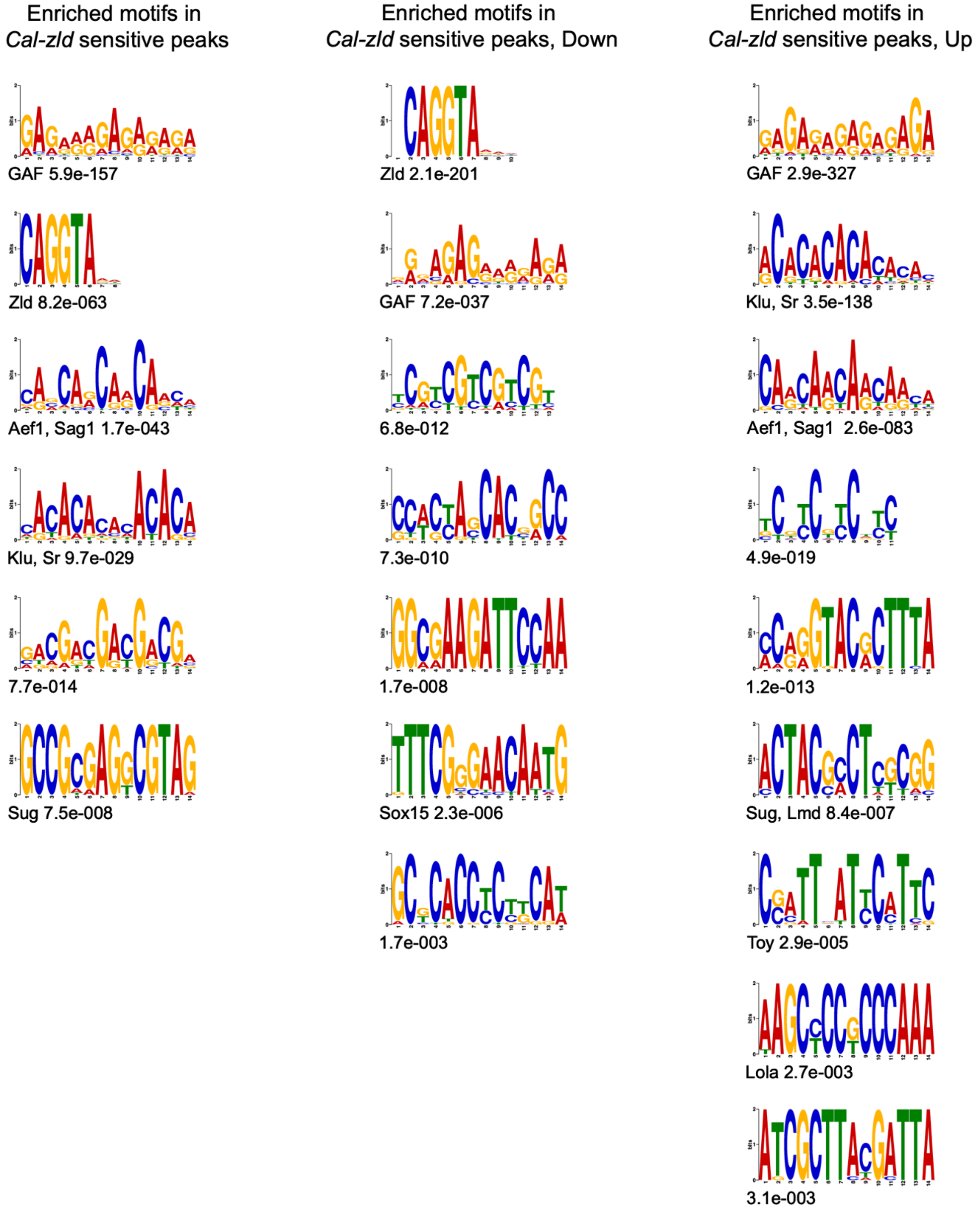


**S9 Figure, relating to Figure 3: Enriched motifs in *Cal-zld* sensitive peaks.** Enriched motifs identified by the MEME suite in all *Cal-zld* sensitive peaks (n = 4102, left), peaks whose accessibility decreases in *Cal-zld* RNAi embryos (n = 2399, center) and peaks whose accessibility increases in *Cal-zld* RNAi embryos (n = 1703, right). Motifs shown as a PWM logo with significance (E-value). DNA binding proteins of *Drosophila melanogaster* that bind to these or similar motifs as identified by MEME are indicated. We note that although not identified by

MEME, CLAMP has been shown to bind GA-rich motifs and Cg has been shown to bind (CA)n like the motifs above (4-7).

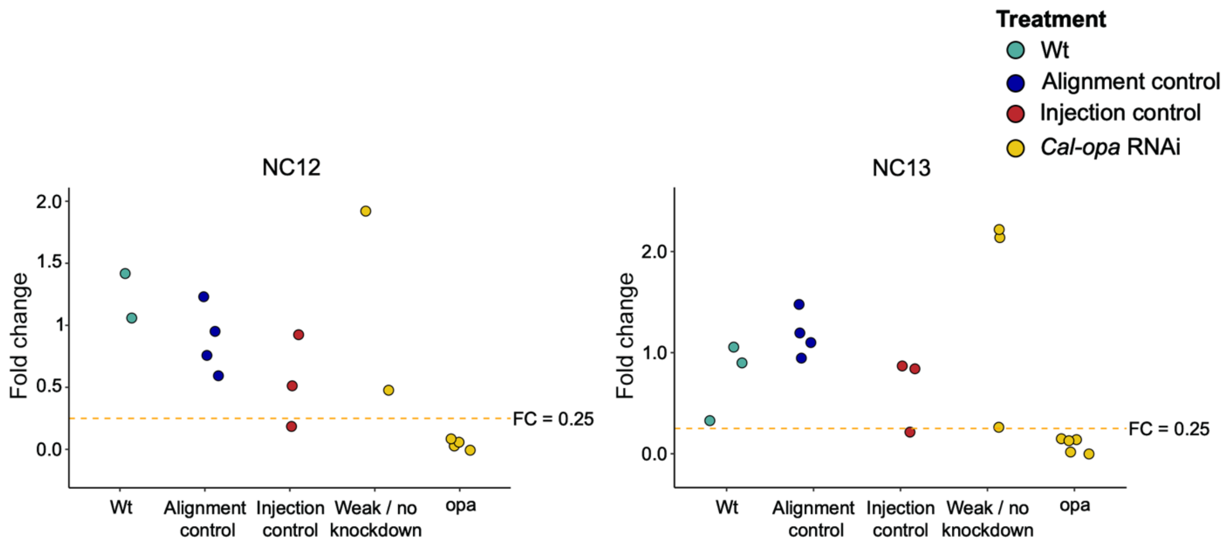


**S10 Figure: Identification of *Cal-opa* RNAi embryos with strong knockdowns.** Plots of *Cal-opa* fold change (FC) for individual embryos at NC12 and NC13. Each embryo is represented by a circle. Embryos are grouped along the x-axis as indicated, spacing within a group was done arbitrarily for visualization. Treatment group of each embryo is indicated by color. FC for each embryo was calculated by dividing the embryo's CPM of *Cal-opa* by the average of wild-type and alignment control *Cal-opa* CPMs. FC was calculated using embryos from a single female (same activation batch). *Cal-opa* RNAi embryos with FC > 0.25 (weak / no knockdown of *Cal-opa*) were excluded from further analysis. WT, wild type.

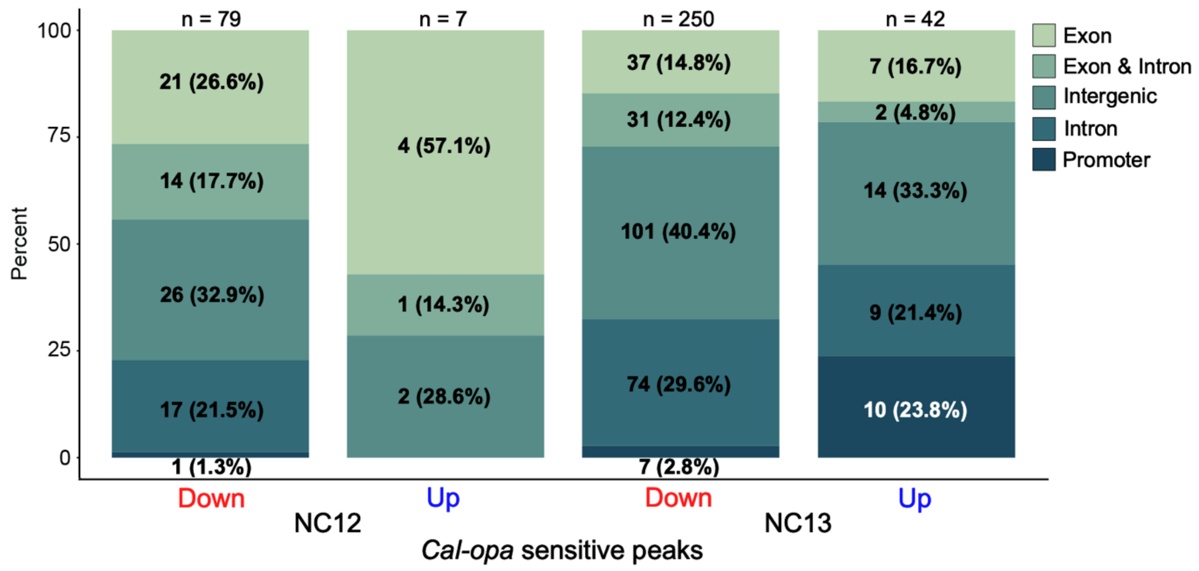


**S11 Figure, relating to Figure 4: Genomic distribution of *Cal-opa* sensitive peaks at NC12 and NC13.**

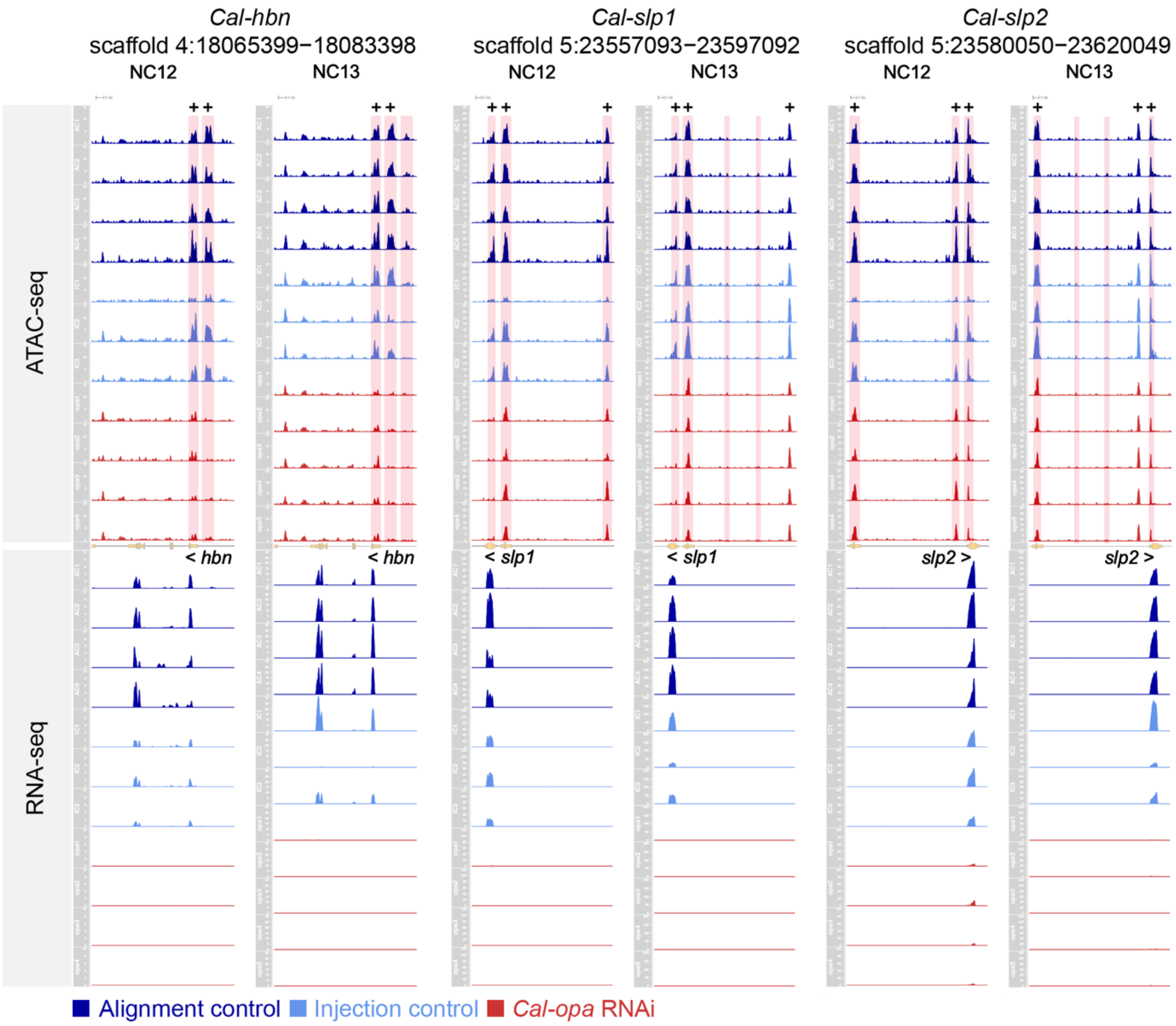


**S12 Figure, relating to Figures 6 and 7: ATAC-seq and RNA-seq tracks of *Cal-hbn*, *Cal-slp1* and *Cal-slp2* plotted for individual replicates.** Alignment controls (blue), injection controls (light blue), and *Cal-opa* RNAi embryos (red) are shown at NC12 and NC13. Top tracks: CPM of ATAC-seq data. Middle track: gene model in pale yellow. Bottom tracks: CPM of RNA-seq data. Peaks with significant changes in accessibility highlighted in pink. Peaks containing Opa motifs are marked with a plus sign. The y-axis is the same for all tracks of the same data type at each locus. *Cal-hbn* ATAC-seq at NC12 and NC13: 0-15 CPM. *Cal-hbn* RNA-seq at NC12: 0-10 CPM. *Cal-hbn* RNA-seq at NC13: 0-30 CPM. *Cal-slp1* ATAC-seq at NC12 and NC13: 0-30 CPM. *Cal-slp1* RNA-seq at NC12: 0-120 CPM. *Cal-slp1* RNA-seq at NC13: 0-200 CPM. *Cal-slp2* ATAC-seq at NC12 and NC13: 0-30 CPM. *Cal-slp2* RNA-seq at NC12: 0-100 CPM. *Cal-slp2* RNA-seq at NC13: 0-120 CPM.

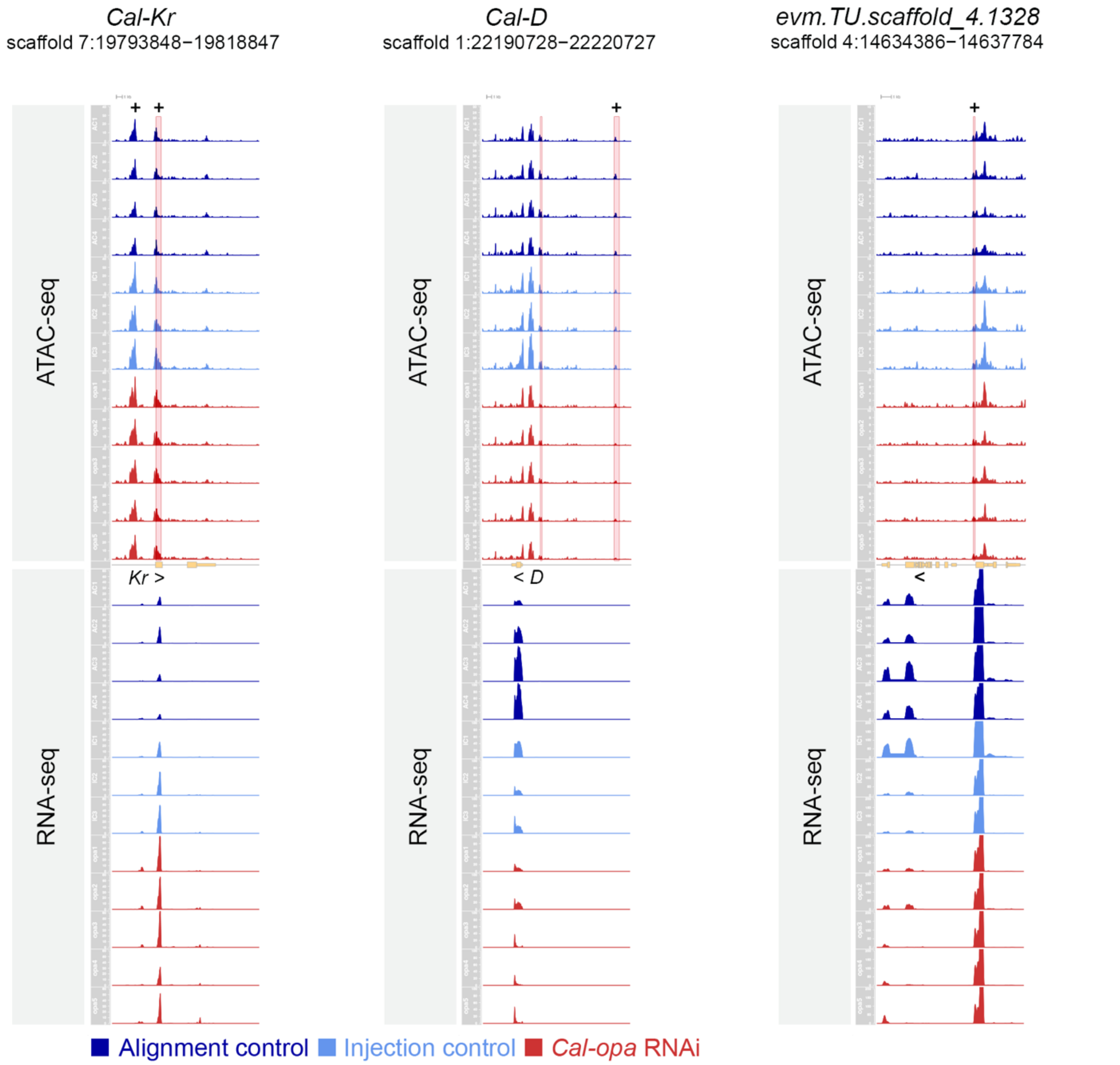


**S13 Figure relating to Figures 6 and 7: ATAC-seq and RNA-seq tracks of *Cal-Kr*, *Cal-D* and *evm.TU.scaffold_4.1328* plotted for individual replicates.** Alignment controls (blue), injection controls (light blue), and *Cal-opa* RNAi embryos (red) at NC13. Top tracks: CPM of ATAC-seq data. Middle track gene model in pale yellow. Bottom tracks: CPM of RNA-seq data. Peaks with significant changes in accessibility highlighted in pink. Peaks containing Opa motifs are marked with a plus sign. The y-axis is the same for all tracks of the same data type at each locus. *Cal-Kr* ATAC-seq: 0-40 CPM. *Cal-Kr* RNA-seq: 0-60 CPM. *Cal-D* ATAC-seq: 0-30 CPM. *Cal-D* RNA-seq: 0-100 CPM. *evm.TU.scaffold_4.1328* ATAC-seq: 0-10 CPM. *evm.TU.scaffold_4.1328* RNA-seq: 0-200 CPM.

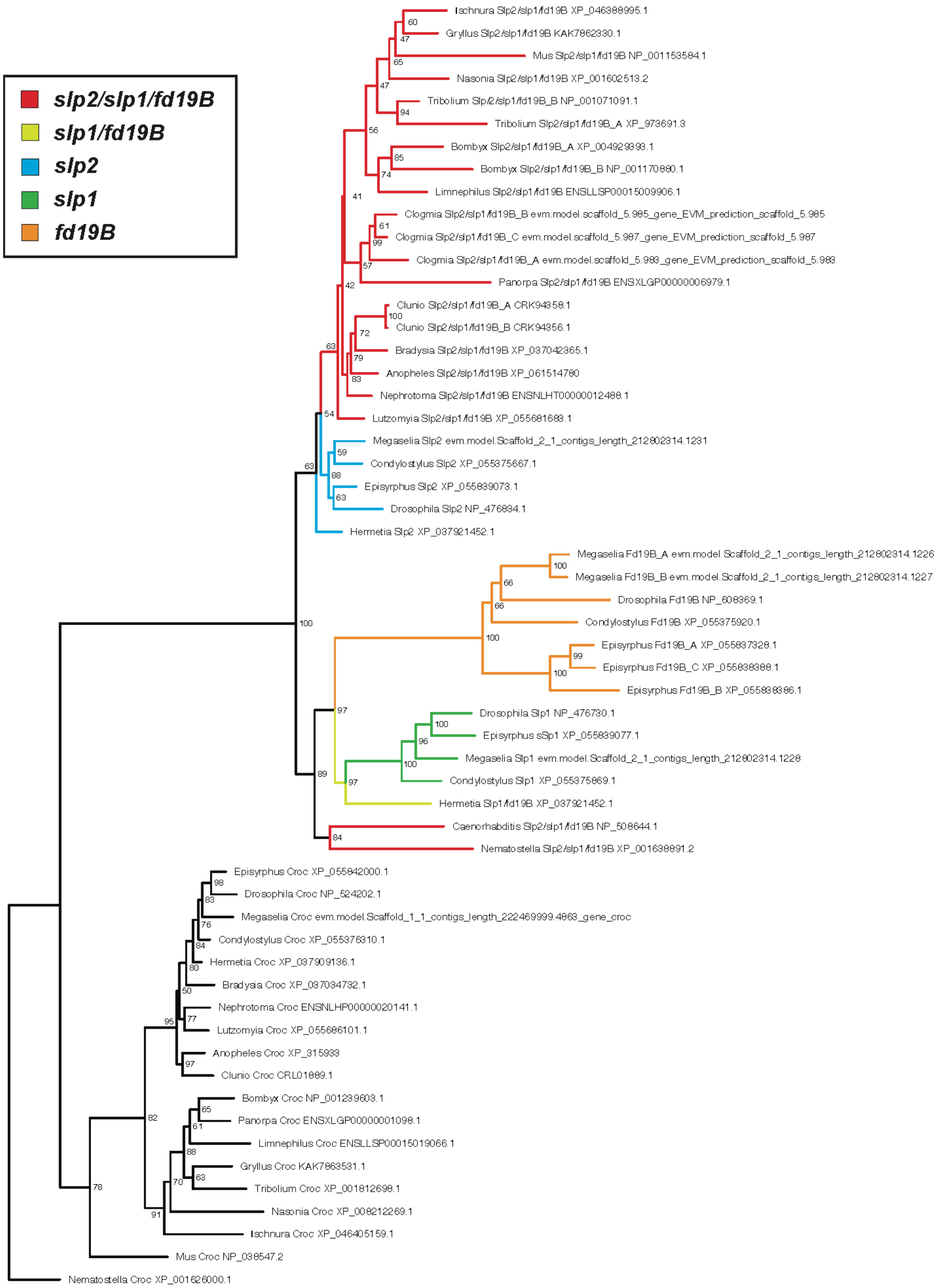


**S14 Figure: Maximum-likelihood tree of Sloppy-paired homologs.** Reciprocal protein BLAST was performed using Slp1 (NP_476730.1), Slp2 (NP_476834.1), Fd19B (NP_608369.1), and Crocodile (Croc; outgroup; NP_524202.1) as queries. Homologous protein sequences in *Nematostella vectensis*, *Mus musculus*, *Caenorhapditis elegans*, *Ischnura elegans*,

*Gryllus longicernus*, *Nasonia vitripennis*, *Tribolium castaneum*, *Bombyx mori*, *Clunio marinus*, *Anopheles gambiae*, *Lutzomyia longipalpis*, *Bradysia coprophila*, *Hermetia illucens*, *Condylostylus longicornis*, and *Episyrphus balteatus* were identified by reciprocal Protein BLAST (https://blast.ncbi.nlm.nih.gov/Blast.cgi) (1). Homologous sequences in *Limnephilus lunatus* (10), *Panorpa germanica* (11), and *Nephrotoma appendiculata* (12) were identified by reciprocal BLAST inGeneious Prime 2024.0.7 (https://www.geneious.com/), using respective transcript and protein fasta files available at ensemble-Darwin Tree of life (https://projects.ensembl.org/darwin-tree-of-life/). E value cut-off threshold was set to 0.05 and maximum number of target sequences set to 100. Identified sequences (for accession numbers see tree) were aligned using MAFFT v.7.526 with the L-INS-i strategy (https://mafft.cbrc.jp/alignment/software/) (8). Maximum likelihood trees were constructed using IQ-TREE v.2.3.6. (13). The best molecular substitution model for each partition under the Bayesian information criterion was selected by partition merging strategy with ModelFinder (14). Maximum likelihood trees were constructed under the selected substitution models (VT+F+I+G4), with branch support values estimated by the ultrafast bootstrap approximation (13) using 1000 replicates. A majority-rule consensus tree was then generated based on the bootstrap results. The consensus tree was visualized on FigTree v.1.4.4 (http://tree.bio.ed.ac.uk/software/figtree/) (15). Nematostella Croc sequence was manually chosen as root.

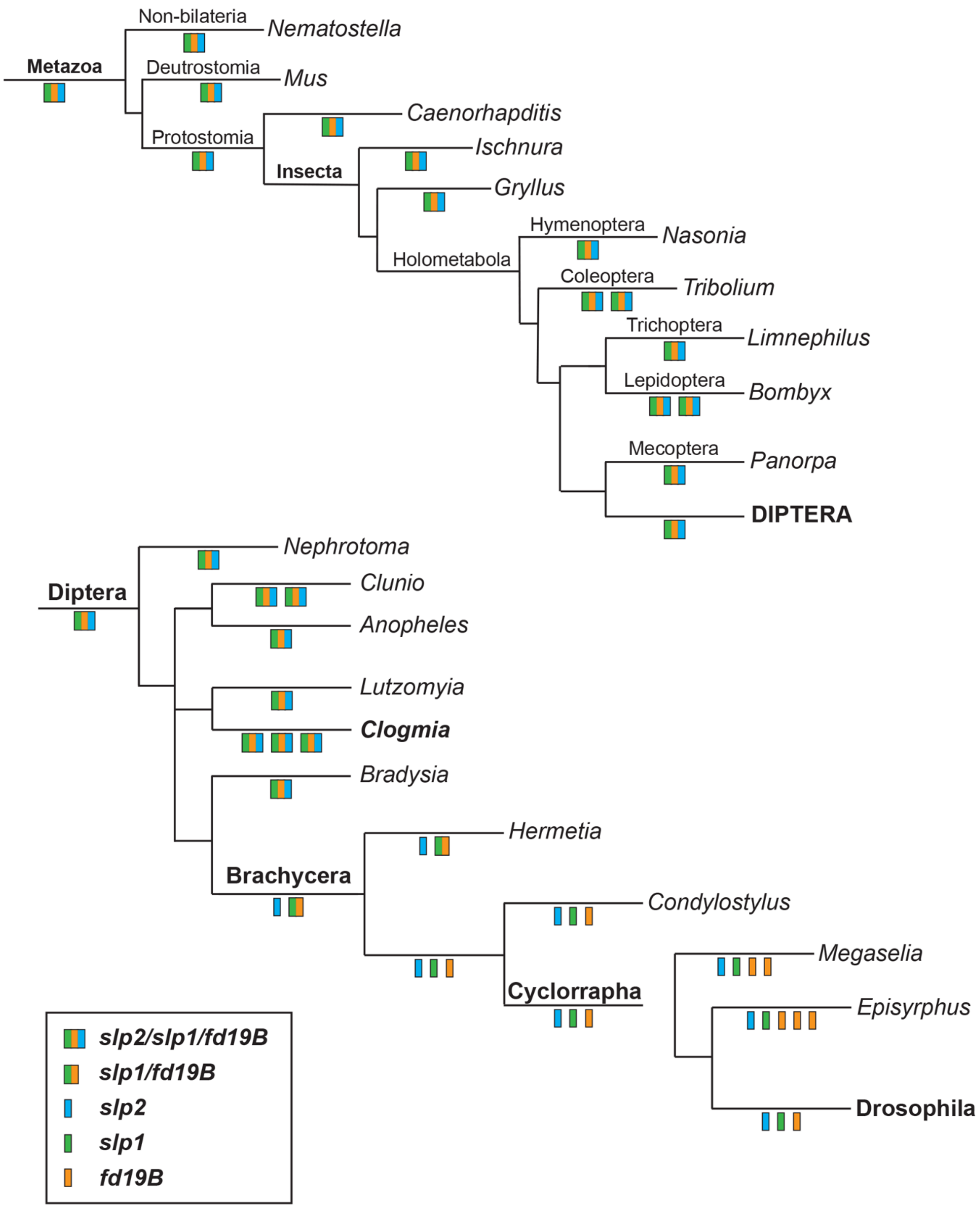


**S15 Figure: The evolution of *sloppy-paired*. Homologs of** *sloppy-paired* are mapped onto simplified trees of Metazoa (top) and Diptera (bottom). Orthologs of *slp1* (green), *Fd19B* (orange), *slp2* (blue), and the inferred *slp1/fd19B/slp2* and *Slp1/fd19B* progenitors are color coded.

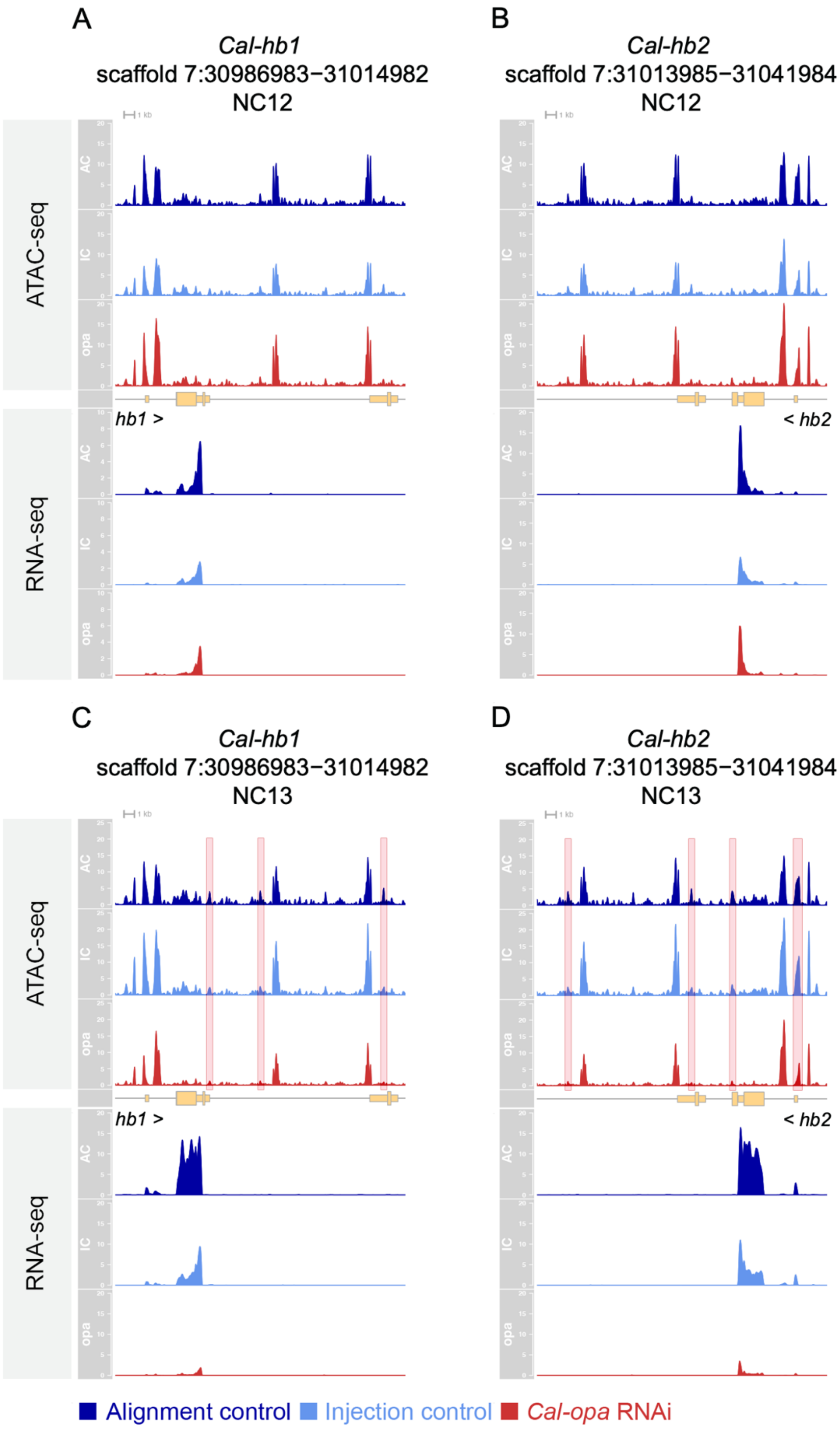


**S16 Figure: ATAC-seq and RNA-seq tracks of *Cal-hb1* and *Cal-hb2* at NC12 and NC13.** Top tracks: CPM of ATAC-seq data. Middle track gene model in pale yellow. Bottom tracks: CPM of

RNA-seq data. Peaks with significant changes in accessibility highlighted in pink. Peaks containing Opa motifs are marked with a plus sign. The y-axis is the same for all tracks of the same data type at each locus. A) *Cal-hb1*at NC12 ATAC-seq 0-20 CPM. *Cal-hb1* at NC12 RNA-seq 0-10 CPM. B) *Cal-hb2* at NC12 ATAC-seq 0-20 CPM. *Cal-hb2* at NC12 RNA-seq 0-20 CPM. C) *Cal-hb1*at NC13 ATAC-seq 0-25 CPM. *Cal-hb1* at NC13 RNA-seq 0-20 CPM. D) *Cal-hb2* at NC13 ATAC-seq 0-25 CPM. *Cal-hb2* at NC13 RNA-seq 0-20 CPM. Blue: Alignment control. Light blue: Injection control. Red: *Cal*-opa RNAi.

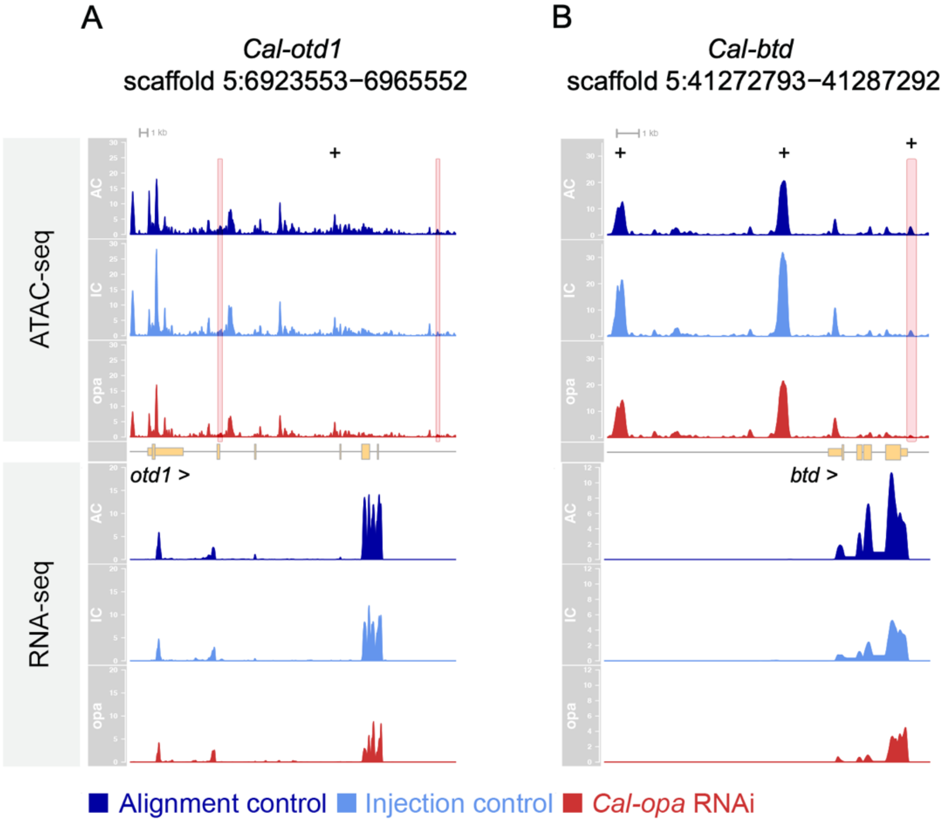


**S17 Figure: ATAC-seq and RNA-seq tracks of *Cal-otd1* and *Cal-btd* at NC13.** Top tracks: CPM of ATAC-seq data. Middle track gene model in pale yellow. Bottom tracks: CPM of RNA-seq data. Peaks with significant changes in accessibility highlighted in pink. Peaks containing Opa motifs are marked with a plus sign. The y-axis is the same for all tracks of the same data type at each locus. *Cal-otd1* ATAC-seq 0-30 CPM. *Cal-otd1* RNA-seq 0-20 CPM. *Cal-btd* ATAC-seq 0-35 CPM. *Cal-btd* RNA-seq 0-12 CPM. Blue: Alignment control. Light blue: Injection control. Red: *Cal*-opa RNAi.

**Movies and Appendix**

**Available at** https://schmidtottlab.uchicago.edu/data/

**S1 Movie: Time-lapse movie from ~NC11 to gastrulation.** Embryos from left to right; 1) *Cal-zld* RNAi 2) *Cal-zld* RNAi, 3) *Cal-zld* RNAi, 4) alignment control, and 5) *Cal-zld* RNAi. Transmitted light imaging was performed on a Leica DM5000B at 1-minute intervals. Imaging was stopped and needed to be restarted twice, this accrues at the ~30 second and ~65 second marks. Scale bar 100 μM.

**S2 Movie: Loss of Drosophila melanogaster *zelda* impairs cellularization and cytoplasmic clearing during zygotic genome activation**. This movie is a composite of two independently recorded movies aligned to coincide with the beginning of nuclear cycle 14, both of Cry2-zelda embryos. Cry2-zelda embryos express a chimeric Zld protein fused to the light-inducible Cry2 module, allowing *zelda* loss of function upon exposure to blue light (16). The top embryo was imaged at a permissive wavelength (zld+, 594 nm) and the bottom one was imaged at the restrictive wavelength (zld-, 488 nm) on a confocal microscope with a 30-second frame rate. The timestamp indicates minutes relative to the beginning of NC14. Note that while the zld+ embryo develops clear cortical cytoplasm and cellularizes during the ~1h period of NC14, the zld- embryo retains dark cytoplasm at the basal surface of the nuclei and fails to undergo cellularization, prior to loss of epithelial integrity (~ +45 minutes, lower right). Scale bar, 50 μm.

**S1 Appendix**: Dovetail/Cantata Bio Hifiasm and HiRise Scaffolding Reports

**S2 Appendix**: Summary *Cal-opa* RNAi analysis at NC12. 1) DESeq2 analysis of *Cal-opa* RNAi verses alignment control RNA-seq data. 2) DESeq2 analysis of *Cal-opa* RNAi verses alignment control ATAC-seq data. 3) Differentially expressed genes and ATAC-seq peaks within their loci. 4) Differentially expressed transcription factors and ATAC-seq peaks within their loci. 5) Differentially expressed transcription factors containing Opa motif positive ATAC-seq peaks (putative direct Cal-Opa targets). 6) Differentially expressed transcription factors with *Cal-opa* sensitive ATAC-seq peaks. 7) Differentially expressed transcription factors with *Cal-opa* sensitive ATAC-seq peaks containing an Opa motif. 8) DESeq2 analysis of *Cal-opa* RNAi verses injection control RNA-seq data. 9) DESeq2 analysis of Cal-*opa* RNAi verses injection control ATAC-seq data.

**S3 Appendix**: Summary *Cal-opa* RNAi analysis at NC13. 1) DESeq2 analysis of *Cal-opa* RNAi verses alignment control RNA-seq data. 2) DESeq2 analysis of *Cal-opa* RNAi verses alignment control ATAC-seq data. 3) Differentially expressed genes and ATAC-seq peaks within their loci. 4) Differentially expressed transcription factors and ATAC-seq peaks within their loci. 5) Differentially expressed transcription factors containing Opa motif positive ATAC-seq peaks (putative direct Cal-Opa targets). 6) Differentially expressed transcription factors with *Cal-opa* sensitive ATAC-seq peaks. 7) Differentially expressed transcription factors with *Cal-opa* sensitive ATAC-seq peaks containing an Opa motif. 8) DESeq2 analysis of *Cal-opa* RNAi verses injection control RNA-seq data. 9) DESeq2 analysis of Cal-*opa* RNAi verses injection control ATAC-seq data.
